# Thousandfold Expansion Microscopy

**DOI:** 10.64898/2026.05.31.729018

**Authors:** Helena Hu, Donatus Krah, Antonios Ntolkeras, Sushovan Chanda, Alina Heimbrodt, Milton Mondal, Jonas Altendorf, Bowen Jing, Bonnie Berger, Ali H. Shaib, Silvio O. Rizzoli, Edward S. Boyden

**Affiliations:** Department of Biological Engineering, MIT, Cambridge, MA 02139; Department of Neuro- and Sensory Physiology, University Medical Center Göttingen, Göttingen, Germany; McGovern Institute, MIT, Cambridge, MA 02139; Departments of Brain and Cognitive Sciences, Media Arts and Sciences, MIT, Cambridge, MA 02139; Koch Institute, MIT, Cambridge, MA 02139; Mindspan Institute, Cambridge, MA 02139; K Lisa Yang Center for Bionics, Yang Tan Collective, MIT, Cambridge, MA 02139; Computer Science and AI Lab, Massachusetts Institute of Technology; Dept. of Mathematics, Massachusetts Institute of Technology

## Abstract

Biological macromolecules, such as proteins, are made of concatenated building blocks. We hypothesized that individual protein residues could be imaged by anchoring their side chains to a swellable polymer, cleaving backbone amide bonds, and expanding residues away from each other to a degree that enables them to be visualized separately. We introduce thousandfold expansion microscopy (1000ExM), a four-network interpenetrating hydrogel architecture that enables successive expansion from ∼18-fold to >1000-fold (one billion-fold in volume). Protein and peptide structures are maintained across these expansion factors, as verified by analyses of proteins with known structures (nanobodies, GFP) and a well-studied peptide (mCLING). Computational analysis indicates that 1000ExM resolves adjacent amino acid residues, thereby achieving sub-nanometer precision on conventional light microscopes. We anticipate that 1000ExM will find wide utility in protein visualization and identification, potentially even in intact cells and tissues.

## Introduction

Biological macromolecules, such as proteins, are made of repeated building blocks in a linear chain, which then attain a tertiary structure through folding. The ability to image individual building blocks of macromolecules, ideally in a way that accurately reports on the structure of the macromolecule being imaged, would be of great use in the analysis of proteins and peptides, both natural and engineered. Peptides, in particular, are too small to image with electron microscopy [1,2], and the limited number of different information channels (or “colors”) that can be employed in electron microscopy limits the number of building block species (amino acids, in the case of peptides) that can be identified. Ideally, one would be able to see the building blocks of a biological macromolecule on ordinary light microscopy hardware, in a strategy extensible to multiple colors, and in a fashion that could in principle be adapted to the mapping of proteins within cells and tissues.

Expansion microscopy (ExM) improves resolution by physically expanding biological specimens, increasing distances between molecules so that features below the diffraction limit become resolvable on ordinary microscopy hardware [6]. Recently, ExM has been used to expand proteolyzed fragments of proteins away from each other by ∼10×. This expansion factor, combined with a super-resolution fluorescence fluctuation analysis that builds on the super-resolution radial fluctuations (SRRF) [3] and super-resolution optical fluctuation imaging (SOFI) [4] concepts, optimized for expansion microscopy, enables the shapes of proteins to be visualized, in what we termed one-step nanoscale expansion microscopy (ONE) [5]. In ONE, super-resolution requires thousands of frames to generate a single two-dimensional image, and three-dimensional reconstructions are assembled from many such images. Ultimately, ONE reveals protein shape, but does not resolve individual amino acids. We therefore asked whether individual amino acids could instead be resolved directly on conventional microscopes, without super-resolution, by increasing the expansion factor further. This requires physical magnification to bring formerly intramolecular distances above the diffraction limit. The diffraction limit is approximately 200 to 300 nanometers (nm) (lateral; the axial resolution of a confocal microscope would be slightly worse, 300 to 500 nm), whereas the distance between adjacent alpha carbon (Cα) atoms in a protein backbone, where side chains are attached, is ∼0.38 nm. After 1000× expansion, this distance would become ∼380 nm, within the resolvable range of conventional microscopy. Thus, sufficient expansion could make individual molecular building blocks directly observable. Of course, such a procedure would have to be isotropic, expanding biological samples evenly in all dimensions, in order to preserve submolecular positional information.

In iterative expansion microscopy (iExM [6–9]), a specimen is expanded in a swellable ionic gel, re-embedded in a neutral gel to stabilize it in the expanded state, and then a second swellable ionic gel is cast. The specimen is then expanded a second time, thus leading to larger expansion factors than the original gel (the original iterative protocols performed 4.5× expansion twice, with a neutral gel-stabilizing step in between, resulting in ∼16-22× total expansion). Here, we identify an interpenetrating hydrogel architecture, enabled by the use of a specific gel formulation, as well as an iterative strategy that omitted the neutral re-embedding step, allowing expansion to proceed through multiple rounds, ultimately reaching 1000-fold linear expansion (at each step along the way, presenting qualitatively new useful outcomes). A prior attempt to recursively expand a specimen by iterating without neutral re-embedding resulted in a small expansion factor (∼10-fold expansion after 4 rounds of expansion), due to gel composition differences [10]. We here show, using nanobodies bearing fluorophores, the peptide mCLING [11], and the protein GFP, that 1000ExM was able to reveal the structures of such proteins on ordinary confocal microscopes, with resolution sufficient to pinpoint the successive amino acids on a linear chain in 3D. This residue-level spatial information suggests that proteins may be identifiable directly from their coordinate patterns. To test this, we simulated protein identification across the human proteome (23,391 canonical proteins), incorporating incomplete residue readout and experimentally measured expansion-induced distortion. Under these conditions, most proteins could be uniquely identified from their spatial amino acid coordinate patterns alone; 1000ExM may thus find broad utility in the visualization of proteins and peptides, potentially in intact cells and tissues.

## Results

### Development of 1000ExM

We developed a four-network interpenetrating polymer network (IPN) architecture, and a series of steps to assemble it, that enable ∼1000× linear expansion. In summary, achieving such a large expansion factor resulted from (1) the choice of swellable ionic network itself, and (2) how to preserve the expanded state in a high-integrity way during sequential casting. The latter is traditionally done with a neutral reembedding gel that keeps an expanded gel in the open state, for a subsequent round of polymerization and expansion [12].

Neutral re-embedding has been classically considered helpful in stabilizing expanded gels [13]. We optimized a gel made with sodium acrylate monomers and dimethylacrylamide crosslinkers (SA/DMAA), decreasing KPS concentration in blank gels from 0.5% to 0.05% w/v in 0.05% decrements. Each gel was polymerized to completion overnight to mature the crosslinking degree. Using the optimal SA/DMAA formulation, we asked whether we could cast a new charged gel directly within an expanded charged gel, eliminating the traditional neutral re-embedding step. Prior attempts to do so with classical ExM formulations did not yield expansion factors greater than those achievable in a single round of expansion [10]. Interestingly, when an expanded gel was exposed to the monomer solution for casting a second network, it did not fully collapse upon gelation. In a representative gel, it contracted from ∼18× to ∼10× and reached a stable, partially swollen state. A similar pattern held across all four rounds: introducing each new monomer solution caused the existing gel to relax to roughly half of its prior expansion factor, rather than collapsing back toward baseline, leaving it swollen enough for the next charged network to form within it. In short, sequential casting of four optimized SA/DMAA networks produced compounding linear expansion across rounds, on the order of 18x to 100x to 500x to 1500x in a representative case, without intermediate neutral re-embedding gels (**Fig. 1A, Table S18**). These values varied somewhat from gel to gel, as expected from natural chemical process variability across four sequential casting steps, and are intended to illustrate the overall trend rather than a fixed trajectory. The protocol used in the majority of the datasets here described is contained in **Supplementary Note 2**; an updated protocol that reduces variability and offers more predictable performance (and, often, higher expansion factors) is contained in **Supplementary Note 1** (and was used to generate **Table S18**). Upon exposure to phosphate-buffered saline (PBS), gels stabilized at a median linear expansion of approximately 1000× (**Fig. 1B**). Thus, our work demonstrates a charged ionic gel that resists electrostatic collapse in a charged monomer solution of defined composition, so that charged-within-charged network formation can occur without requiring an intervening neutral scaffold, thereby permitting large expansion factors. Our updated, and improved, workflow involves having each casting round include two incubations in activated monomer solution before polymerization, to facilitate monomer loading in large gels, and produced four-network gels with a median linear expansion of approximately 1900×, reaching as high as 2534× in some gels. The variability in the procedure is easily compensated for by measuring the expansion factor of individual gels, and suggests that the architecture described here is not intrinsically limited to 1000×, but could support potentially much larger expansion factors, providing a platform for future development in the field (**Fig. S31**).

**Figure 1.**
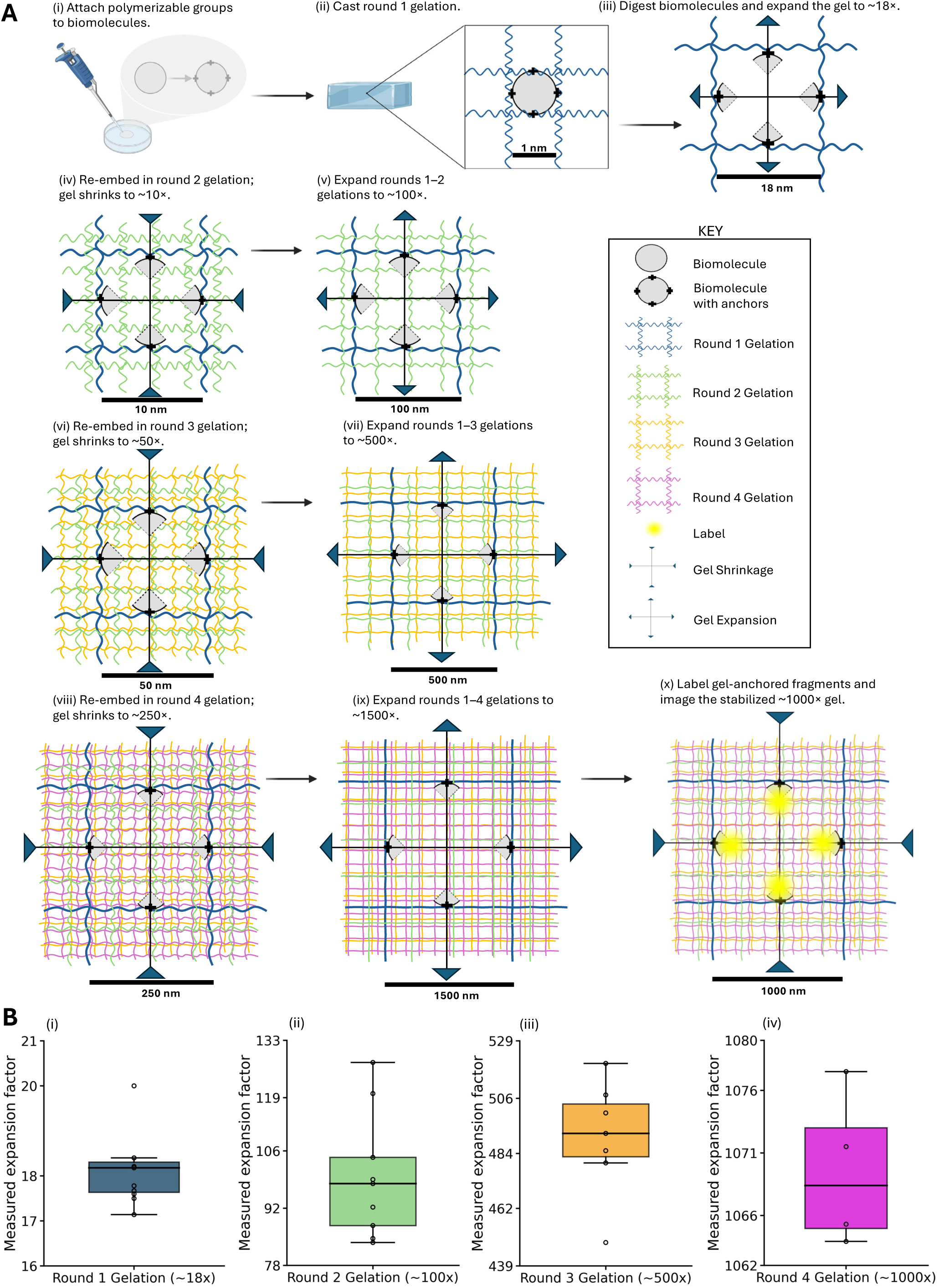
Design of a four-network interpenetrating polymer network (IPN) architecture enabling ∼1000× linear expansion via recursive ionic-in-ionic casting. (A) Schematic of four-round ionic-in-ionic casting sequence used to construct the interpenetrating charged hydrogel architecture. The recursive casting process yields ∼1000× linear (∼10⁹ volumetric) expansion and enables separation of biomolecular building blocks anchored within the gel matrix following macromolecule bond cleavage, including (but not limited to) individual amino acid residues within proteins. (i) Biomolecules are functionalized with polymer-incorporable groups to enable covalent anchoring to the hydrogel network. For proteins, primary amines can be modified with acryloyl-X, SE (AcX) to install polymerizable acrylate groups. (ii) Modified biomolecules are embedded in a swellable charged hydrogel network generated by reacting sodium acrylate (25.60% w/v), N,N-dimethylacrylamide (DMAA, 44.20% v/v), N,N,N′,N′-tetramethylethylenediamine (TEMED, 0.037% v/v), and potassium persulfate (KPS, 0.145% w/v). (iii) Anchored biomolecules are fragmented by cleavage between anchoring points. For proteins, proteinase K (8 U/mL) generates anchored peptide fragments. The gel is expanded in water to ∼18× linear expansion, separating covalently anchored biomolecular fragments (e.g., peptides, individual amino acid residues, or other subunits), depending on the anchoring and fragmentation strategy. (iv) The fully expanded ∼18× gel is infiltrated with a charged monomer solution and polymerized, yielding a composite gel that contracts to ∼10× linear expansion. (v) The composite gel is expanded in water to ∼100× linear expansion. (vi) The fully expanded ∼100× gel is infiltrated with a charged monomer solution and polymerized, yielding a composite gel that contracts to ∼50× linear expansion. (vii) The composite gel is expanded in water to ∼500× linear expansion. (viii) The fully expanded ∼500× gel is infiltrated with a charged monomer solution and polymerized, yielding a composite gel that contracts to ∼250× linear expansion. (ix) The composite gel is expanded in water to ∼1500× linear expansion. (x) Gel-anchored biomolecular fragments are labeled through reactive functional groups generated during fragmentation. For proteins, primary amines exposed by proteinase K digestion can be labeled with NHS-ester fluorophores. The gel is then equilibrated in 1× DPBS, contracting to ∼1000× linear expansion, which stabilizes the sample and enables stable imaging over extended durations. (B) Physical expansion factors measured after full expansion at each round of gelation, shown as boxplots: (i) ∼18× round 1 gelation (n = 11 separate gelations), (ii) ∼100× round 2 gelation (n = 9 separate gelations), (iii) ∼500× round 3 gelation (n = 7 separate gelations), (iv) ∼1000× round 4 gelation (n = 4 separate gelations). Boxplots in this figure: the center line denotes the median, the box spans the interquartile range (IQR), whiskers extend to the farthest data point within 1.5×IQR of each box edge, and individual measurements are shown as open circles. Expansion proceeded through four sequential rounds of gelation and expansion, with a subset of the gels that made it through round N, going through round N+1 (simply for convenience); for a set of gels that were all advanced in parallel, using a slightly modified protocol (two monomer incubations per round), see **Supp Fig. 31**.

### Proportional scaling of molecular distances across expansion rounds

To test whether molecular distances scale with physical expansion, we measured fluorophore–fluorophore separations using two independent nanorulers across multiple expansion factors (**Fig. 2**). We used a site-directed conjugated nanobody with known crystal structure [14] carrying two fluorophores embedded in gel-anchoring tags (called ExM cassettes) consisting of 4 lysines at the N- and C-termini of the nanobody [15]. The fluorophores are separated by ∼4 nm **(Fig. 2A,B**; additional representative images for each expansion factor are shown in **Supplementary Fig. 2)**. This provides a test case at the single-digit nanometer scale. At low expansion factors, this separation is below the diffraction limit (4 nm × 18× ≈ 72 nm), so early rounds were imaged using one-step nanoscale expansion (ONE) microscopy [5]. Higher expansion factors were also imaged using conventional confocal microscopy. Representative nanobodies for each expansion factor are shown in **Fig. 2A**. Across all expansion factors, measured fluorophore–fluorophore distances scaled proportionally with the physical expansion factor, defined as the ratio of post-expansion to pre-expansion linear dimensions (**Fig. 1B**). Distances were normalized using the condition-level approximate expansion factors of ∼18×, ∼100×, ∼500×, and ∼1000×. This normalization yielded an inferred biological distance of ∼4 nm, consistent with the expected labeling distance (**Fig. 2B**; all raw data and summary statistics in **Table S17**). Corrected distance distributions were consistent across gels and closely matched distributions modeled by projecting, on a 2D plane, two fluorophore positions found at a distance corresponding to the approximate positions of the fluorophores on the known structure of the nanobodies [14] (**Fig. 2B**, right), indicating that nanoscale distances are preserved through all rounds of expansion. Variability in the measurements, implying values such as those beyond the expected size of the nanobodies, is most likely due to measurements in which only one fluorophore on a specific nanobody survives the gelation procedure, implying that its nearest neighbor will be found on an adjacent nanobody. Because the exact positions of the 4-lysine ExM cassettes flanking the fluorophore attachment sites and lysine anchoring positions on the nanobody are not precisely known (albeit they are known to be placed at the two poles of the longest axis of the nanobody [5]), the computational model does not use an atomic structure or residue-level coordinates. The nanobody is approximated as a rigid ∼4 nm globular object with two fluorophores attached via ∼1 nm flexible lysine linkers. Fluorophore positions were sampled by linker flexibility and random three-dimensional orientations of the linkers. Distances were measured from two-dimensional projections of these configurations, generating simulated distance distributions directly comparable to the experimentally measured, expansion-corrected data.

**Figure 2.**
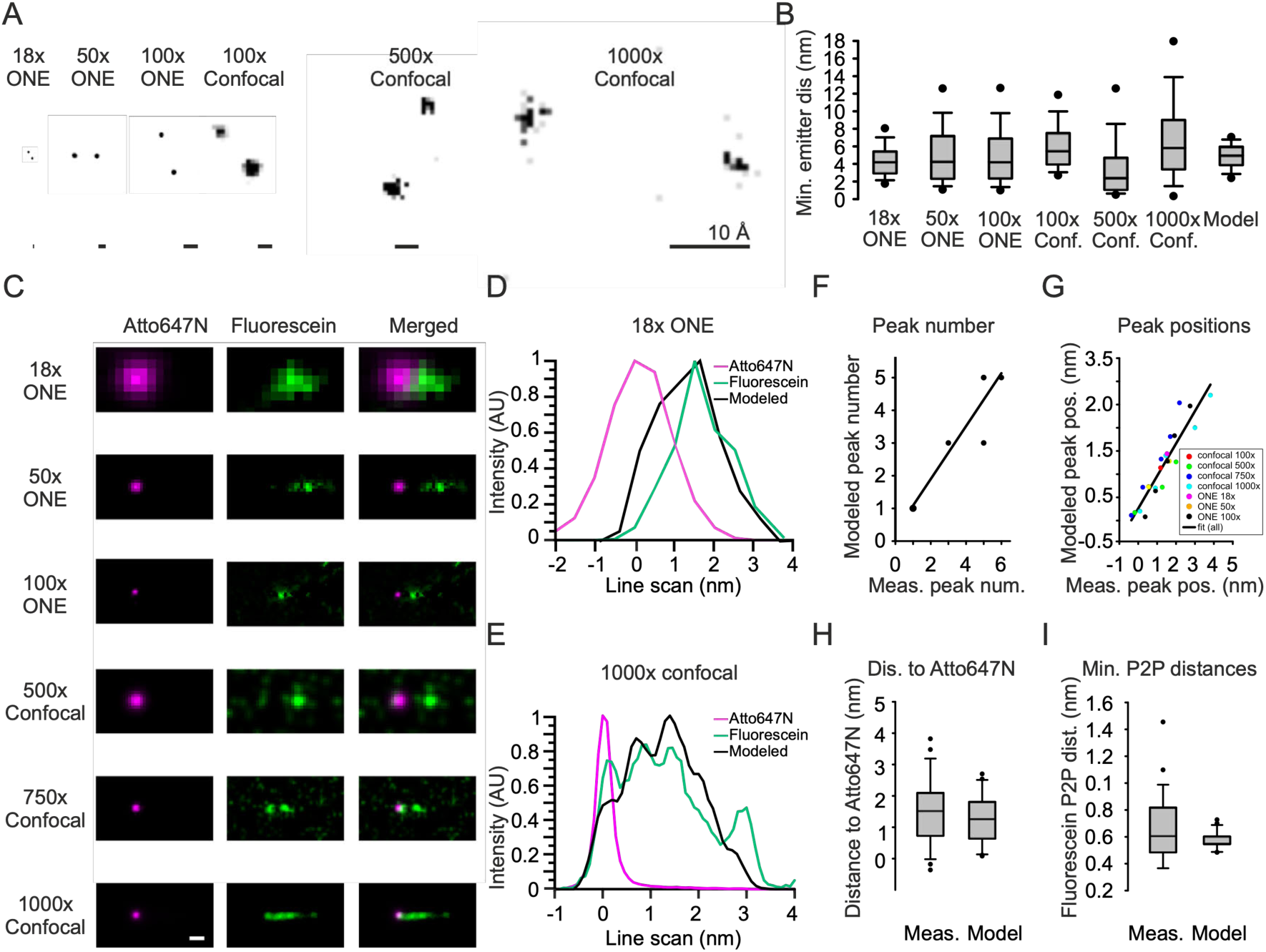
Large expansion factor gels resolve fluorescence of adjacent amino acids in a peptide. (A) Exemplary (in terms of brightness; n = 9395, 956, 892, 326824, 16974, 735 nanobodies from 18×/ONE, 50×/ONE, 100×/ONE, 100×/confocal, 500×/confocal, 1000×/confocal gels, respectively; representative datasets are shown, from n = 4 experimental replicates for each condition) images of nanobodies carrying two fluorophores, imaged using ONE microscopy [5,42,43] or confocal microscopy, with different expansion factors. Scale bar (in biological units, used throughout), 10 Å. (B) Distance between centroids of spots, for nanobody fluorophore pairs. Gels preserve the ∼4 nm distance (in biological units) between fluorophores, as expected from the literature, and predicted by a computational model (right). n numbers as in panel a, for the microscopy measurements, and 1000 simulated nanobodies, for the model; full statistics in **Table S17**). Middle line, median; box boundaries, the 25th and 75th percentiles; whiskers, the 10th and 90th percentiles; dots, the 5th and 95th percentiles. (C) mCLING peptides carrying an Atto647N fluorophore (magenta) were fragmented and expanded and were labelled with fluorescein (green) on all amino acids bound to the gel with cleaved-and-exposed N termini. Larger expansion factors reveal increasingly more details in the green signal. The images show averages (after aligning all centroids of the Atto647 dyes) of all mCLING peptides identified in the respective gels (n = 1210 from 7 gel replicates, 924 from 3 gel replicates, 1099 from 2 gel replicates, 1017 from 2 gel replicates, 232 from 3 gel replicates, 18900 from 5 gel replicates, for the 18×/ONE, 50×/ONE, 100×/ONE, 500×/confocal, 750×/confocal, 1000×/confocal gels, respectively). Scale bar, 10 Å. (D) Fluorescein peak positions compared to mCLING amino acid positions predicted by molecular dynamics (MD). The modelled signal distribution (black; shown is a simulated line cut through the middle of the average of 10,000 model images; the modeled Atto647N is not shown) closely follows that measured for 18× gels (ONE microscopy, n = 1210 initial images; shown is a line cut through the middle of the averaged image of the corresponding panels of c, aligned to the maximum intensity center of the Atto647N spot). (E) As in d, but for 1000× gels (confocal microscopy, n = 18900 initial images). (F-G) Comparisons between the numbers of fluorescein peaks (f) and peak positions (g), for modeled vs. measured distributions, combined across gel types. Each dot represents a given peak averaged across all molecules in all gels; bigger expansion factor gels have more peaks visible; color code is shown in the key in g. A high similarity is observed (R2 = 0.86 for the number of spots, p = 0.0028, n = 7 gel types; R2 = 0.85 for the spot positions, p<0.0001, n = 22 spot positions). (H) Distances between the first fluorescein peak and the Atto647N signal (C-terminus) vs. the modelled ones (p = 0.216, Mann-Whitney Ranksum test). The box plots show the median (middle line), the lower and upper quartiles (box), the minimum and maximum numbers that are not outliers (the whiskers), and the outliers (calculated using the interquartile range). (I) As in h, but for the distances between adjacent fluorescein peaks (p = 0.383, Mann-Whitney Ranksum test).

Having established accuracy in the single-digit nanometer regime, we next tested whether this scaling extends to sub-nanometer separations. We next examined how gel expansion scales distances between anchored amino acids using mCLING (**Fig. 2C–I**), a peptide that serves as a well-defined molecular ruler. The chemical structure, anchoring strategy, proteolytic cleavage, and post-expansion labeling chemistry are shown in **Supplementary Fig. 4**; in summary, the C-terminus of mCLING exhibits an Atto647N dye, and the lysines along the chain both present anchoring opportunities as well as fluorescein anchor sites, post-cleavage. With increasing expansion factor, additional fluorescein intensity peaks became separable with respect to one another and with respect to the Atto647N reference (**Fig. 2C**; additional representative images for each expansion factor are shown in **Supplementary Fig. 3**). For comparison across expansion states, peptide images (fluorescein channel) were aligned to a linear structure, placed to the right of the Atto647N centroid, as shown in the figure, and were averaged, yielding spatial signal distributions for each condition. Fluorescein peak positions measured from these averages matched positions predicted by MD ensemble model, both at low expansion (18×; **Fig. 2D**) and at high expansion (1000×; **Fig. 2E**). Across gels, the number of fluorescein peaks we could detect, and their relative positions, scaled with expansion and agreed with model predictions (**Fig. 2F,G**). Distances between the Atto647N signal and the first fluorescein peak, as well as distances between adjacent fluorescein peaks, were statistically indistinguishable from modeled values (**Fig. 2H,I**). Overall, this analysis indicates that a simple 2D averaging of signals is sufficient to reveal amino acid-level positions in mCLING images, without any specialized (or user-biased) analysis procedures.

### Evaluation of expansion isotropy at the single-molecule level

In the past, the isotropy of expansion microscopy has been evaluated primarily by registering pre- and post-expansion images of the same sample [6], but for small molecules as explored here, pre-expansion imaging would be impractical. To directly probe isotropy at the molecular scale, we assessed whether 1000× linear expansion preserves and linearly scales sub-nanometer inter-residue distances into a diffraction-resolvable regime. A molecular ground truth for comparison with experimental data was defined by generating an ensemble of 1001 mCLING peptide conformations using molecular dynamics (MD) simulation (*see Methods, mCLING molecular dynamics simulations*). This ensemble captures a range of physically accessible peptide conformations prior to chemical processing. Each molecule then undergoes anchoring, proteolytic digestion, and post-expansion labeling, with less than 100% efficiency at each step. We explicitly modeled the full chemical processing sequence (**Supplementary Fig. 4**), including pan-lysine NHS–acrylate anchoring to primary amines, covalent attachment of anchoring nitrogens to the hydrogel network, Proteinase K digestion with potential cleavage or retention of Atto647N, and post-expansion fluorescein labeling of remaining (and newly exposed) free primary amines. Because anchoring occurs at specific amine nitrogens that become covalently attached to the gel, the measured fluorophore positions after expansion correspond to the displaced coordinates of those anchoring nitrogen atoms (**Supplementary Figs. 4–6**).

For each of the 1001 conformations, we applied the defined anchoring, cleavage, and labeling scenarios to generate plausible labeling outcomes (**Supplementary Fig. 6**), resulting in a combined ensemble defined by peptide conformations and their corresponding labeling outcomes. A putative peptide was defined as containing one Atto647N dye accompanied by at least four fluorescein dyes. The mCLING peptide contains a central palmitoyl tail that interacts hydrophobically with the C-terminal Atto647N, resulting in a predominantly curved, C-shaped conformation in the model, which is the dominant conformation observed across the molecular dynamics ensemble. Accurate structural matching therefore would benefit from capturing the full arc of the molecule: with four or five fluorophores, we reasoned that the geometry captured would sufficiently contain any such curvature; with fewer labeling sites, partial sampling of the arc might permit under-constrained fitting.

MD-derived conformations were converted into modeled MD peptides by assigning fluorophores to the atomic coordinates of their anchoring nitrogen atoms. To quantify three-dimensional expansion distortion at the single-molecule level, experimentally derived dye coordinates from putative peptides were aligned to these modeled MD peptides. From 27,426 Atto647N localizations, 6,284 putative peptides containing one Atto647N dye and 4–5 fluorescein dyes were identified using nearest-neighbor criteria (*see Methods, Identification of putative peptides*). For each putative peptide, measured dye coordinates were aligned to each member of the full computationally defined ensemble of modeled peptides to identify the best-matching conformation, and then uniformly scaled to determine the scale factor that minimized root mean square deviation (RMSD), yielding a per-molecule expansion factor. The minimized RMSD provides an upper bound on expansion-induced distortion, as it reflects both distortion introduced during expansion and uncertainty in the modeled conformational ensemble, which may not fully capture the complete spectrum of physically accessible conformations. Molecule-to-molecule variation in best-fit MD matching was visualized by overlays of experimental dye coordinates and corresponding best-fit modeled MD peptides (**Fig. 3B**). Overlays stratified by RMSD below the interquartile range (IQR) (**Supplementary Fig. 15**), within the IQR (**Supplementary Fig. 16**), and above the IQR (**Supplementary Fig. 17**) show progressively reduced alignment: peptides below the IQR exhibit near-complete correspondence, with at most one displaced site; those within the IQR show typically ∼two displaced sites; and those above the IQR typically show misalignment at more than two sites.

**Figure 3.**
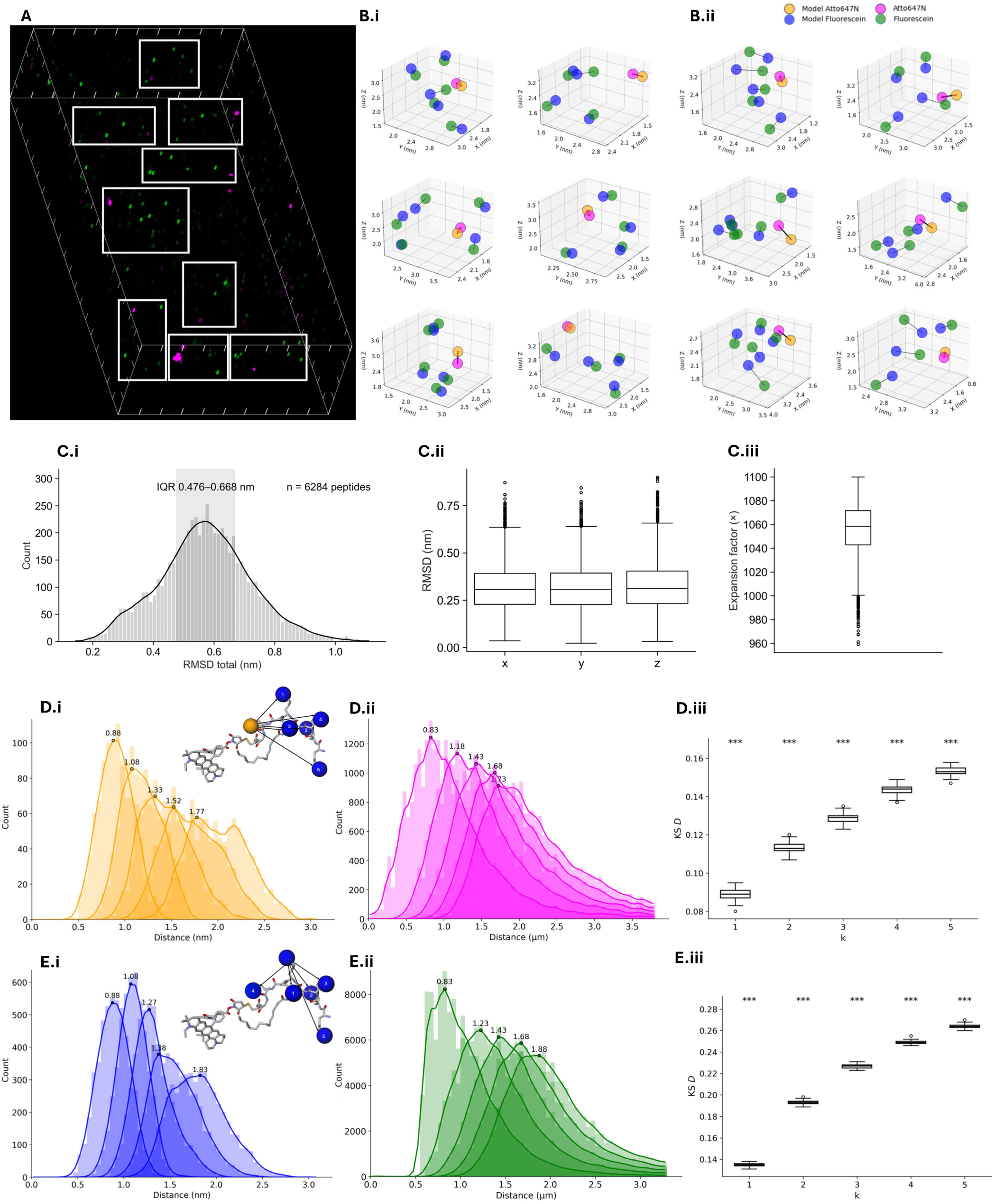
Thousand-fold expansion microscopy (1000ExM) enables three-dimensional multicolor residue-resolution imaging on standard light microscopes. (A) Three-dimensional rendering of a 10 × 20 µm field spanning the full z depth (4.4 µm) after ∼1000× linear expansion. Boxed regions indicate putative mCLING peptides, defined as clusters (hand-drawn, in this example) containing one Atto647N signal (magenta) and ≥4 proximal fluorescein signals (green). (B) Overlays comparing experimentally measured dye positions with molecular-dynamics (MD) predictions, for individual putative mCLING peptides. Experimental Atto647N and fluorescein positions are shown in magenta and green; model positions are shown in orange and blue. Lines connect matched centroids after optimal alignment of the experimental data with the best-match MD prediction, as computed by root mean square deviation (RMSD). (i) Exemplar overlays with RMSD below the interquartile range (IQR) of the peptide population. (ii) Representative overlay with RMSD within the IQR. (C) Quantitative comparison of experimental and MD-predicted geometries for the full population of putative mCLING peptides (n = 6,284 from 8 regions across 3 gels). (i) RMSD distribution computed after optimal alignment to an MD reference ensemble. Each peptide contains one Atto647N signal and ≥4 associated fluorescein signals identified using a k-nearest-neighbor (kNN; k = 1–5) definition (see Methods: Identification of putative mCLING peptides). (ii) RMSD along x, y, and z. Boxplots throughout Fig. 3: center line, median; box, IQR; whiskers, farthest data point within 1.5×IQR of each box edge; points, outliers. (iii) Single-molecule expansion factors estimated by fitting an isotropic scale factor that minimizes RMSD. (D) Atto647N–fluorescein distances. (i) kNN (1–5) distances from MD simulations of the mCLING peptide. (ii) Corresponding distances measured after ∼1000× expansion (n = 27,426 signals from 8 regions across 3 gels). (iii) Observed kNN (1–5) distance distributions were compared to randomized controls using the two-sample Kolmogorov–Smirnov test. Boxplots show Kolmogorov–Smirnov D statistics for each kNN rank; asterisks (***) indicate P < 0.001. (E) Fluorescein–fluorescein distances. (i) kNN (1–5) distances from MD simulations of the mCLING peptide. (ii) Corresponding distances measured after ∼1000× expansion (n = 125,395 signals from 8 regions across 3 gels). (iii) Observed kNN (1–5) distance distributions were compared to randomized controls using the two-sample Kolmogorov–Smirnov test. Boxplots show Kolmogorov–Smirnov D statistics for each kNN rank; asterisks (***) indicate P < 0.001. Distance distributions are shown as count histograms (bin width 0.05 nm for MD and 0.05 µm for expanded datasets). For expanded datasets, kNN ranks were pooled to define the display range, and the upper x-axis limit was set to the 95th percentile of the pooled distribution. Smoothed curves represent Gaussian-convolved counts; peak distances correspond to the bin center at the curve maximum. Cartoon legend (Di, Ei). Atoms are colored carbon (gray), nitrogen (purple), oxygen (red), and sulfur (yellow, small circle). Large spheres denote nitrogen atoms covalently linked to the hydrogel and displaced with network expansion. The orange sphere marks the nitrogen associated with the Atto647N label; blue spheres mark lysine ε-amine nitrogens.

To assess the robustness of this metric to the size of the MD reference ensemble, we evaluated how RMSD estimates varied with conformational sampling. Because each putative peptide is matched to the best-fit modeled MD peptide, limited sampling could bias RMSD values. We therefore repeated the matching using ensembles ranging from 10 to 1001 conformations. With 10–200 conformations, RMSD distributions were bimodal with a tail extending to ∼1.2 nm, consistent with incomplete sampling of accessible conformations. By 600 conformations, the distribution became unimodal (0.585 ± 0.154 nm, mean ± standard deviation used throughout), and increasing to 1001 conformations yielded 0.573 ± 0.151 nm (**Supplementary Figs. 13–14**). RMSD estimates stabilized beyond ∼600 conformations, indicating that 1001 conformations provided sufficient conformational coverage for stable matching.

Across all putative peptides (n = 6,284), each matched against the full 1001-conformation MD ensemble, RMSD values were sub-nanometer, with a mean of 0.573 ± 0.151 nm (**Fig. 3C.i**), and per-axis RMSD values of 0.317 ± 0.121 nm (x), 0.316 ± 0.120 nm (y), and 0.325 ± 0.126 nm (z) (**Fig. 3C.ii**). Per-axis RMSD values were comparable across x, y, and z, consistent with isotropic expansion at the single-molecule level. The optimal scale factor from this alignment and scaling procedure yielded a single-molecule expansion factor of 1055.92 ± 21.554 (**Fig. 3C.iii**). Distributions of RMSD, per-axis RMSD, and single-molecule expansion factors were consistent across 3 gels and 8 regions (**Supplementary Figs. 9–11**; pooled statistics in **Table S13**; gel- and region-stratified values in **Tables S14 and S15**), supporting reproducibility across samples and imaging regions.

To examine 1000ExM peptide expansion fidelity through a third kind of statistical analysis, we examined Euclidean k-nearest-neighbor (kNN, ranks 1–5) peak distances. The ensemble of modeled MD peptides of mCLING yielded kNN peak distances spanning 0.88–1.77 nm for Atto647N–fluorescein pairs and 0.88–1.83 nm for fluorescein–fluorescein pairs (**Fig. 3D.i, Fig. 3E.i**). After ∼1000× expansion, experimental kNN (1–5) peak distances spanned 0.83–1.73 µm and 0.83–1.88 µm, respectively, pooling data across eight regions from three gels (**Fig. 3D.ii, Fig. 3E.ii**). Distance distributions stratified by region and gel were consistent across independent gel runs (**Supplementary Fig. 19**), with pooled statistics in **Table S5** and gel-specific statistics in **Tables S6–S8**. As a measure of how likely such distributions could have arisen from random fluorophore placement, we randomized emitter coordinates within each 3D imaging volume while preserving emitter counts (*see Methods, Randomization of k-nearest-neighbor distances*). For both Atto647N–fluorescein and fluorescein–fluorescein pairs, two-sample Kolmogorov–Smirnov (KS) tests showed that measured distributions differed significantly from their randomized counterparts for kNN ranks 1–5 (p < 0.001 in all cases; **Fig. 3D.iii, Fig. 3E.iii; Supplementary Fig. 20**). The KS D statistic increased, going from kNN1 to kNN5, indicating that higher-order nearest-neighbor structures were progressively less consistent with random spatial organization. In other words: going from clusters of two dyes, up to four to five dyes, becomes increasingly unlikely, vs. random placement, consistent with structured molecular organization. Analyses performed separately for each gel yielded consistent results (**Supplementary Figs. 21–23**).

### GFP 3D reconstruction approaches amino acid resolution

We have two complementary strategies for achieving amino acid resolution: 1000-fold expansion followed by confocal imaging, or 100-fold expansion followed by ONE microscopy imaging. 1000-fold expansion attains this resolution through extreme physical separation of anchored residues within single molecules. In contrast, 100-fold expansion followed by ONE microscopy imaging combines more moderate physical magnification with sharpening of the point spread function, to generate high resolution two-dimensional projections, which can be then assembled into three-dimensional reconstructions of individual molecules. 1000-fold expansion also results in larger volumes, with corresponding working distance and imaging time requirements, vs. those incurred by the smaller volume of 100-fold expansion. We acquired two dimensional ONE microscopy images of individual GFP molecules (**Fig. 4A**). For each condition, these 2D projections were combined to reconstruct a three dimensional GFP model from the ensemble of views (**Fig. 4B**; see **Supplementary Note 2**). An effective spatial resolution was then estimated using Fourier shell correlation (FSC) between the reconstructed volume and the reference crystal structure of GFP (PDB entry 1EMA, at 1.90 Å resolution; [5]). Reconstructed resolution improved with increasing expansion, measuring 17.37 Å at 10× expansion (n = 885 from 3 independent gels [5], 14.54 Å at 50× expansion (n = 5,590 from 3 gels), and 11.63 Å at 100× expansion (n = 5,324 from 3 gels) (**Fig. 4C**). The FSC comparison to the 3D GFP structure provides a direct structural benchmark, showing that increasing physical expansion improves the correspondence between the reconstructed GFP density map and the crystallographic structure at finer length scales. Importantly, expansion microscopy does not recover all atomic positions. Anchoring occurs through chemically reactive amino acid side chains, such as lysine residues modified via NHS ester acrylate, where each modified side chain is covalently incorporated into the polymer network during gelation. In contrast to electron microscopy, which detects electron density from essentially all atoms, expansion microscopy reports only the positions of anchored residues. The distance between adjacent anchored side chains can vary. For example, neighboring lysines may be separated by approximately 0.44 nm to 1.41 nm depending on side chain orientation, such as parallel or antiparallel configurations (**Supplementary Fig. 5**). Thus, the fundamental sampling limit is defined by amino acid spacing rather than atomic spacing, placing an upper bound near amino acid level resolution of approximately 10 Å. The 11.63 Å reconstruction at 100× expansion approaches this chemically imposed limit.

**Figure 4.**
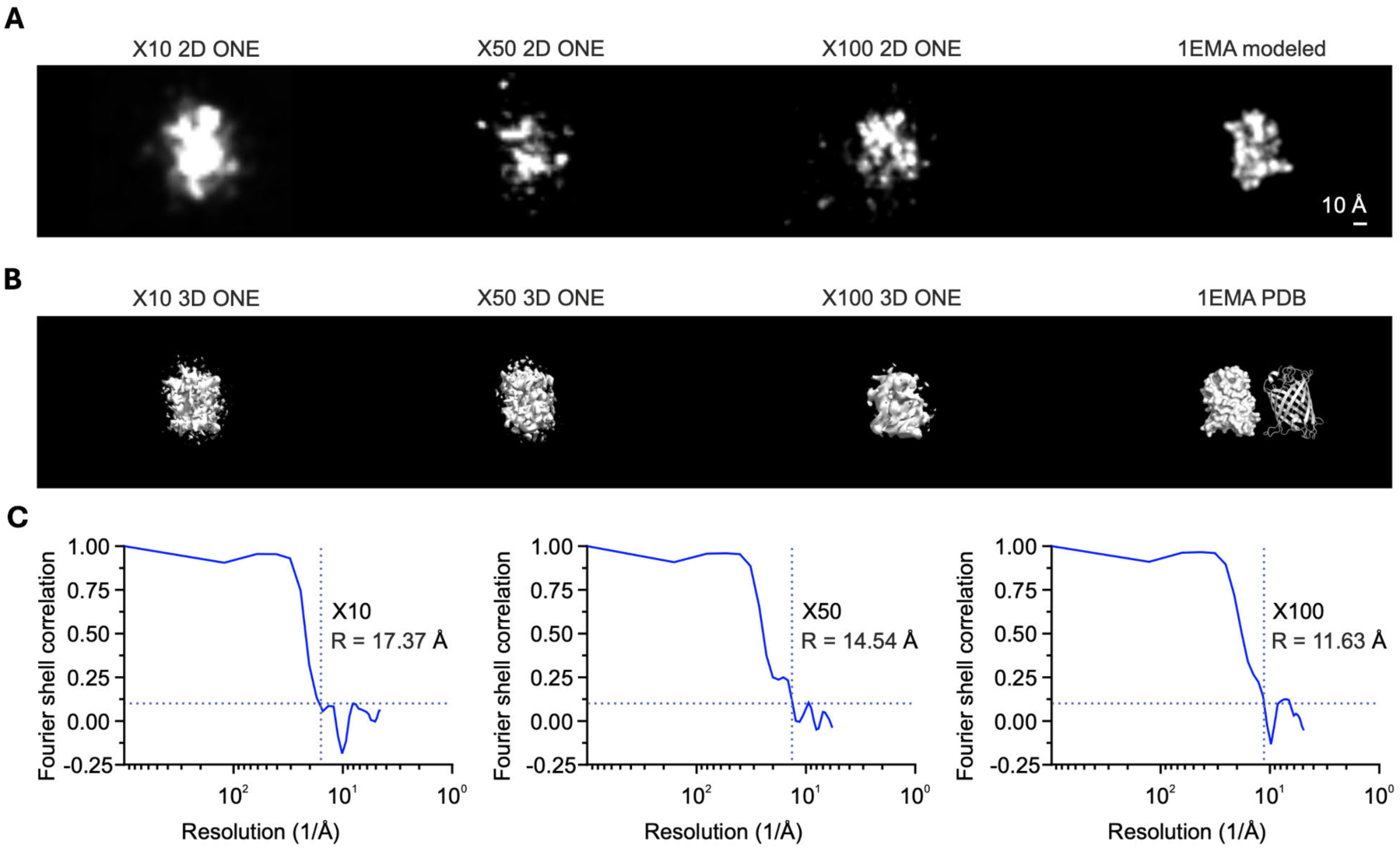
Imaging of GFP after physical expansion with xy resolution enhancement using ONE microscopy. (A) Segmented GFP molecules imaged by ONE microscopy at different linear expansion factors (×10, ×50, ×100), shown alongside the GFP crystal structure (PDB: 1EMA). Images are representative of the full segmented populations at each condition (×10, n = 885 from 3 independent gels [5]; ×50: n = 5,590 from 3 gels; ×100: n = 5,324 from 3 gels). The structural model was rendered from the 1EMA atomic coordinates and convolved with the experimentally determined ONE point spread function to simulate imaging. Scale bar, 10 Å. (B) Three-dimensional reconstructions generated by alignment and averaging of segmented molecules at each expansion factor (see A for molecule counts). The same reconstruction pipeline was applied across datasets. The 1EMA structure is shown for comparison. (C) Fourier shell correlation (FSC) analysis of the 3D reconstructions, computed from independently reconstructed half-datasets. Estimated resolution improves from ∼17 Å at ×10 expansion to ∼11 Å at ×100 expansion.

### Simulated identifiability of the human proteome under 1000× expansion

In the single color lysine implementation utilized above, lysine side chains are first covalently anchored to the hydrogel using NHS ester acrylate. Following proteinase K digestion, peptide backbones are cleaved, and the newly exposed N termini on gel anchored fragments are then labeled with an NHS ester fluorophore. As a result, all fluorescence from a given fragment is registered at the coordinate defined by the original lysine anchoring site. This strategy is compatible with single color labeling, but it does not generalize to multicolor mapping, as it might enable the identification of many different amino acid identities. To enable true multicolor residue-resolved mapping, each residue class must be targeted separately, to ensure its fluorescent labeling with a distinct-spectrum fluorophore, as well as gel anchoring, before expansion. In short, we would need a trivalent molecule that would target a given amino acid selectively, allow for coupling to the polymer backbone, and permit fluorophore attachment.

While creating such a molecule is beyond the scope of the current study, which is focused on the principles of 1000× expansion, we sought to computationally model what this might look like, to guide future developments in the field. In particular, assuming that such trivalent molecules for residue-specific anchoring and labeling could be made, we asked whether such residue-resolved 3D spatial fingerprints, under realistic 1000× expansion distortion, could help support proteome-scale molecular identification of individual molecules *in situ*. For true *in situ* identification in cells or tissues, nearby fragments originating from different proteins would need to be assigned to their correct parent molecules. This remains a bioinformatic challenge that is beyond the scope of the present study, which focuses on establishing the underlying expansion chemistry and molecular-scale spatial preservation. To determine this, we performed computational simulations of the fraction of the human proteome uniquely distinguishable from the coordinates of chemically labeled surface residues. Identifiability was evaluated using RMSD after rigid Kabsch alignment of 3D residue coordinates of expanded proteins compared to ground truth PDB entries (**Fig. 5**). Simulations were performed on 23,391 human proteins with solved structures, using surface-exposed residues as candidate anchoring and labeling sites. Expansion-induced distortion was modeled by adding independent Gaussian noise to residue coordinates, with values grounded in the experimentally measured RMSD distribution for 1000× expanded mCLING peptides (mean 0.573 ± 0.151 nm; **Fig. 3C.i**). Accordingly, 5 Å distortion reflects typical experimental performance, while 10 Å distortion represents a more conservative, suboptimal condition exceeding the 90th percentile. Labeling efficiency was modeled by randomly subsampling surface residues of each class (e.g., 0.5 corresponds to labeling 50% of sites). A protein was considered identifiable if its closest RMSD match was its own undistorted structure (**Fig. 5A**).

**Figure 5.**
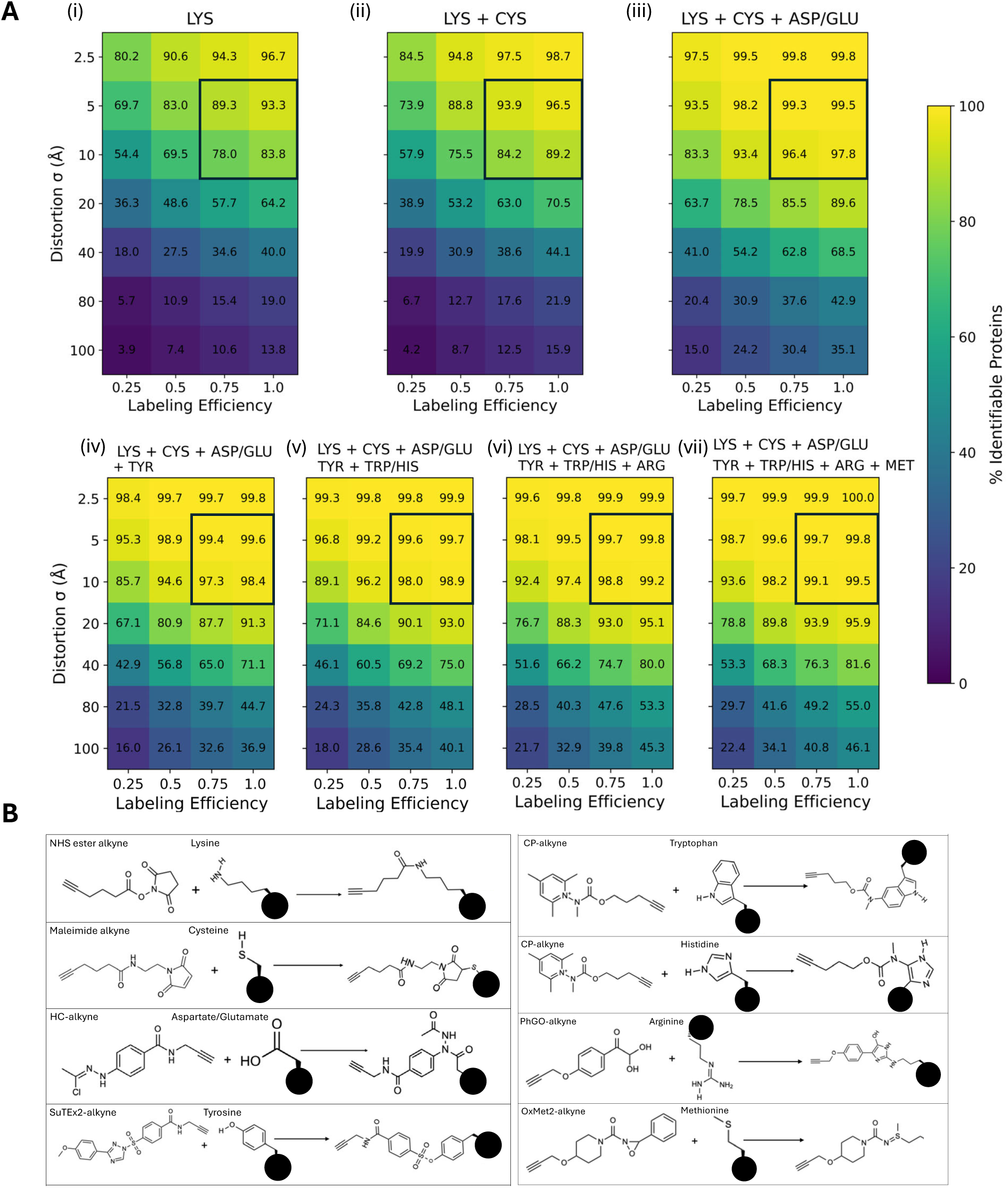
Simulation of human proteome identifiability using existing and prospective residue-specific labels. Protein identifiability was simulated by labeling surface-exposed residues in a single static reference structure per protein (∼23,000 human proteins), without modeling conformational variability. Fragment separation was simulated by adding Gaussian spatial distortion to labeled residue coordinates. Identifiability was evaluated by comparing each distorted structure to all others using RMSD after Kabsch alignment; a protein was considered identifiable if its closest match was its own undistorted reference structure. (A) Fraction of human proteins that remain uniquely identifiable when only surface residues are labeled. Heat maps show identifiability as a function of labeling efficiency (x-axis) and spatial distortion (y-axis; Gaussian noise with the indicated standard deviation added to residue coordinates). Anticipated target conditions are outlined in black. (i) Surface lysines (1 color). (ii) Surface lysines and cysteines (2 colors). (iii) Surface lysines, cysteines, and aspartate/glutamate (3 colors). (iv) Surface lysines, cysteines, aspartate/glutamate, and tyrosines (4 colors). (v) Surface lysines, cysteines, aspartate/glutamate, tyrosines, and tryptophan/histidines (5 colors). (vi) Surface lysines, cysteines, aspartate/glutamate, tyrosines, tryptophan/histidines, and arginines (6 colors). (vii) Surface lysines, cysteines, aspartate/glutamate, tyrosines, tryptophan/histidines, arginines, and methionines (7 colors). (B) Chemoselective reaction mechanisms corresponding to the residue-specific labels evaluated in panel A [52].

With single-color lysine labeling, 89.3–93.3% of proteins were identifiable at 5 Å distortion, for 75%–100% labeling efficiency, and 78.0–83.8% at 10 Å distortion and the same labeling efficiency (**Fig. 5A.i**). Adding cysteine labeling increased identifiability to 93.9–96.5% and 84.2–89.2%, respectively, assuming the same labeling efficiency as for lysine (**Fig. 5A.ii**). Incorporating acidic residues as a third class increased identifiability to 99.3–99.5% at 5 Å and 96.4–97.8% at 10 Å, again holding labeling efficiency constant (**Fig. 5A.iii**); additional residue classes yielded diminishing gains (**Fig. 5A.iv–vii**). Thus, these simulations demonstrate that near-complete proteome identifiability (>99%) might be achievable with only three chemically addressable residue classes under experimentally measured 1000× expansion distortion, establishing residue-level spatial mapping as a potentially sufficient basis for large-scale *in situ* protein identification.

## Discussion

Here, we present 1000ExM, a four-network IPN (interpenetrating polymer network) formed by an initial ionic gel and three successive ionic-in-ionic casting steps. This process yields ∼1000-fold expansion in each dimension while preserving three-dimensional (3D) molecular geometry. Physical expansion converts sub-nanometer-scale molecular distances into micrometer-scale separations that are resolvable using standard light microscopy, enabling direct readout of 3D amino acid positions without the need for averaging. Across multiple independent statistical analyses on three molecular species, we observe preservation of spatial information (i.e., effective resolution) down to individual amino acid residues; using the peptide mCLING, preservation was demonstrated by a root-mean-square deviation (RMSD) of ∼6 Å between experimentally observed molecular coordinates and molecular-dynamics reference structures. When experimentally measured distortion and incomplete residue readout are taken into account, simulations across the human proteome (23,391 canonical proteins) suggest that spatial amino acid coordinate patterns alone are sufficient to uniquely identify most proteins.

Our work builds on a decade of innovation in iterative ExM methods, which expand an initial gel, stabilize it with a neutral re-embedding gel, and then cast a second swellable gel to enable further expansion. This strategy, used in iExM [7], iU-ExM [8], and pan-ExM [17], typically achieves 16- to 25-fold linear expansion. This established regime does not approach the expansion required for amino acid–level analysis. DMAA gels [18] have been shown to preserve protein structure [5], but only within this lower expansion range, and it remains unclear whether other chemistries can achieve comparable fidelity. Here, we introduce an architecture that enables a step-change in expansion factor: we eliminate the neutral stabilizing gel and instead re-embed expanded gels by re-exposure to the same charged monomer solution. This produces shrinkage perhaps unexpectedly comparable to neutral re-embedding, and enables ionic-in-ionic casting, which might be expected to fail due to collapse of the expanded polyelectrolyte network. This approach yields a four-network expansion sequence in which each stage produces net expansion, resulting in ∼1000× linear expansion, far beyond prior iterative methods. The current gel architecture could be a platform for future innovation: for example, repeated monomer incubations before a given polymerization step pushed expansion factor up to 2000x-2500x, in some exploratory work that was not the focus of the current paper, but which points to rich downstream possibilities. A natural question is how ∼1000× expansion can occur while preserving nanoscale structure. One possibility is bond scission during swelling, followed by reincorporation of polymer fragments into successive interpenetrating gels during free-radical polymerization. Alternatively, elastic stretching may still occur in DMAA-based gels, as observed for hydrogels with long crosslinkers [19].

We highlight one future application that could, in principle, be enabled by 1000ExM: *in situ* protein identification. The functions of biological entities are governed by the organization of their biomolecules. Their copy numbers, locations, interactions and structures determine corresponding biological outputs, across different scales, from organelles to cells, tissues and whole organisms. As biomolecules work largely through contact with one another, mapping out their spatial locations at the resolution of their interactions is key to understanding biology. This can only be achieved by technologies that can localize molecules and decode their identities and shapes, at a resolution better than molecular size. Such technologies are still missing, for the protein domain. Their structural complexity implies that not only their identities, but also their local shapes and arrangements need to be revealed. The technology of deep visual proteomics, in which regions identified by fluorescence microscopy are microdissected and analysed by mass spectrometry, currently achieves cellular resolution [23]. Even if protein mass spectrometry could be brought to higher resolutions, which is being explored with expansion microscopy [24–26], its sensitivity remains short of single molecules [27,28]. Of course, for proteins that are touching, or in clusters, assigning expanded protein fragments to their parent protein would be a further bioinformatic issue to be solved, but might be feasible, given the richness of data available, especially if multiple colors are available for labeling of different amino acids.

To address protein mapping via imaging, proteins are currently labelled either through genetic tagging or affinity tools. While the wide majority of experimenters describe up to 3-5 different proteins, by using a handful of antibodies and colour channels, a few laboratories have enhanced this procedure, to attain impressive multiplexing results. A brute force approach has been used in “toponome” imaging, in which dozens of antibodies are applied to samples and imaged sequentially [29]. Multiplexed imaging can also be achieved by combining antibodies carrying different metal (lanthanide) tags [30] or DNA barcodes [31]. Overall, up to 100 targets can be imaged with such methods, at resolutions of 200-300 nm. To improve the resolution, DNA barcoding has been combined with DNA-PAINT [32]. This enables resolutions of ∼15 nm, for up to ∼30 targets. Expansion-based multiplexing methods that use antibodies, DNA conjugated or not, have also been put forth [33]. However, expansion workflows require covalent anchoring to residues followed by enzymatic digestion or chemical softening prior to swelling. These steps can disrupt structure or erase epitopes to the point of leaving antibodies nonfunctional. In addition, the use of antibodies, or other affinity tools, implies that the exact protein orientations and shapes remain obscure in these experiments. Finally, trusted antibodies are not available for all proteins, much less protein isoforms (e.g., splice variants), and generating new ones remains laborious and time-consuming.

An alternative analysis relies on cryo-electron tomography (cryo-ET). For example, a stereotypically organized biological object, the axonemal microtubule arrangement in sperm, was averaged across large image numbers, to obtain reconstructions with resolutions of 3.5-6 Å; this analysis was sufficient to reveal the positions and structures of unknown proteins, whose identities were later revealed by comparison to AlphaFold-derived structural predictions [34]. This approach is promising, but is limited by the need to average objects that are stereotypically organized, and have a sizable molecular weight, to obtain structures. When analysing loosely organized objects, as synaptic vesicles, only a handful of proteins can be revealed by cryo-ET approaches [35].

In terms of resolution, 1000ExM enters the optics domain of “Ångstrom-scale resolution”, which already includes several super-resolution optics technologies, from MINimal emission FLUXes microscopy (MINFLUX) [36] to Cryogenic Optical Localization in 3D (COLD) [37] or REsolution Enhancement by Sequential Imaging (RESI) [38]. These technologies rely on very precise determinations of fluorophore positions [39]. The fluorophores can be placed either on different biological molecules, or on different sites within single molecules, resulting in measurements of structural features [40]. However, the fact that these technologies require the decoration of native biological samples with fluorophores is a strong limitation. The fluorophore size is comparable to that of the biological molecules, with a small dye, as fluorescein, being only about 3- to 4-fold smaller than a compact protein as GFP, implying that few sites on the molecules can be labeled and visualized. This problem is alleviated by the application of fluorophores after expansion, implying that far more components of biological molecules are revealed, leading to true molecular visualizations. This has been exploited in the generation of EM-like images for cells and tissues, both with limited resolution, for wide-scale tissue analyses [9] and with nanometer resolution, for the analysis of protein structures [5]. Overall, we concur with a recent review of the field [39], in that ExM-based analyses, including 1000ExM, can reveal substantially more components of biological samples than optics-based approaches, simply due to the ability to fit more fluorophores within an expanded structure.

To make 1000ExM work in practice, imaging is performed on a confocal microscope equipped with high-sensitivity detectors (e.g., hybrid detectors such as HyD-X) and a high numerical aperture objective (e.g., 100× oil immersion, NA ≥ 1.4–1.51). In this study, single fluorophore labels were used. After ∼1000× expansion, fluorophores are no longer diffraction limited, but their signal is diluted, making samples extremely dim and often difficult to locate. At the same time, the large expansion factor imposes substantial constraints on imaging volume and acquisition time. At 1000× linear expansion, 1 nm maps to 1 µm, so a ∼10 nm protein expands to ∼10 µm, comparable to a non-expanded cell, and a ∼10 µm cell expands to ∼10 mm, comparable to the size of a mouse brain. This increase in specimen size directly scales the imaging burden, making throughput a limiting factor and motivating the use of approaches such as lattice light sheet microscopy [41]. In potential support of the need for long acquisitions, Supp Fig. 31 shows that with our modified protocol, expanded gels maintained on a dry surface at room temperature changed in linear expansion factor by only 0.3 ± 0.5% over 98 hours, which could potentially synergize with extended automated imaging.

In addition to these practical constraints, there are fundamental limits to precision. The positions of amino acids depend on the locations of their side chains at the time of gel anchoring. For example, lysine side chains are ∼1 nm in length, and their terminal amines can sample positions over ∼2 nm under conditions of rotational freedom. Although this variability is reduced in structured protein domains, it nonetheless limits reconstruction precision to approximately the side chain length, on the order of ∼10 Å. This is consistent with the precision obtained for GFP and remains below the resolving power of the best cryo-EM approaches [42]. Cryo-EM achieves its highest resolution on purified structures of intermediate size, but in intact cellular environments resolution is substantially reduced, and small, low-molecular-weight molecules cannot be resolved.

Despite these practical and physical constraints, 1000ExM produces a 3D map of amino acid positions that could, in principle, enable direct protein identification in intact cells and tissues. More broadly, it defines a data modality based on spatially resolved, multicolor molecular coordinates of biomolecular building blocks, which could, with future work, extend beyond proteins to other classes of biomolecules composed of repeated subunits with distinct chemical reactivities that could be similarly resolved.

## Author Contributions

The four-network ionic interpenetrating polymer network architecture enabling 1000× linear expansion was conceived and reduced to practice by H.H. Gelation experiments were initially performed by H.H., with additional experiments subsequently performed by D.K. and A.H. The single-molecule validation experiments using nanobodies, mCLING, and GFP were designed by A.H.S. and S.O.R., together with H.H. and E.S.B. Other experimental designs, steps and imaging were done by H.H., A.H.S., D.K., A.N., K.R., and A.H. Two-dimensional analysis for Figure 2 was done by S.O.R. Three-dimensional analysis for Figure 3 was done by H.H. The GFP analysis for Figure 4 was performed by J.A., M.M., and S.C. Molecular dynamics simulations were done by B.J. and B.B. Molecular dynamics analysis was done by H.H. Simulation results for Figure 5 were generated by H.H. The manuscript was written by H.H. and E.S.B. Supervision was provided by A.H.S., S.O.R., and E.S.B.

## Acknowledgements

H.H. and E.S.B. thank Robert S. Langer for his early encouragement and support, and for thoughtful discussions and insights that contributed to the development, validation, and application of 1000ExM. H.H. and E.S.B. thank Paula Hammond, Jeremiah Johnson, Kecheng Wang, and Michael Lang for discussions on high-expansion hydrogels from polymer chemistry and polymer physics perspectives. H.H. and E.S.B. thank Deblina Sarkar, Jinyoung Kang, Tay Shin, Camille Mitchell, Shiwei Wang, Hao Wang, Anu Sinha, and Debarati Ghosh for discussions around expansion. H.H. and E.S.B. thank Chi Zhang, Yixi Liu, and Amy Keating for discussions on protein identification strategies. H.H. and E.S.B. thank Jeremy Wohlwend, Mateo Reveiz, and Regina Barzilay for discussions on machine learning applications. H.H. and E.S.B. thank Matteo Mazzoca, Anders Hansen, Maya Anjur-Dietrich, Penny Chisholm, Sangeeta Bhatia, Daniel Kim, Viktor Adalsteinsson, Dan Anderson, Evan Collins, Li-Huei Tsai, Joey Davis, Samantha Sedor, Eliezer Calo, Shahar Bracha, and Shoh Asano for helpful discussions on biological applications. H.H. thanks Burcu Ataman and Chi Zhang for teaching and mentorship. H.H. thanks Emily Cronin Furman and Jeff Wykoff for advice on imaging. H.H. thanks Eric Alm, Katya Moniz, Anna Rasmussen, and Neil Rasmussen for support through the Rasmussen Fellowship and for early encouragement. H.H. and E.S.B. thank Chris Vallace, Lauren Foster, Chris Baxter, and Patrea Pabst for discussions around IP. We thank Abed Alrahman Chouaib for guidance on density point generation for GFP 3D reconstruction. We thank Felipe Opazo for GFP purification. We thank Sven Truckenbrodt and Felipe Opazo for valuable comments on the manuscript. We thank Maxwell Hu for testing the 1000ExM protocol and providing insights that informed the updated protocol.

The project leading to this application received funding from the European Research Council under the European Union’s Horizon 2020 research and innovation programme, grant agreement No. 835102, to E.S.B. and S.O.R. E.S.B. also acknowledges funding from Ashar Aziz, Lisa Yang, W.M. Keck Foundation, Richard King Mellon Foundation, HHMI, John Doerr, NIH R01AG087374, NIH R01AG070831, NIH R01EB024261, Kathleen Octavio, Lore McGovern, Jed McCaleb, James Fickel, Good Ventures/Open Philanthropy, Tom Stocky, and Avni Shah. B.J. and B.B. acknowledge funding from NIH R35GM141861. S.O.R. acknowledges funding from the German Research Foundation, Deutsche Forschungsgemeinschaft, grants GRK2824 and SFB1690/A05, and from the German Ministry for Education and Research, grant DATI-Pilot. The authors gratefully acknowledge the computing time granted by the Resource Allocation Board and provided on the supercomputer Emmy/Grete at NHR-Nord@Göttingen as part of the NHR infrastructure. The calculations for this research were conducted with computing resources under the project nib00040.

## Methods

### 1000-fold linear expansion of proteins nanorulers

#### Anchoring protein nanorulers

Proteins were modified with acryloyl-X, SE (AcX; A-20770, Thermo Fisher Scientific). For mCLING peptides, 2 µL of mCLING-Atto 647N (710 006AT1, Synaptic Systems; 1.0 nmol mL⁻¹ stock) were mixed with 100 µL DPBS and 3 µL acryloyl-X, SE (AcX; 10 mg mL⁻¹ in DMSO). For nanobodies, 1 µL FluoTag-X2 anti-ALFA AbberiorStar635P (N1502, NanoTag Biotechnologies; 10 µM stock) was mixed with 100 µL DPBS and 3 µL AcX (10 mg mL⁻¹ in DMSO). For GFP, 1 µL GFP (10 µM stock) was mixed with 100 µL DPBS and 3 µL AcX (10 mg mL⁻¹ in DMSO). GFP was purified as previously described [5]. Briefly, His-tagged GFP was expressed in NebExpress Escherichia coli and cultured in Terrific Broth at 37 °C, followed by induction with 0.4 mM IPTG for 16 h at 30 °C. Bacterial pellets were lysed by sonication on ice in 50 mM HEPES (pH 8.0), 500 mM NaCl, 5 mM MgCl₂ and 10% glycerol. After centrifugation to remove debris, the supernatant was incubated with Ni²⁺ affinity resin at 4 °C. The resin was washed extensively, and GFP was enzymatically eluted via SUMO protease-mediated cleavage of the N-terminal His-tag. Protein concentration was determined by absorbance at 280 nm using the calculated extinction coefficient. All AcX–nanoruler mixtures were pipetted onto 18-mm round coverslips (Alkali Scientific, SM036) and spread evenly. Coverslips were placed in Petri dishes, which were covered with parafilm, then sealed with lids, and incubated at 4 °C overnight. After incubation, coverslips were partially air-dried in a chemical fume hood for ∼30 min, until only a thin residual liquid layer remained and the solution became visibly viscous. Excess liquid (10–15 µL) was gently removed, leaving a uniform thin protein film without visible salts or particulates. Protein-coated coverslips were used immediately for gelation or stored at 4 °C for up to 1 h.

#### Preparation of 1000ExM gelation solution

The 1000ExM gelation solution consisted of sodium acrylate (25.6% w/v; AK Scientific, R624), N,N-dimethylacrylamide (DMAA, 44.2% v/v; Sigma, 274135), N,N,N′,N′-tetramethylethylenediamine (TEMED, 0.037% v/v; Sigma, T9281), and potassium persulfate (KPS, 0.145% w/v; Sigma, 216224) in acidified Tris buffer. The full preparation protocol is described in **Supplementary Note 1,** including details of many steps below, which are in summary form. The monomer solution was degassed with nitrogen for 10 min and used immediately.

#### First round of gelation

Activated monomer solution (80 µL) was pipetted onto a hydrophobic glass slide (CytoSlide Fluorosilane, CYTONIX). The protein-coated coverslip was inverted sample-side down onto the droplet. The sealed Tupperware container (Rubbermaid, B079M8FPTW) was purged with nitrogen gas for 1-3 h through a punctured hole (∼2 mm diameter) large enough to insert a pipette tip, and then left overnight to allow polymerization to complete.

#### Digestion and first round of expansion

Proteins anchored to the hydrogel were digested with proteinase K (8 U mL⁻¹; P4850, Sigma-Aldrich, now Merck) in digestion buffer containing 800 mM guanidine HCl, 2 mM CaCl₂, and 0.5% Triton X-100 in 50 mM Tris (8382J008706, Merck). Digestion was performed at 50 °C for >16 h. Following digestion, gels were expanded ∼3× in the digestion buffer, then transferred to deionized water and incubated at 20 °C with three successive water exchanges (1 h each) until the gel reached ∼18× linear expansion.

#### Multiple rounds of gelation and expansion

For the second gelation step, fully expanded (∼18×) gels were transferred to a 6-well plate and infiltrated with activated monomer solution, during which the gels contracted. Activated monomer solution was added in sufficient volume to fully immerse the gels from all sides. The container was degassed with nitrogen for 10 min and incubated on a shaker for 35 min to facilitate infiltration. Gels were then rapidly removed, placed sample-side down on hydrophobic glass, covered with a coverslip within ∼10 s, and transferred to a sealed container purged with nitrogen gas for 1 h. Polymerization proceeded overnight (>8 h). The resulting composite gel was expanded in water to ∼100× linear expansion. For the third gelation step, fully expanded (∼100×) gels underwent the same monomer infiltration, degassing, and overnight polymerization procedure, followed by expansion in water to ∼500× linear expansion. For the fourth gelation step, fully expanded (∼500×) gels underwent an additional cycle of monomer infiltration, degassing, and overnight polymerization, followed by expansion in water to ∼1500× linear expansion. Detailed protocol information is provided in **Supplementary Note 1**.

#### Post-expansion labeling and stabilization

Gels were post-expansion labeled with NHS-ester fluorescein (46409, Thermo Fisher Scientific). Samples were labeled using a 20-fold molar excess (compared to peptide) of NHS-ester fluorescein in NaHCO₃ buffer at pH 8.3 for 3 h before the washing procedure that induced the final expansion. Gels were equilibrated in 1× DPBS to a stable ∼1000× linear expansion state. In this partially equilibrated condition, the gels do not exhibit rapid shrinkage, making them amenable to overnight imaging. A biological replicate of a 1000× gel was defined as one complete four-round gelation sequence each network cast independently.

### Image acquisition

#### ONE imaging

Images were acquired on a Leica STELLARIS 8 microscope using HyD-X detectors and a 1.51 NA 100x oil immersion objective with an xy pixel size of 91 nm. Imaging was performed with a 12 kHz resonant scanner in line-scan format with line accumulation of 3, 25 ms acquisition time, and 8-bit depth (Dynamic Signal Enhancement (DSE) 15, weight 0.4). Excitation used a 633 nm laser at 40% power and a 488 nm laser at 30% power. For each 11.72 × 11.72 µm field of view, 2000 sequential frames acquired with the 12 kHz resonant scanner were used as the input to the ONE plugin [5,43,44],(https://github.com/Rizzoli-Lab/ONE-Microscopy-Java-Plugin), that builds on SRRF NanoJ core [45], with parameters temporal radiality auto-correlation (TRAC) = 4 and radiality = 10. Additional datasets were acquired on a Leica SP5 microscope using an 8 kHz resonant scanner with line accumulation of 3 and processed using TRAC = 4 and radiality = 10.

#### Confocal z-stack acquisition

Confocal z-stacks were acquired on a Leica STELLARIS 8 microscope equipped with HyD-X detectors using a 100× oil immersion objective (NA 1.51). The z-step size was 0.20 µm and the xy pixel size was 0.091 µm. Excitation was performed with a 647 nm laser at 40% of 3.5 mW power (HyD-X4 detector) and a 488 nm laser at 30% of 1.6 mW power (HyD-X2 detector).

### Image analysis

#### Nanobody analysis

For the analysis of distances between nanobody signals (**Fig. 2B**), spots were identified by thresholding band-pass filtered images, relying on empiric thresholds and band-pass filters (adjusted so that realistic signals would be obtained for every image set, as evaluated by experienced investigators), organized in the form of semi-automated routines in Matlab (version 2023b, The Mathworks, Inc, Natick, MA, USA). Their intensity center positions were then determined, to measure the respective distances between the spots (in the same colour channel, corresponding to Star635P).

#### mCLING 2D line-scan analysis

The averaging analysis of mCLING molecules was performed using Matlab. In brief, molecules were detected automatically, as particles with intensities above an empirically-derived threshold (derived as in the previous section), in band-pass filtered images, relying on the Atto647N channel. All particles were centered on the intensity maxima of the respective Atto647N spots. The fields containing the individual particles were then rotated to fit their fluorescein signals in the horizontal position, lateral to the Atto647N spot. They were then all overlaid, generating an average image for the respective condition. The same exact analysis was performed for a mirror image of the fluorescein signals, and the mirror image result was subtracted from the original raw image of the fluorescein signals, thereby obtaining a noise-corrected final image. The alignment procedure will place any bright noise events in a horizontal position, implying that some amount of fluorescence will be observed on the horizontal line, even if no real signals are present in the sample. It is therefore important to subtract a random image (a mirror image) from the original results, to account for such effects. The positions of fluorescence intensity peaks were determined by line scans across the final images and compared to those obtained from a structural model of mCLING. A set of peptide conformations was generated by molecular dynamics (MD) simulation. For each conformation, chemically accessible anchoring sites were identified and fluorophores were assigned to those positions, yielding an ensemble of model peptides defined by predicted fluorophore coordinates comprising one Atto647N and four or five fluoresceins (as described in **Supplementary Figures 4 and 6**). Thousands of molecules were 2D-projected on top of each other, which was, as for the experimental situation, the Atto647N fluorophore. To account for errors in the determination of the fluorophore positions, due to the uncertain emission position from the relatively large fluorophore molecule, a fluorophore positioning error of up to 3.6 Å (for fluorescein) and up to 13.6 Å (for Atto647N) was incorporated. The pixel size of each model result was adjusted to the pixel size of each microscopy analysis, and the peak positions were determined.

Plots and statistics were generated using SigmaPlot 14 (Systat Software Inc., San Jose, CA, USA), or using Matlab. Statistics details are presented in the respective figures.

#### mCLING 3d analysis

##### 3D emitter detection

For 1000-fold expanded mCLING gels, eight 3D volumes were captured across three independent gelation experiments. The x, y, and z dimensions and total imaged volume for each of the eight regions are summarized in **Table S1.**

After acquisition, raw z-stacks were smoothed using an anisotropic Gaussian filter to suppress noise while preserving point-like emitters. The Gaussian standard deviation was arbitrarily set to 0.05 µm along each axis and was implemented in voxel units by dividing 0.05 µm by the measured sampling along each dimension. Specifically, σx and σy were set to 0.05 µm divided by the x, y pixel size, and σz was set to 0.05 µm divided by the z-step size.

Local maxima were detected by sliding a 3 × 3 × 3 voxel window through the filtered z-stack one voxel at a time. A voxel was classified as a local maximum if its intensity exceeded that of all 26 neighboring voxels. The window size was calculated to match the expected spatial extent of a single emitter: after Gaussian smoothing (σ = 0.05 µm), the effective emitter FWHM is on the order of 0.1–0.15 µm, corresponding to approximately 1–2 voxels in x, y and ∼1 voxel in z. A 3 × 3 × 3 neighborhood therefore captures the full emitter footprint while minimizing merging of nearby emitters. At this stage, detections are dominated by background and detector noise, as single-emitter signals are only slightly above dark counts. The number of Atto647N and fluorescein emitters detected after local maxima detection, and the fluorescein-to-Atto647N ratio for each region, are reported in **Table S2**.

Next, local maxima were filtered using two quantitative criteria designed to suppress noise-driven detections while retaining point-like emitters. First, candidate emitters were required to have a center-voxel intensity greater than the regional mean intensity of all detected maxima. Noise-driven local maxima arise from symmetric fluctuations around the background and therefore cluster near the mean intensity, whereas true single emitters would be expected to produce a higher-amplitude peak. This criterion preferentially removes maxima consistent with stochastic background and detector noise while retaining higher-contrast peaks. Second, the apparent size of each candidate was arbitrarily defined as the number of voxels within a local window whose intensity exceeded 50% of the center-voxel intensity, without requiring spatial contiguity. True emitters produce a spatially extended intensity profile determined by the point-spread function, resulting in multiple above-threshold voxels. In contrast, noise-driven peaks are typically confined to one or a few voxels. Candidates with an apparent size of ≥ 12 voxels were therefore retained as likely single emitters, while smaller features were excluded as noise. Emitter counts after intensity- and size-based filtering are summarized in **Table S3**.

Remaining candidates were evaluated for point spread function (PSF) conformity, as the lateral intensity profile of an isolated fluorophore imaged under confocal conditions is well approximated by a Gaussian function. For each candidate, a local cubic patch centered on the peak voxel was extracted and compared to a Gaussian model with the same standard deviation as the preprocessing filter. The Pearson correlation coefficient between the observed intensity distribution and the Gaussian template was computed to quantify agreement. A threshold of r ≥ 0.7 was applied to retain peaks whose spatial profiles are consistent with a single diffraction-limited emitter, while remaining tolerant to realistic deviations from ideal Gaussian shape due to shot noise, pixelation, and sampling anisotropy. Final emitter counts after Gaussian shape filtering are reported in **Table S4**.

Overall, intensity and size thresholding follow standard fluorescence microscopy practices [46,47] for emitter detection. By additionally enforcing quantitative PSF conformity through Gaussian correlation analysis, we extend these conventional steps with a physically grounded shape validation, increasing robustness while remaining consistent with established analytical workflows.

##### k-nearest neighbor (kNN) analysis of emitter distances

Using Atto647N and fluorescein emitters obtained from three-dimensional (3D) z-stacks after emitter detection, three-dimensional Cartesian coordinates were used to compute Euclidean k-nearest neighbor (kNN) distances for k = 1–5. Distances were computed separately for Atto647N-to-fluorescein and fluorescein-to-fluorescein emitter pairs using the cKDTree implementation in the SciPy spatial module (scipy.spatial.cKDTree). Distances were aggregated at three pooling levels: all gels and all regions combined (**Table S5**), all regions within each individual gel (**Tables S6–S8** for gels 1–3).

Each statistics table reports values for k = 1–5 and includes the number of distances (n), mean, standard deviation, median, 25th and 75th percentiles (abbreviated P25 and P75), 5th and 95th percentiles (abbreviated P5 and P95), and 1st and 99th percentiles (abbreviated P1 and P99), all in microns. Each table also reports two peak-related metrics: the kernel density estimate (KDE) mode, defined as the location of the global maximum of a Gaussian kernel density estimate fit to the distance distribution, and the peak value, defined as the center of the histogram bin with the highest count, using a bin width of 0.05 µm and Gaussian smoothing with σ = 1 bin.

##### Randomization of k-nearest-neighbor distances

Randomization of locations was used to assess whether measured k-nearest-neighbor (kNN) distance distributions reflect structured, molecule-related spatial relationships rather than random spatial coincidence within the imaged volume. For each of the eight 3D imaging volumes, the measured x, y, z coordinates of all detected Atto647N and fluorescein emitters were used, with each emitter represented by a single 3D coordinate. Null datasets were generated by randomizing emitter locations within each imaging volume while preserving the number of emitters and the overall volume geometry. Specifically, x, y, and z coordinates were independently resampled for each emitter from uniform distributions spanning the experimentally observed x, y, and z bounds of each imaging volume, thereby preserving the overall spatial extent and emitter density while eliminating molecule-specific spatial correlations. For each observed and randomized dataset, kNN distances (k = 1–5) were computed using the same analysis pipeline. This randomization procedure was repeated 100 times per imaging volume, generating a null ensemble of kNN distance distributions. Observed and randomized distributions were compared using the two-sample Kolmogorov–Smirnov (KS) test, with the KS D statistic defined as the maximum absolute difference between their empirical cumulative distribution functions and used as the primary quantitative measure of deviation from the null hypothesis of spatial randomness. kNN distance distributions and corresponding KS D statistics were summarized across all gels (**Table S9**) and by individual gels (**Tables S10–S12**).

##### Identification of putative peptides

Putative peptides were identified by grouping spatially proximal fluorophores into candidate mCLING molecules using k-nearest-neighbor (kNN) analysis (k = 1–5). Each mCLING molecule contains one Atto647N dye and four to five fluorescein dyes, as defined in **Supplementary Figure 4**. For each Atto647N localization, the five nearest fluorescein neighbors were identified in 3D space.

Because each fluorescein dye originates from a single peptide, fluorescein localizations were assigned to Atto647N dyes using a greedy nearest-neighbor assignment, in which each fluorescein was associated with the closest Atto647N and could not be assigned to multiple peptides. This constraint prevents cross-assignment of fluorescein dyes between nearby peptides and ensures a one-to-one mapping between fluorophores and candidate peptides.

Using this procedure, 10,696 coarse peptides containing one Atto647N dye and 4–5 associated fluorescein dyes were initially identified. To suppress false groupings arising from spatial overlap between neighboring peptides, candidates were filtered using bounds on the first and last kNN distances, which together define the spatial extent of a single peptide. Putative peptides were retained if the first nearest-neighbor distance lay within the 10th–90th percentile of the kNN1 distribution (0.350–1.343 µm) and the last nearest-neighbor distance lay between the 10th percentile of the kNN4 distribution and the 80th percentile of the kNN5 distribution (1.410–2.562 µm).

These bounds were chosen to enforce separation between fluorophores belonging to the same peptide versus those originating from neighboring peptides. Large terminal kNN distances are indicative of cross-peptide associations in densely labeled samples, while unusually small first-neighbor distances might reflect noise, over-segmentation, or repeated detections of a single fluorophore. Constraining only the terminal kNN distances is sufficient: if both the closest and furthest associated fluorescein dyes fall within expected bounds, intermediate kNN distances are implicitly constrained. The selected percentile ranges also align with ground-truth molecular-dynamics distance relationships, in which kNN1 distances fall near 0.5–1.5 nm, kNN4 distances begin near ∼1 nm, and kNN5 distances extend to ∼2.5 nm (**Supplementary Figure 6**), which map closely to the observed experimental distance ranges after expansion. After filtering, 6,284 putative peptides were retained for downstream analysis.

##### RMSD analysis of putative peptides

RMSD analysis was used to quantify agreement between experimentally reconstructed fluorophore positions within expanded peptides and atomic-resolution molecular-dynamics (MD) reference models, and to estimate gel distortion, isotropy, and single-molecule expansion factors. A total of 6,284 putative peptides were analyzed across 8 imaging regions from 3 gels. Each putative peptide consisted of one Atto647N dye and four or five associated fluorescein dyes. As reference structures, an MD ensemble of 1001 peptide conformations was used, with a 5 choose 4 subset representing variability arising from different anchoring, digestion, and labeling outcomes.

For each experimental peptide, fluorophore coordinates were compared independently to each MD conformation. Experimental coordinates were aligned to the MD reference using a similarity transform consisting of rotation, translation, and a single isotropic scale factor, chosen to minimize the sum of squared distances between corresponding attachment sites (Eq. 1). The fitted scale factor captures the relative linear scaling between the experimental coordinates and the MD reference during alignment.

After optimal alignment, the total RMSD was computed as the root-mean-square of Euclidean distances between all matched attachment sites (Eq. 2). To assess directional distortion, per-axis RMSDs were computed by decomposing residual displacement vectors into their x, y, and z components after alignment and calculating the root-mean-square residual along each axis separately (Eq. 3).

The single-molecule expansion factor was then derived from the fitted scale factor using the definition given in Eq. 4, which converts the relative similarity scaling into an absolute linear expansion factor referenced to the nominal 1000× expansion. This definition preserves the sign and magnitude of deviations from the nominal expansion at the level of individual molecules.

This procedure was repeated for each experimental peptide against all 1001 MD conformations. For each peptide, the MD conformation yielding the minimum total RMSD was selected as the best structural match, and the corresponding total RMSD, per-axis RMSDs, and expansion factor were reported (**Fig. 3C.I–C.III**).

Summary statistics for these quantities are reported for all peptides pooled across gels and regions (**Table S13**), stratified by gel (**Table S14**), and stratified by imaging region (**Table S15**).

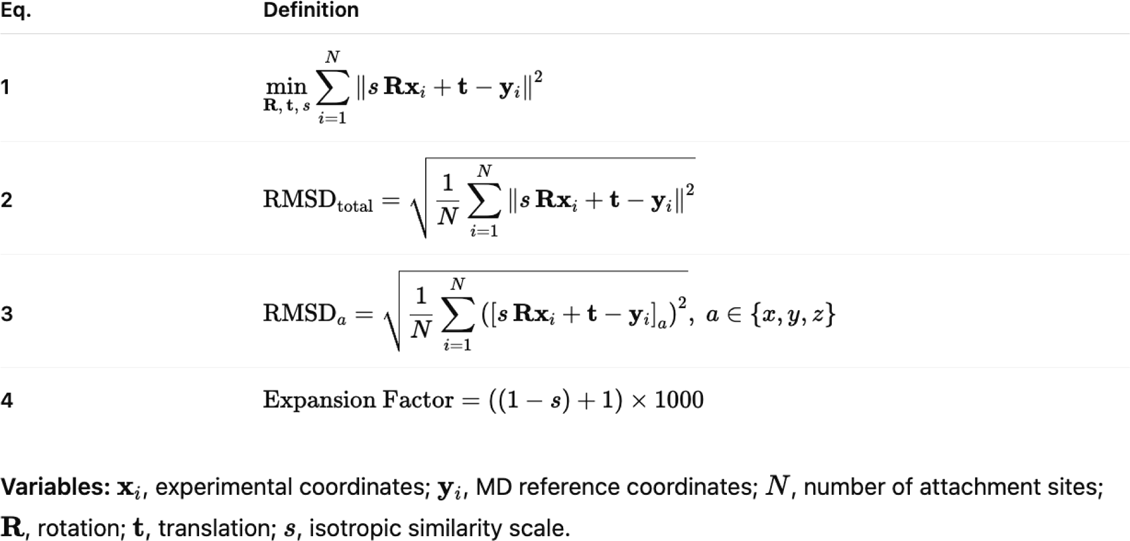

##### mCLING molecular dynamics simulations

The molecular dynamics (MD) simulations for mCLING-Atto647N were carried out using the Gromacs software package [48]. Functional groups were protonated to neutral pH to reflect buffer conditions. To accommodate the nonstandard Atto647N moiety, custom force field parameters were computed with Espaloma [49]. The starting configuration was obtained via RDKIT ETKDGv2 [50] and energy minimization in the MMFF94s force field. The molecule was solvated in a 6 nm box of TIP3P water with periodic boundaries and the system was neutralized with sodium chloride. Simulations were warmed up with a standard 3-stage equilibration protocol [51]. First, the energy of the system was first minimized via steepest descent for 50000 steps. Then, the system was equilibrated in the NVT ensemble for 200 ps with a step size of 2 ps and temperature of 300K with a modified Berendsen thermostat, and then in the NPT ensemble for 1 ns at 1 bar with a Parrinello-Rahman barostat. Production simulations were carried out in the NPT ensemble for 1 us, with one configuration saved every 10 ps. To enable larger step sizes and thus increase the simulation throughput, bonds to hydrogen atoms were constrained with the LINCS algorithm and nonbonded interactions were calculated with Particle Mesh Ewald and a 1 nm cutoff. Nonwater heavy atoms were then extracted for analysis.

##### GFP 3D reconstruction

The *ab initio* 3D protein reconstructions from 2D ONE images were performed as previously described <u>[5]</u>. Briefly, the pipeline implements an unsupervised amortized-inference framework that transforms preprocessed 2D ONE images into a coherent 3D molecular density. Preprocessing includes intensity normalization and background suppression to correct illumination variability and isolate relevant structural content. Local contrast enhancement is applied to emphasize edges and fine-scale features, followed by controlled smoothing to reduce high-frequency noise while preserving biologically meaningful boundaries. An enrichment step based on total variation regularization further stabilizes the projections by suppressing artifacts and enhancing structural continuity. The transformed projections are integrated and denoised to remove residual background fluctuations while maintaining sharp transitions, yielding a robust 2D representation suitable for inference.

Processed images are then input into a variational autoencoder (VAE) comprising convolutional layers with batch normalization, dropout, and nonlinear activations. The encoder maps each image into a latent representation that captures pose parameters and structural variability under constraints promoting smoothness, symmetry, and statistical consistency. The decoder reconstructs denoised projections using a positional multi-layer perceptron. The volume generator is evaluated slice-by-slice across a grid of frequency coordinates to build a 3D Fourier volume. An inverse Fourier transform then produces a volumetric density map that represents a physically plausible 3D molecular structure obtained without imposing predefined atomic coordinates or using external templates or supervision, see **Supplementary Figures 24–28**. The FSC threshold 0.143 has been considered for final resolution calculation.

## Supplementary Figures

**Supplementary Figure 1.**
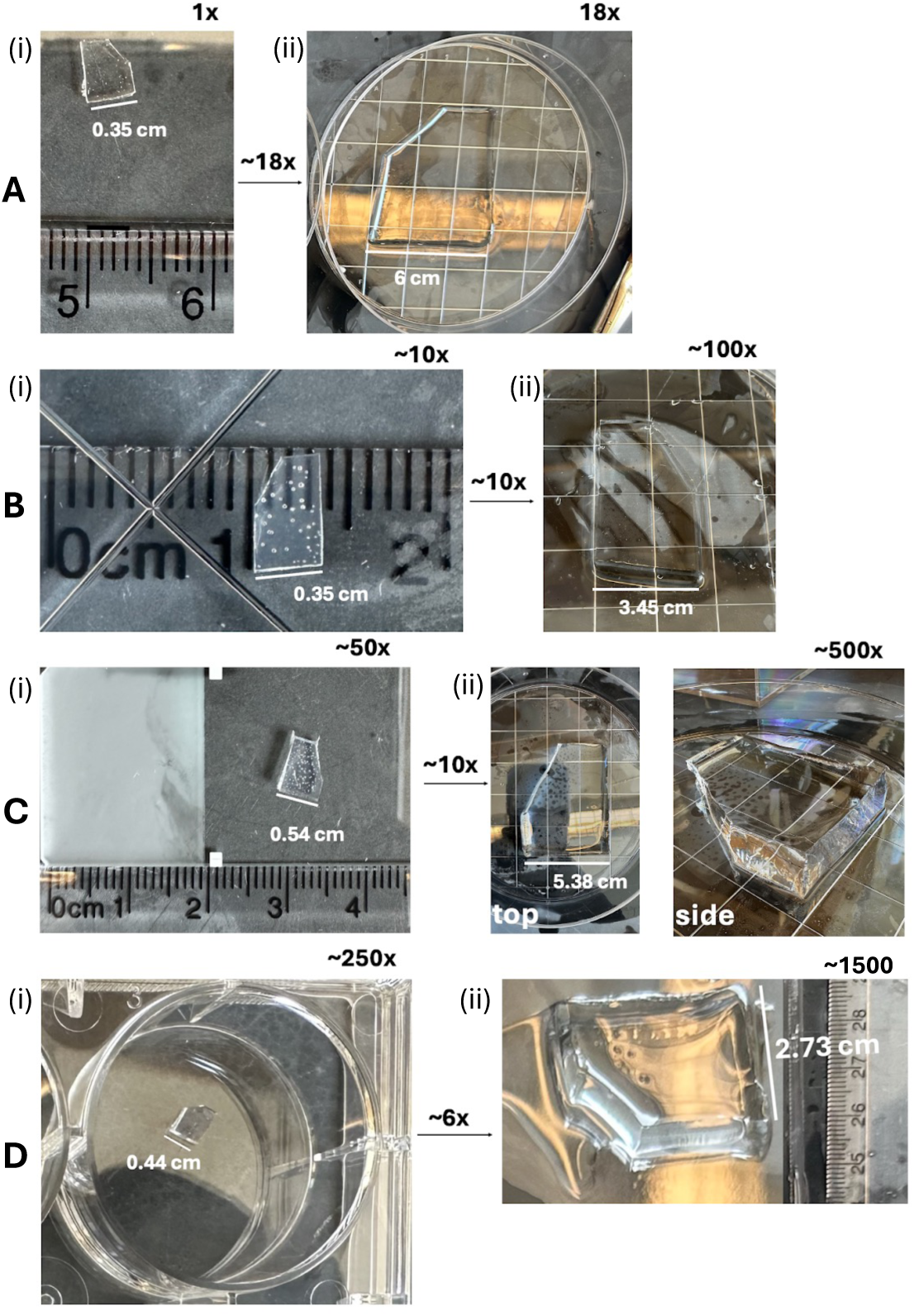
Representative hydrogels at successive expansion stages. The hydrogel consists of four interpenetrating polymer networks generated by sequential monomer infiltration, polymerization under deoxygenated conditions (>8 h, 20 °C), and re-expansion in water. Linear expansion was determined by measuring one gel edge with a ruler (minimum spacing 0.1 cm). (A) First network (∼18×). The primary swellable gel was cast from the monomer solution (Fig. 1A). At (i) 1×, the gel is colorless and transparent with white specks consistent with trapped air. After expansion in water to (ii) ∼18×, the gel is transparent with straight edges and flat surfaces. (B) Second network (∼100× total). The fully expanded ∼18× gel was infiltrated with a charged monomer solution and polymerized, yielding a composite gel that contracts to (i) ∼10× linear expansion. At this stage, the gel is translucent with spherical bubbles. Re-expansion to (ii) ∼100× (∼10× from i) produces a transparent gel with straight edges and flat surfaces that stands upright without visible axial bending. (C) Third network (∼500× total). The fully expanded ∼100× gel was infiltrated with a charged monomer solution and polymerized, yielding a composite gel that contracts to (i) ∼50× linear expansion. The gel is translucent with spherical bubbles. Re-expansion to (ii) ∼500× (∼10× from i) produces a transparent gel with straight edges and flat surfaces that stands upright without visible axial bending. (D) Fourth network (∼1500× total). The fully expanded ∼500× gel was infiltrated with a charged monomer solution and polymerized, yielding a composite gel that contracts to (i) ∼250× linear expansion. Re-expansion to (ii) ∼1500× (∼6× from i) produces a transparent gel with slightly inward-curved (concave) surfaces, indicating resistance to further swelling, perhaps imposed by the covalently crosslinked network. (This did not seem to be gravity-induced: the gel remained rigid and self-supporting, with the slight inward curvature reflecting elastic constraint opposing further swelling, for this specific gel condition.) Upon equilibration in 1× PBS for imaging, the gel contracts to ∼1000× linear expansion, and the slight concavity goes away. Furthermore, with optimization of the gelling process (**Supp. Fig. 31**), round 4 gels expand without any perceptible concavity.

**Supplementary Figure 2.**
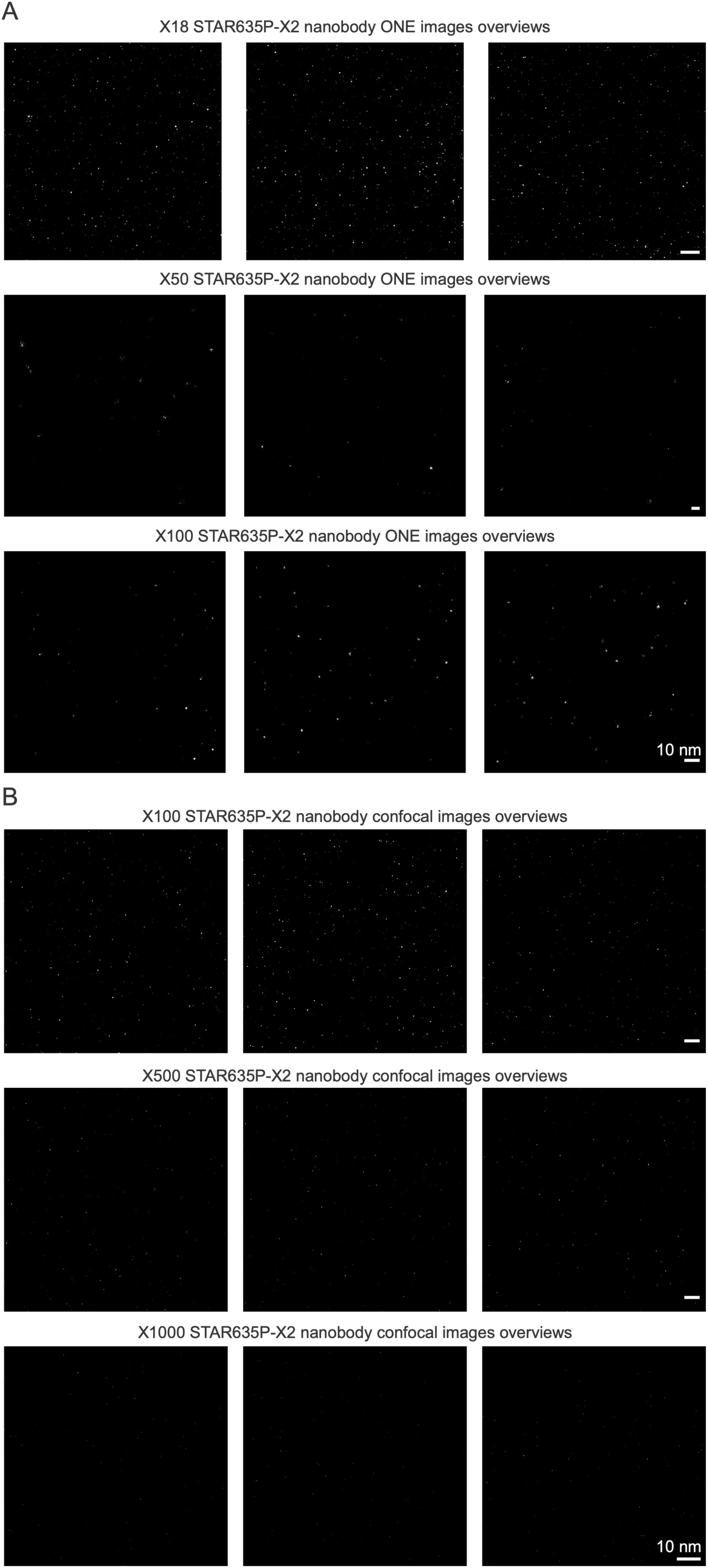
Nanobody imaging across different linear expansion factors. (A) ONE microscopy at ∼18×, ∼50×, and ∼100× expansion, of representative fields of view from representative gels. (B) Confocal imaging at ∼100×, ∼500×, and ∼1000× expansion. Representative datasets are shown, from 2-7 experimental replicates.

**Supplementary Figure 3.**
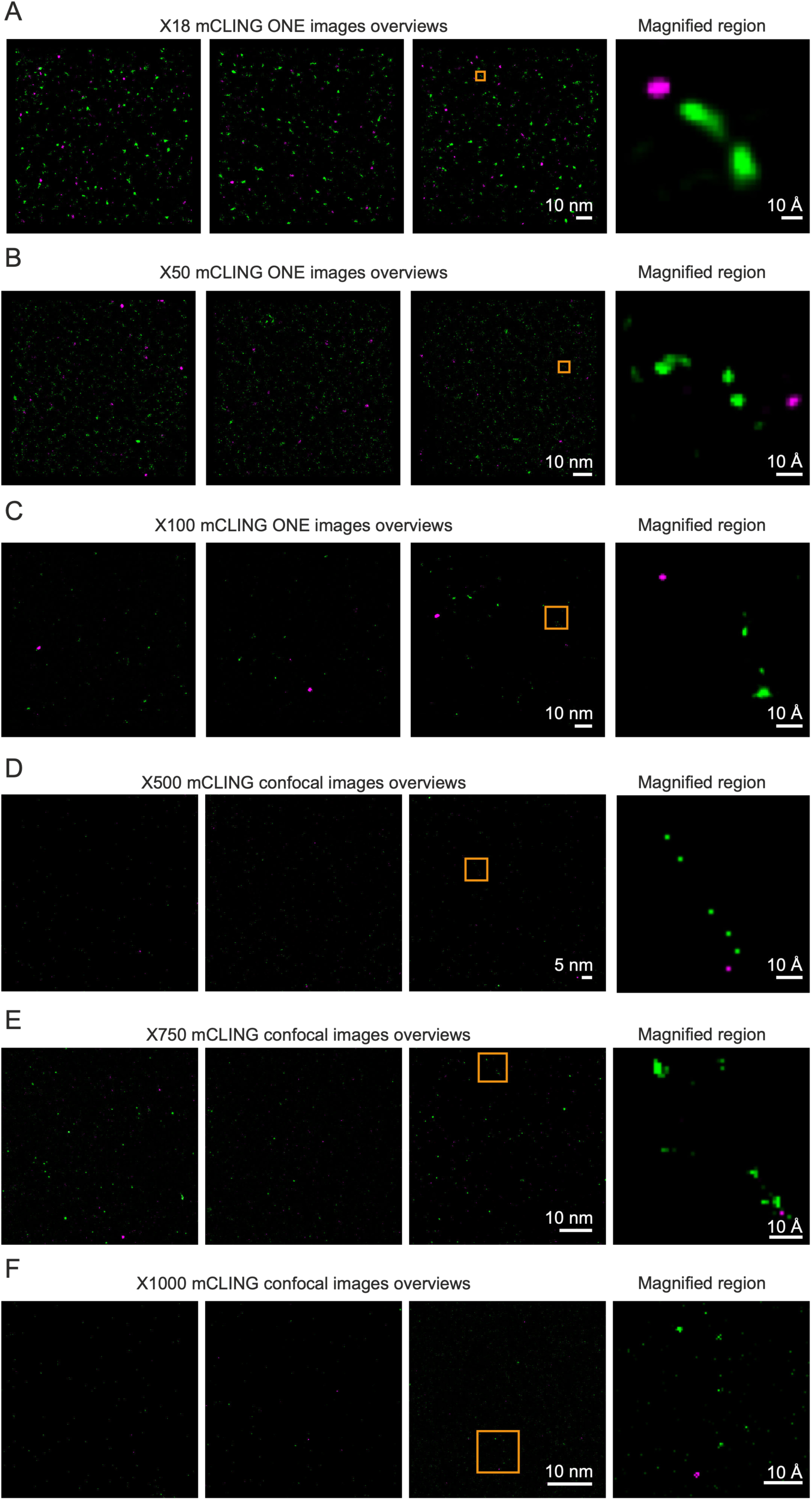
mCLING peptide imaging across different linear expansion factors. (A–C) ONE microscopy at ∼18×, ∼50×, and ∼100× expansion. (D–F) Confocal imaging at ∼500×, ∼750×, and ∼1000× expansion.

**Supplementary Figure 4.**
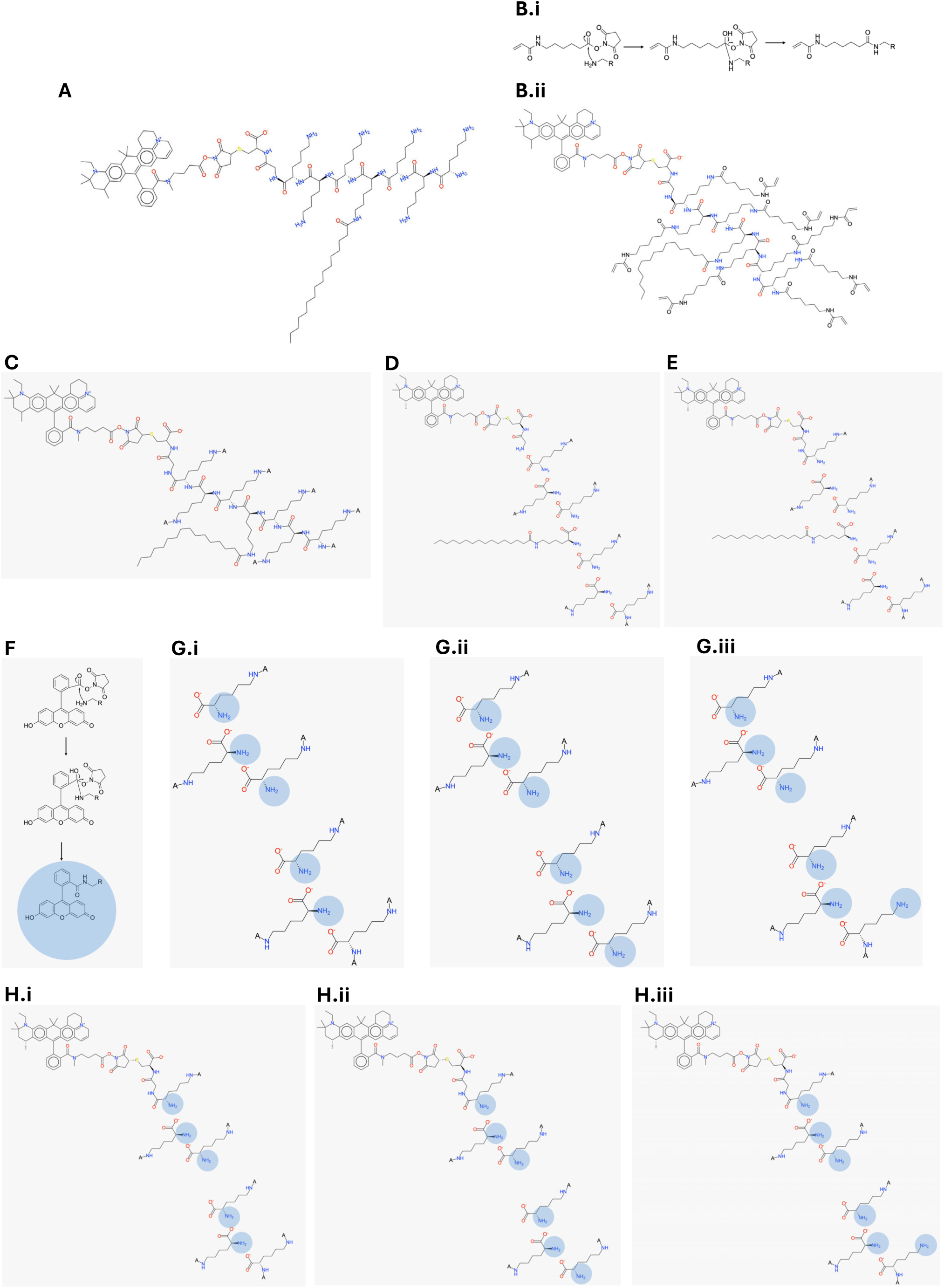
Pan-lysine anchoring, proteolytic cleavage, and post-expansion labeling illustrated using the mCLING peptide. Gray boxes indicate reactions performed inside the expansion hydrogel; all other steps were performed in solution. A. Chemical structure of the mCLING peptide. B.i NHS-acrylate (Acryloyl-X) reacts with primary amines via NHS ester–amine coupling. B.ii All primary amines on the mCLING peptide, ideally, are functionalized with polymerizable acrylate groups. C. Hydrogel polymerization (gray box). Acrylate-functionalized amines (abbreviated A) are covalently attached to the gel network, anchoring the peptide at the labeled sites. D. Proteinase K digestion in which the amide bond between the anchoring site and Atto647N is cleaved. All peptide backbone amide bonds are digested. Fragments lacking an anchoring site are removed during expansion. Because Atto647N is not anchored, that dye is lost. E. Proteinase K digestion in which the amide bond between the anchoring site and Atto647N is not cleaved. Atto647N remains attached to an anchored fragment and is therefore retained and displaced during expansion according to the anchoring atom. Cleavage of this bond is unlikely due to steric constraints of the Atto647N substituent in the Proteinase K substrate recognition site [53]. F. NHS-ester fluorescein labeling of remaining free primary amines. The blue sphere denotes a primary amine that has been conjugated with fluorescein. In this panel, the full chemical conjugation product is shown. G. I–III. Labeling outcomes for case D (Atto647N cleaved and removed). In G and H, the blue sphere is used as a symbolic shorthand indicating an NH₂ group conjugated with fluorescein. G.I Both the N-terminal amine and the lysine side-chain amine are anchored. No free primary amines remain; no fluorescein labeling occurs. G.II Only the lysine side-chain amine is anchored. The free N-terminal amine is fluorescein-labeled (blue sphere). Upon expansion, the dye is displaced to the position of the lysine anchoring atom. G.III Only the N-terminal amine is anchored. The lysine side-chain amine is fluorescein-labeled (blue sphere). The dye is displaced to the position of the N-terminal anchoring atom. H. I–III. Labeling outcomes for case E (Atto647N retained). Configurations are the same as G.I–III, except that Atto647N remains attached and is displaced according to its anchoring site.

**Supplementary Figure 5.**
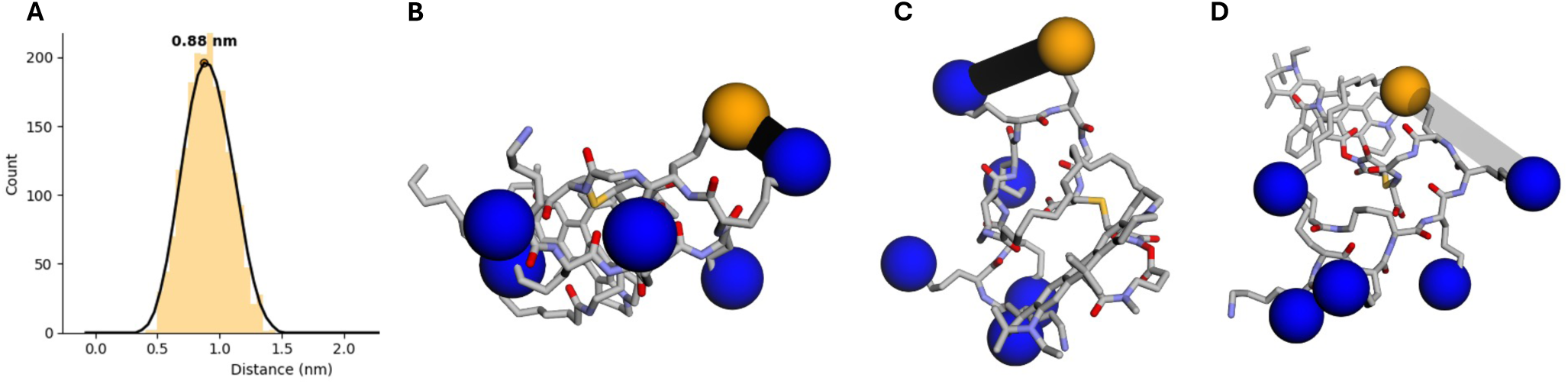
Distance distribution between adjacent lysine side chains. Primary amines were functionalized with NHS–ester acrylate to covalently anchor the nitrogen atom to the polymer network. In the schematic, the nitrogen of the first primary amine bearing the Atto647N dye is shown as an orange sphere; adjacent lysine ε-amine nitrogens are shown as blue spheres. These nitrogen atoms are attached to the hydrogel and displaced during expansion. Thus, the measured positions of Atto647N and fluorescein correspond to the displaced coordinates of their respective anchoring nitrogens. Inter-lysine distances were quantified by molecular dynamics. (A) Distribution of distances (nm). (B) Minimum distance (∼0.44 nm), parallel side chains. (C) Peak distance (∼0.88 nm), staggered orientation. (D) Larger distance (∼1.41 nm), antiparallel orientation.

**Supplementary Figure 6.**
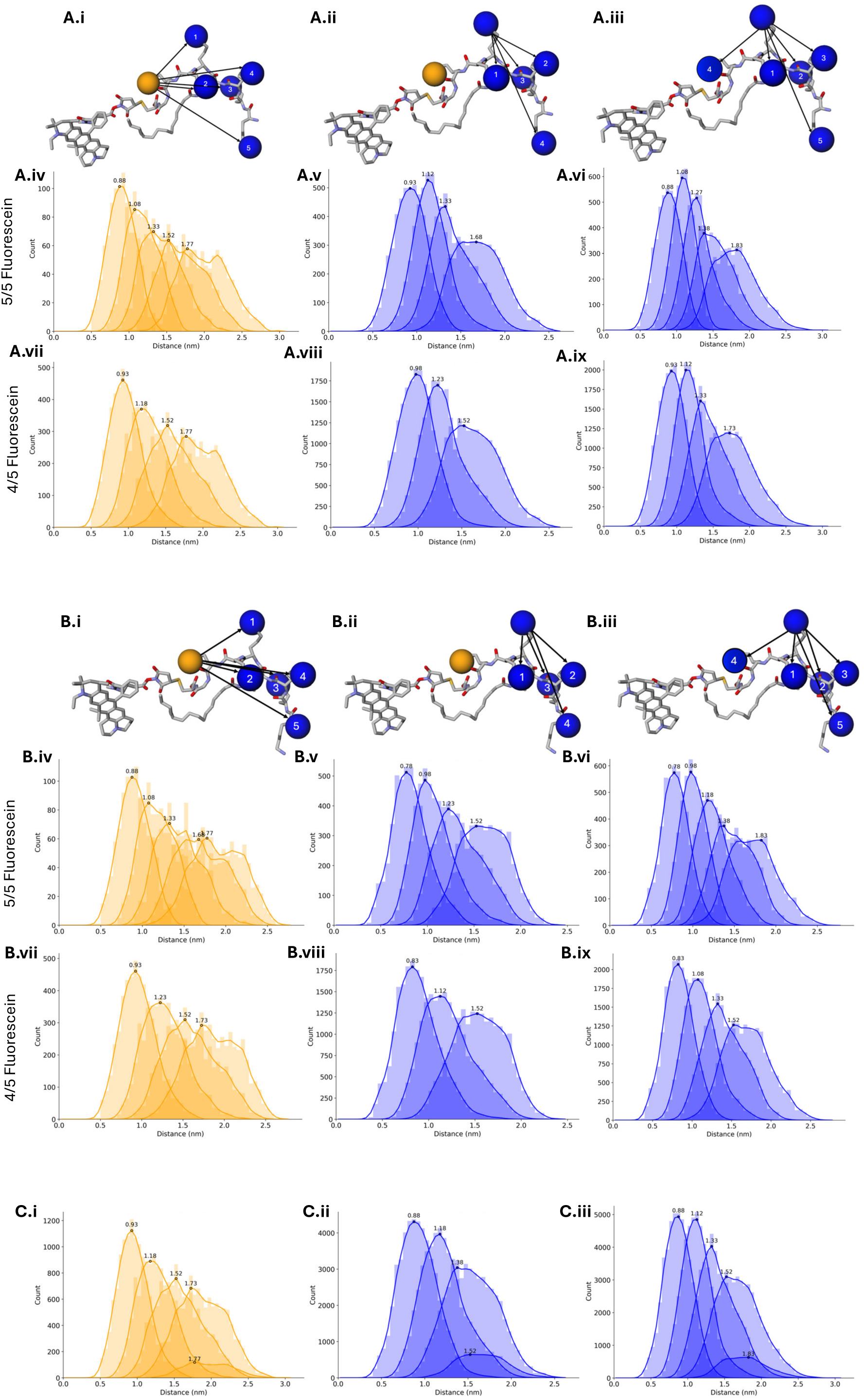
Molecular dynamics (MD) simulation of k-nearest-neighbor (kNN) distance distributions for the mCLING peptide followed by analysis under defined anchoring, cleavage, and labeling scenarios. Molecular dynamics (MD) simulations generated 1001 conformations of the mCLING peptide (see Methods: mCLING molecular dynamics simulations). All kNN distance distributions shown were computed across the full ensemble of 1001 conformations. Atomic model panels display the first conformation. For each conformation, fluorophore positions were assigned to the atomic coordinates of gel-anchoring atoms defined in **Supplementary Figure 4**. Orange spheres denote Atto647N anchoring sites and blue spheres denote fluorescein anchoring sites. k-nearest-neighbor (kNN) distances were computed as Euclidean distances between anchoring-site atomic coordinates. In atomic model panels, arrows originate from a selected anchoring-site atom and point to its k-nearest neighboring sites; neighboring sites are numbered by increasing Euclidean distance, with labels 1–5 or 1–4 indicating the kNN rank for the representative conformation. All quantitative distributions were computed using all 1001 conformations, with one distance per kNN rank per conformation. Two terminal fluorescein labeling scenarios defined in **Supplementary Figure 4** were simulated: (A) terminal fluorescein assigned to a lysine side-chain amine (**Supplementary Fig. 4G.ii and 4H.ii**), and (B) terminal fluorescein assigned to the N-terminal amine (**Supplementary Fig. 4G.iii and 4H.iii**). In **Supplementary Figure 4**, panel G denotes configurations in which Atto647N is cleaved and removed, whereas panel H denotes configurations in which Atto647N is retained. (C) shows pooled distance distributions combining both terminal fluorescein labeling scenarios. (A) Terminal fluorescein assigned to a lysine side-chain amine. (i) Representative atomic model illustrating the anchoring configuration corresponding to **Supplementary Fig. 4H.ii**, in which Atto647N is retained and the terminal fluorescein is assigned to a lysine side-chain amine. Arrows originate from the Atto647N anchoring-site atom and point to its k-nearest fluorescein neighbors (kNN ranks 1–5). (ii) Same model as (i), with arrows originating from each fluorescein anchoring-site atom in turn and pointing to its nearest fluorescein neighbors (kNN ranks 1–4). (iii) Representative atomic model for the corresponding Atto-cleaved configuration (**Supplementary Fig. 4G.ii**), in which Atto647N is removed and the newly generated N-terminus is labeled with fluorescein (kNN ranks 1–5). (iv) Atto647N→fluorescein kNN distance distributions (ranks 1–5) computed from the configuration in (i) across all 1001 conformations, with the Atto647N anchoring site queried against fluorescein-labeled sites (one distance per kNN rank per conformation). (v) Fluorescein→fluorescein kNN distance distributions (ranks 1–4) computed from the same anchoring configuration as in (iv), using only fluorescein-labeled sites. (vi) Fluorescein→fluorescein kNN distance distributions (ranks 1–5) computed from the same anchoring geometry as in (iv), after treating the Atto647N anchoring-site coordinate as a fluorescein site for analysis. This corresponds to the Atto-cleaved case in which the newly generated N-terminus is labeled with fluorescein while the anchoring atom remains unchanged, such that the underlying anchoring-site coordinates are identical. (vii–ix) Five-choose-four subsampling was used to reflect molecule-to-molecule variation in which anchoring and labeling sites are present after anchoring, proteolysis, and post-expansion labeling—including outcomes such as **Supplementary Fig. 4G.i and 4H.i**, in which both the N-terminal amine and a lysine side-chain amine are anchored. All subsets of four labeling-site coordinates were selected from the five available sites for each peptide conformation. (vii) Atto647N→fluorescein kNN distance distributions computed from five-choose-four subsampled fluorescein site sets in Atto-retained configurations. (viii) Fluorescein→fluorescein kNN distance distributions computed from the same five-choose-four subsampled fluorescein site sets as in (vii). (ix) Fluorescein→fluorescein kNN distance distributions computed using five-choose-four subsampling in Atto-cleaved configurations, with the former Atto647N anchoring-site coordinate treated as a fluorescein site for analysis. (B) Terminal fluorescein assigned to the N-terminal amine. (i–ix) Same anchoring configurations, kNN definitions, and five-choose-four subsampling procedures described in (A)(i–ix), applied to configurations in which the terminal fluorescein is assigned to the N-terminal amine (**Supplementary Fig. 4G.iii and 4H.iii**). (C) Combined distance distributions. (i) Combined Atto647N→fluorescein kNN distance distributions pooled across both terminal fluorescein labeling scenarios for configurations in which Atto647N is retained (**Supplementary Fig. 4H**). (ii) Combined fluorescein→fluorescein kNN distance distributions pooled across both terminal fluorescein labeling scenarios for configurations in which Atto647N is retained (**Supplementary Fig. 4H**). (iii) Combined fluorescein→fluorescein kNN distance distributions pooled across both terminal fluorescein labeling scenarios for configurations in which Atto647N is cleaved (**Supplementary Fig. 4G**).

**Supplementary Figure 7.**
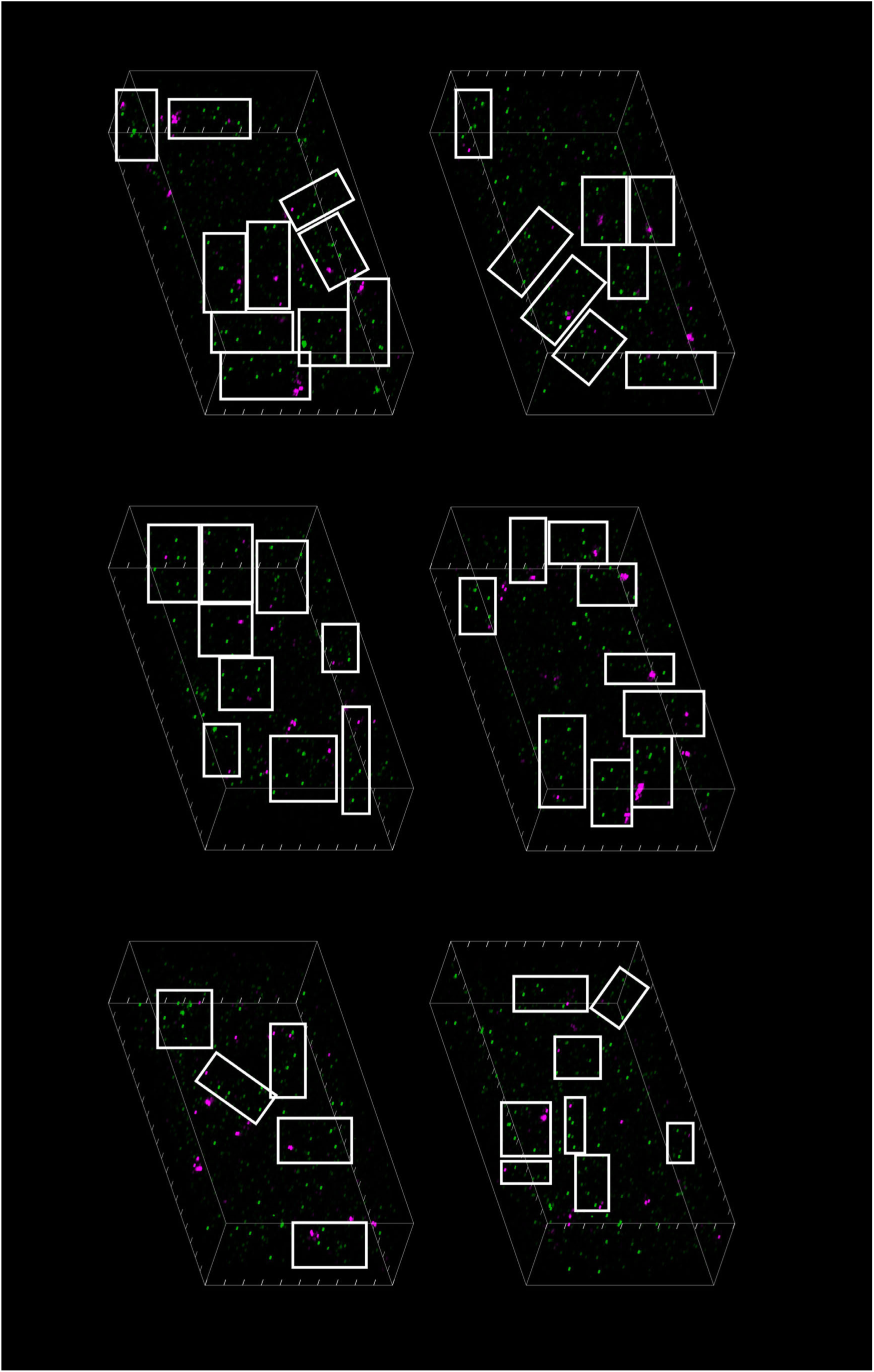
3D regions containing putative mCLING peptides after ∼1000× expansion. Three-dimensional renderings of six regions selected to be representative with respect to signal brightness within a 10 × 20 µm field spanning the full z depth (4.4 µm) after ∼1000× linear expansion. Boxed regions highlight examples of putative mCLING peptides, defined as kNN-derived clusters containing one Atto647N signal (magenta) and at least four proximal fluorescein signals (green). Boxes are shown for illustrative examples and do not mark all such clusters within the field. Because mCLING peptides contain a hydrophobic palmitoyl tail, intermolecular interactions may occur, and some boxed regions may contain more than one putative peptide due to spatial overlap; in cases of overlap, one box was randomly selected.

**Supplementary Figure 8.**
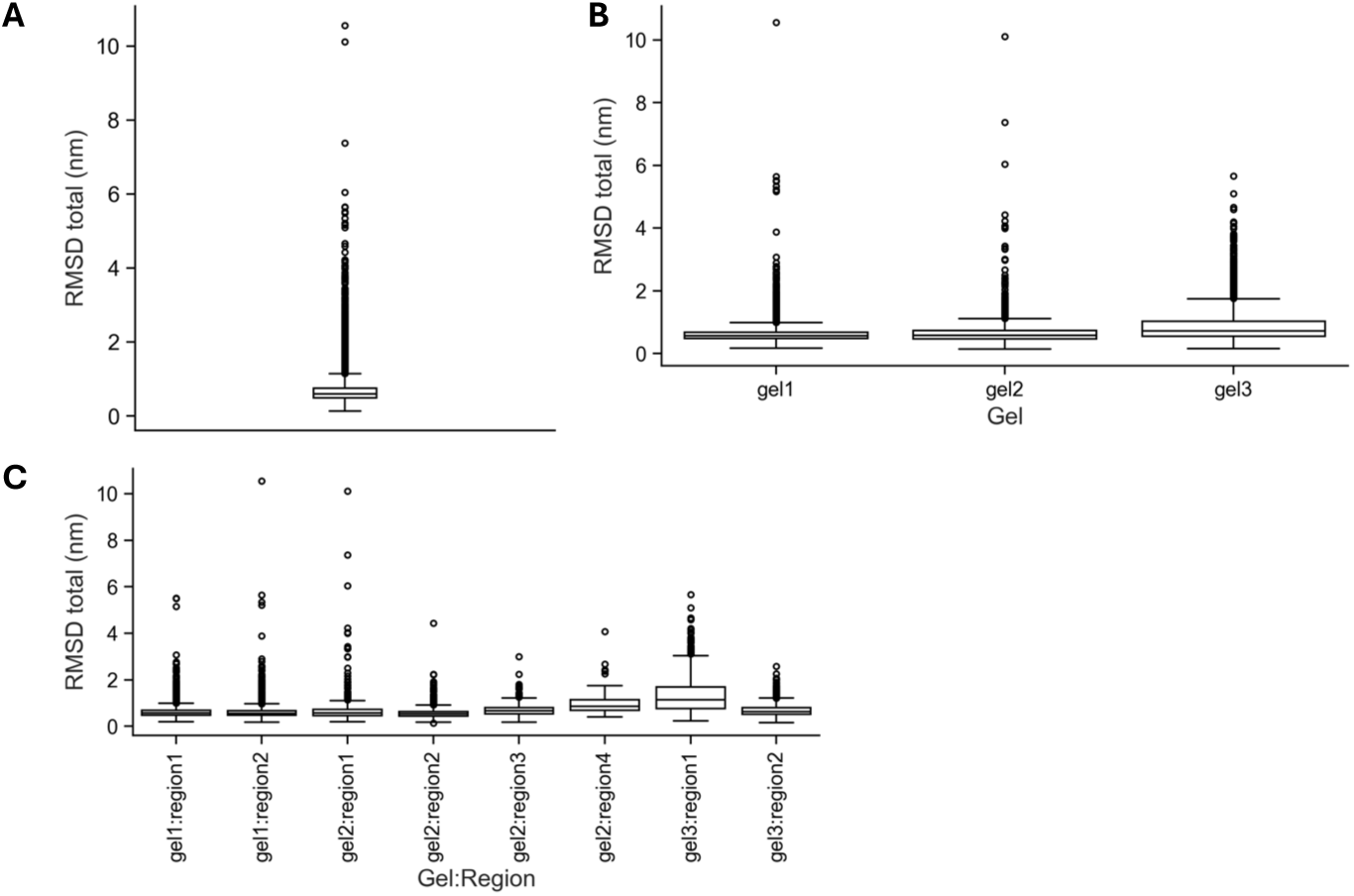
RMSD of coarse putative peptides before nearest-neighbor filtering. From 27,426 Atto647N localizations (three gels, eight regions), fluorescein signals were associated with the nearest Atto647N, yielding 10,696 coarse putative peptides (see Methods, Identification of putative peptides). Each contains one Atto647N and four or five fluoresceins; assignments may include signals from neighboring peptides. For each coarse putative peptide, observed coordinates were aligned to a 1,001-conformation molecular-dynamics ensemble spanning anchoring, digestion, and labeling outcomes (**Supplementary Figs. 4, 6**), and RMSD was computed. Boxplots show the median, IQR, whiskers extending to the farthest data point within 1.5×IQR of each box edge, and outliers. (A) All gels. (B) Individual gels. (C) Individual regions.

**Supplementary Figure 9.**
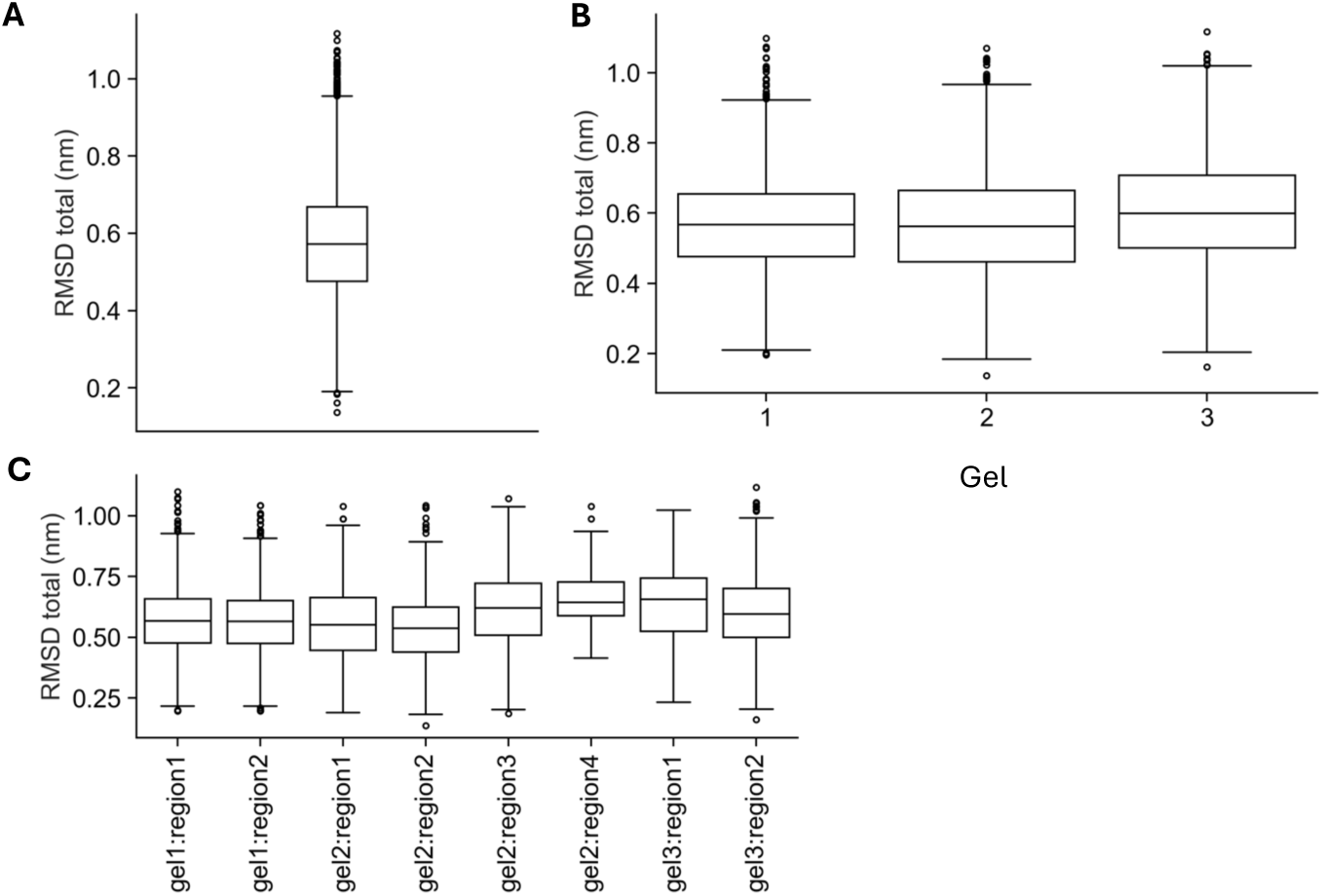
RMSD of putative peptides after nearest-neighbor filtering. Nearest-neighbor filtering (see Methods; filtered subset of **Supplementary Fig. 8**; same peptide set as Fig. 3C) yields the final putative peptides. RMSD was computed by alignment to the molecular-dynamics ensemble and shown as boxplots. (A) All gels. (B) By gel. (C) By region. Summary statistics: Table S13 (A), Table S14 (B), Table S15 (C).

**Supplementary Figure 10.**
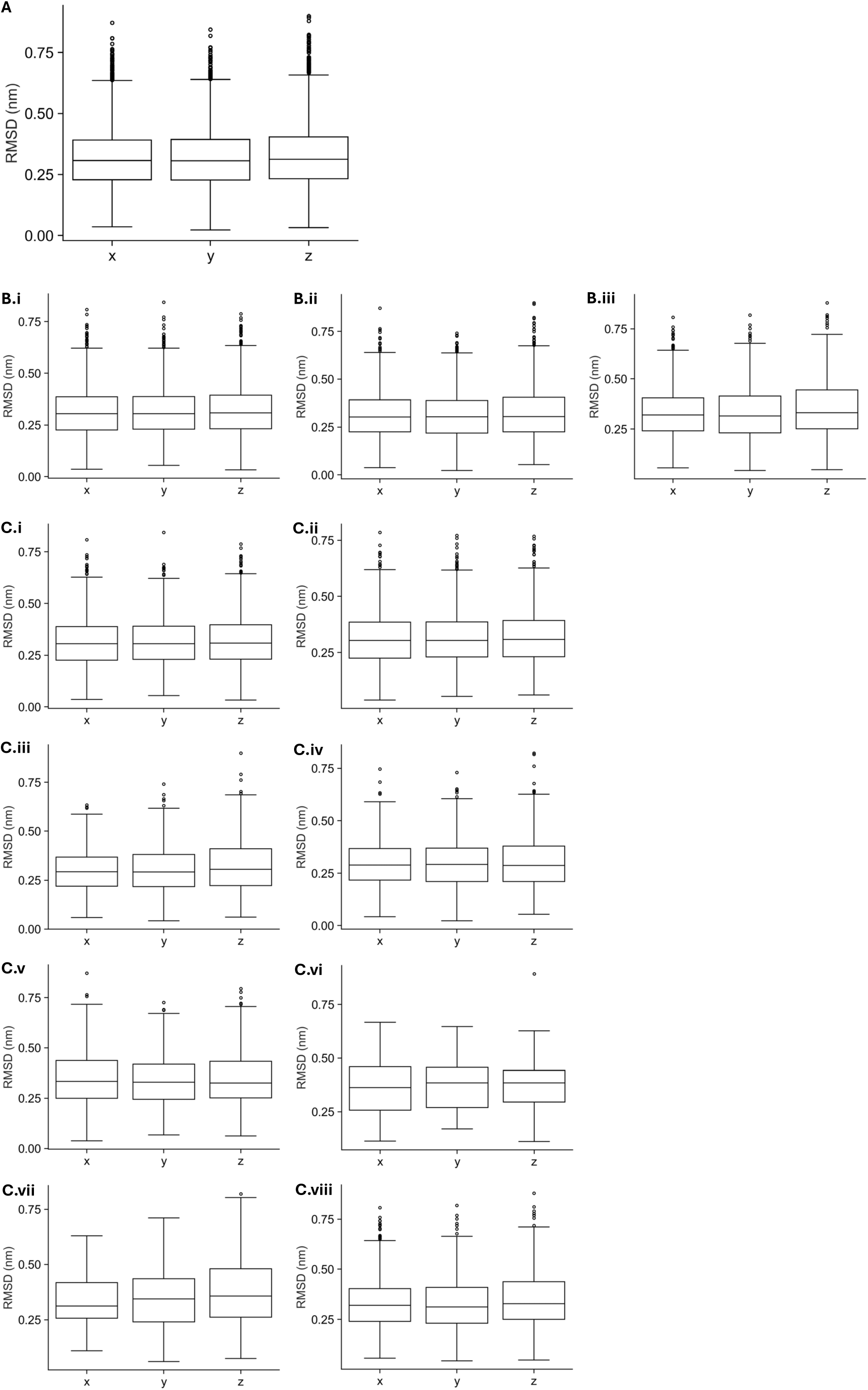
Per-axis RMSD of putative peptides (same peptide set as Fig. 3C). Per-axis RMSD (x, y, z) was computed using the same analysis as Supplementary Fig. 9. Fig. 3C shows pooled data; here the same data are shown pooled, by gel, and by region to assess variation. Summary statistics: Table S13 (A), Table S14 (B), Table S15 (C).

**Supplementary Figure 11.**
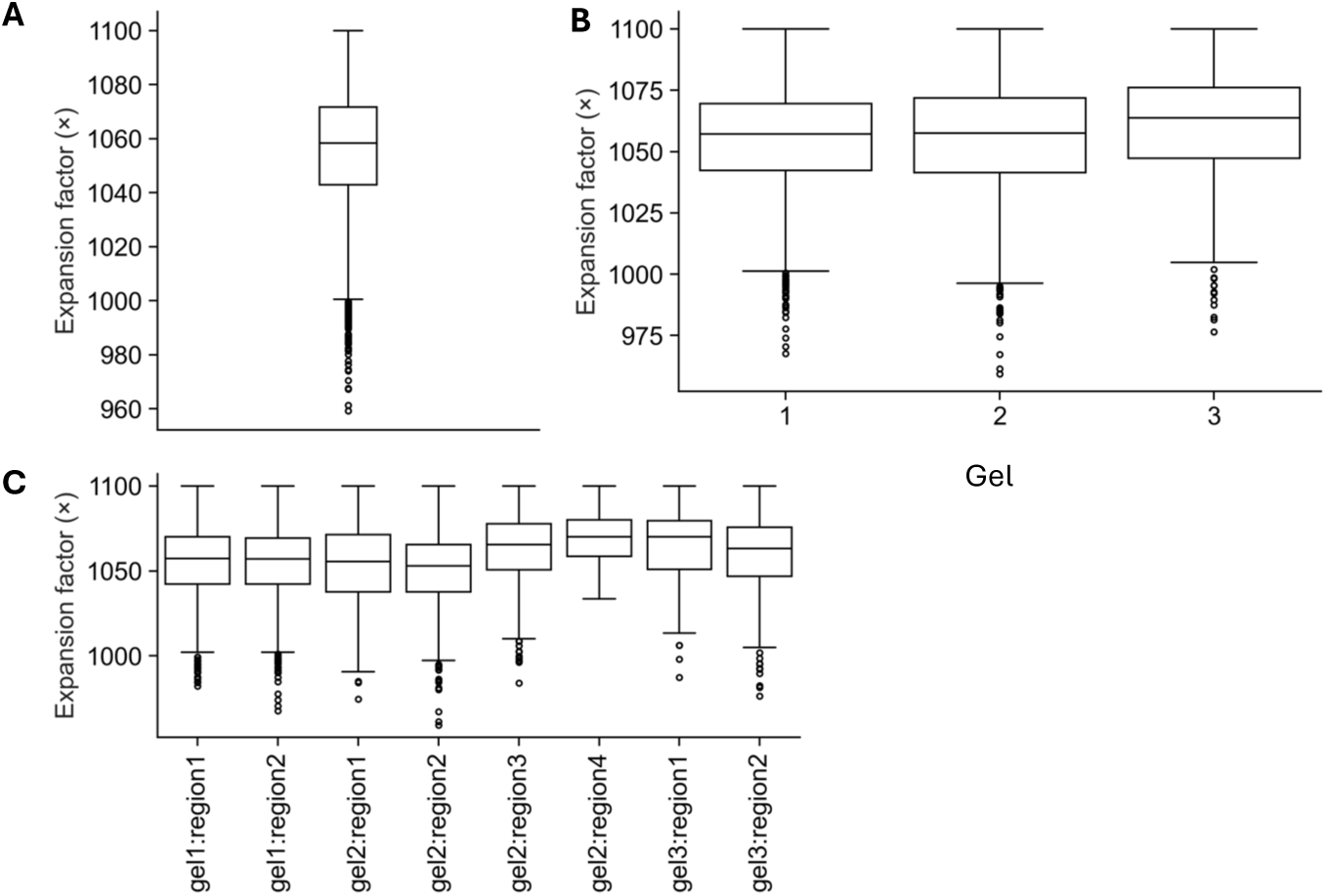
Single-molecule expansion factors (same peptide set as Fig. 3C). An isotropic expansion factor was estimated for each peptide by minimizing RMSD to the molecular-dynamics ensemble, using the same analysis as Supplementary Fig. 9. Fig. 3C shows pooled data; here the same data are shown pooled, by gel, and by region to assess variation. (A) All gels combined. (B) By gel. (C) By region. Summary statistics: Table S13 (A), Table S14 (B), Table S15 (C).

**Supplementary Figure 12.**
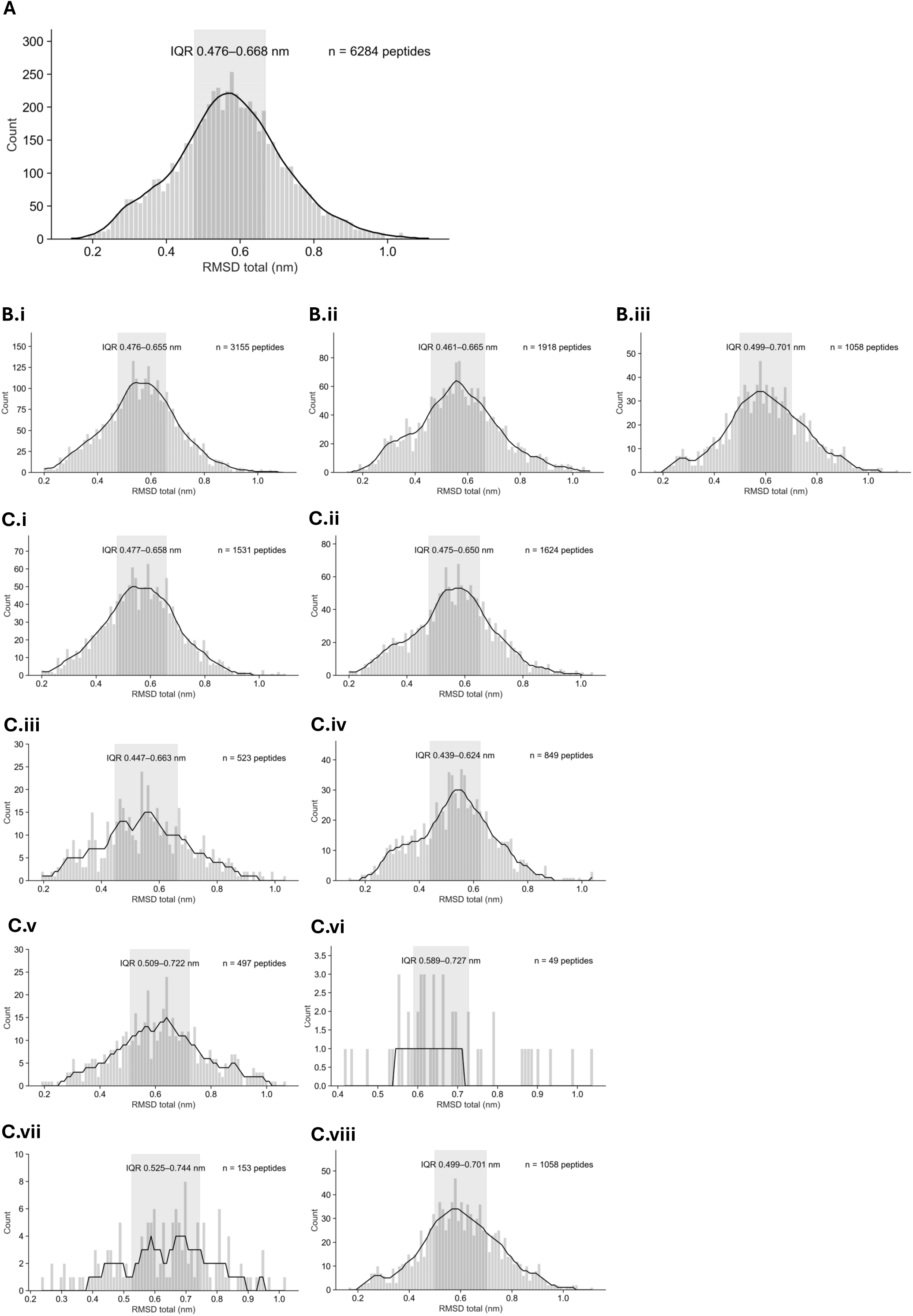
RMSD histograms (same peptide set and RMSD values as Fig. 3C). The same RMSD values are replotted as histograms with interquartile ranges; only the visualization differs. Fig. 3C shows pooled data; here the same data are shown pooled, by gel, and by region to assess variation. (A) All data combined. (B) By gel. (C) By region. Summary statistics: Table S13 (A), Table S14 (B), Table S15 (C).

**Supplementary Figure 13.**
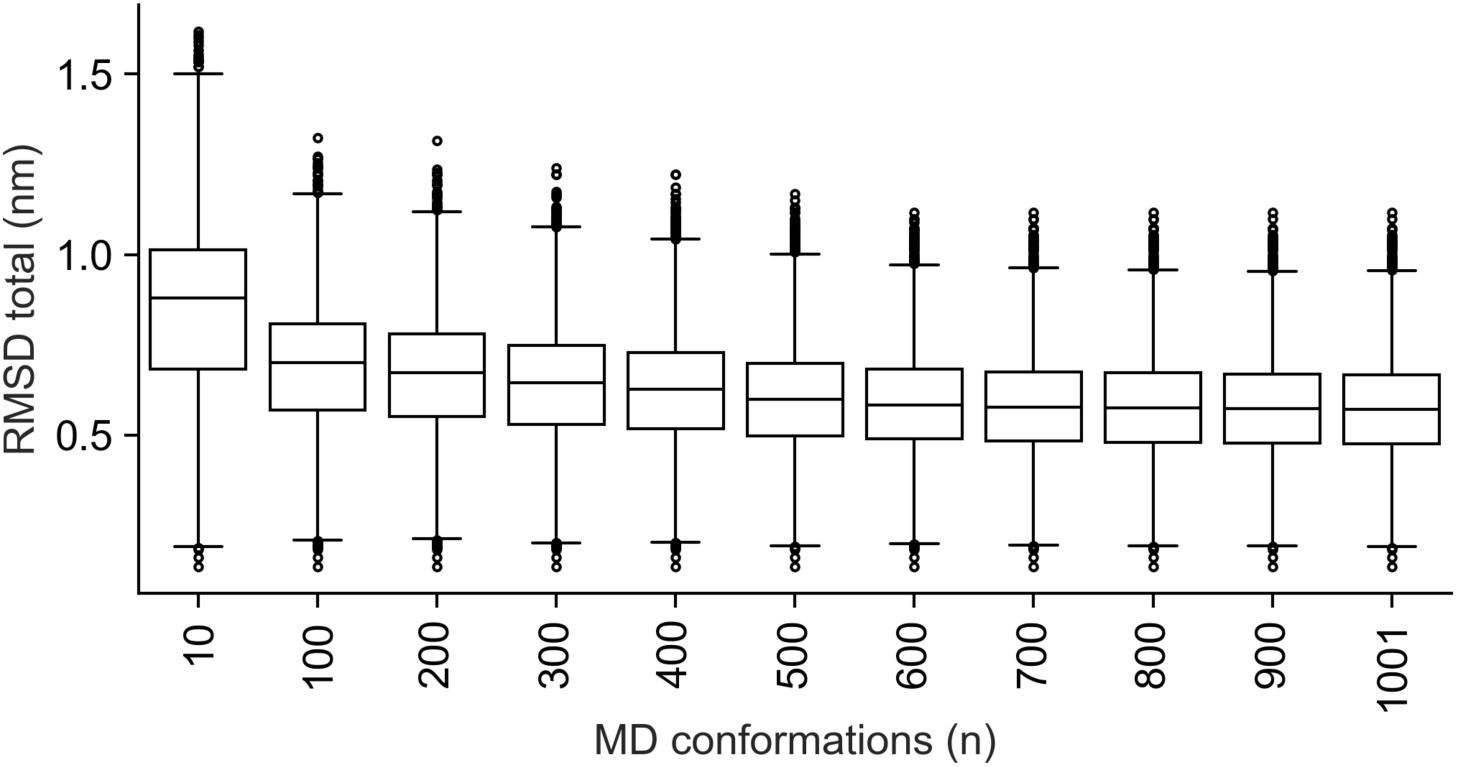
Robustness of RMSD to MD ensemble size. Using the filtered putative peptides defined in Supplementary Fig. 9, total RMSD was recomputed after alignment to MD reference ensembles containing 10–1,001 conformations. For each ensemble size, RMSD distributions are summarized as boxplots (median, IQR, whiskers to farthest data points within 1.5×IQR of box edges, outliers). Summary statistics are reported in **Table S16**.

**Supplementary Figure 14.**
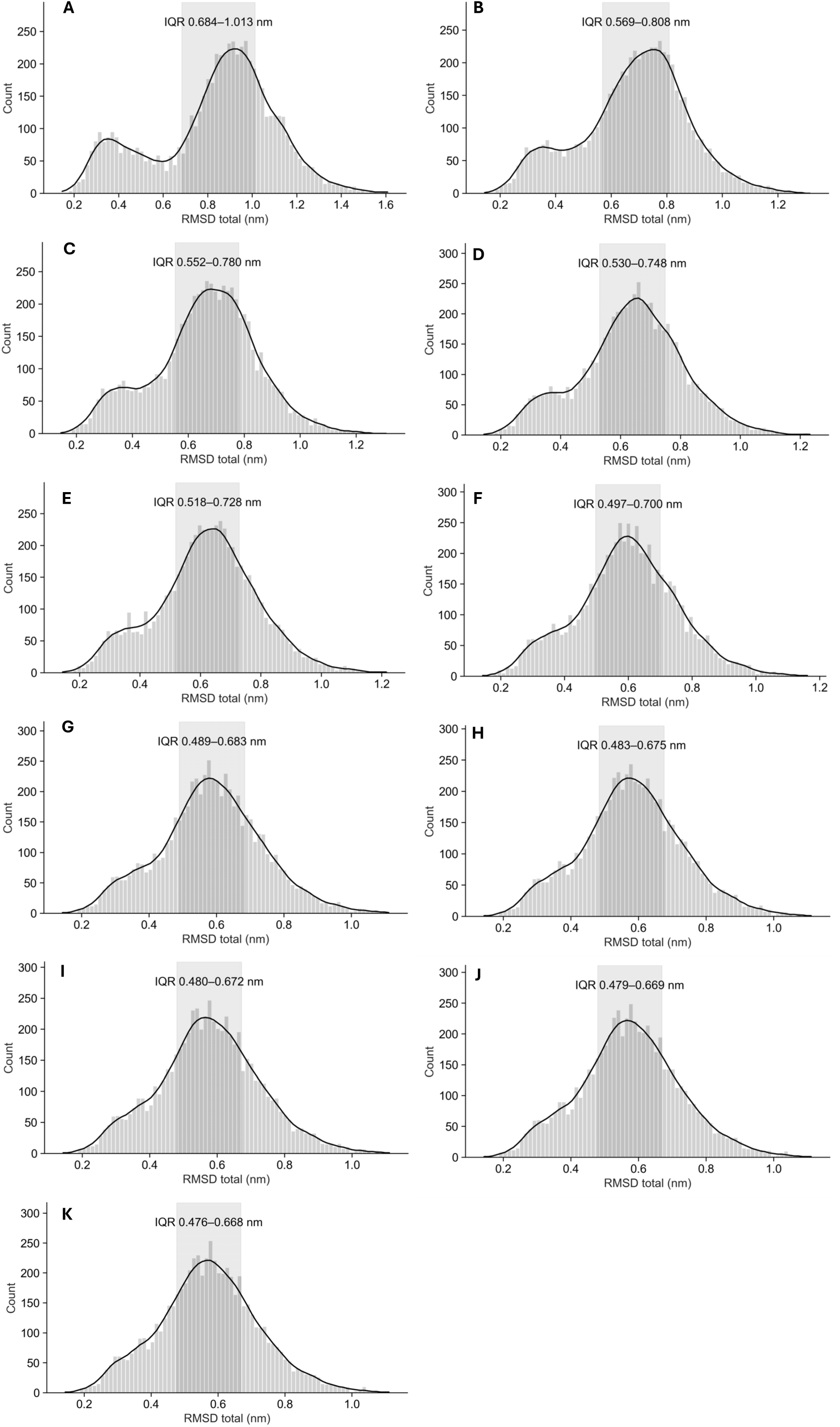
RMSD histograms across MD ensemble sizes. For the same peptides and RMSD values analyzed in Supplementary Fig. 13, distributions are replotted as histograms to visualize distribution shape as a function of MD ensemble size. Panels correspond to MD ensemble sizes: (A) 10; (B) 100; (C) 200; (D) 300; (E) 400; (F) 500; (G) 600; (H) 700; (I) 800; (J) 900; (K) 1,001 conformations. Summary statistics are reported in **Table S16.**

**Supplementary Figure 15.**
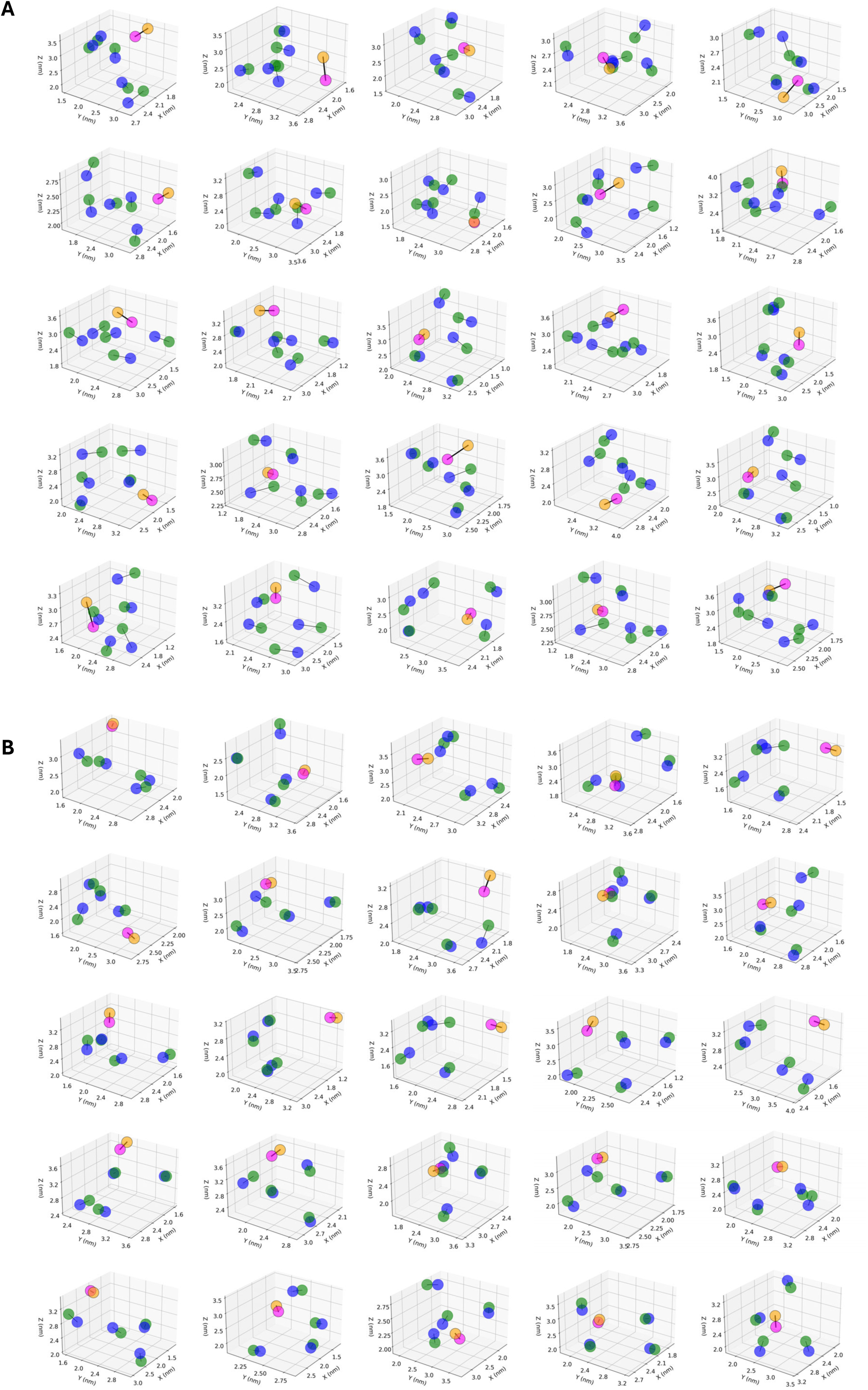
Peptide overlays below the IQR of the Fig. 3C.i RMSD distribution. Putative peptides with total RMSD below the lower quartile of Fig. 3C**.i** are shown. Fluorophore coordinates are overlaid after optimal alignment to the molecular-dynamics ensemble (**Supplementary Fig. 6**). Peptide length reflects anchoring, cleavage, and labeling outcomes (**Supplementary Fig. 4**), yielding four or five fluorescein sites. (A) 1 Atto647N followed by 5 fluoresceins. (B) 1 Atto647N followed by 4 fluoresceins.

**Supplementary Figure 16.**
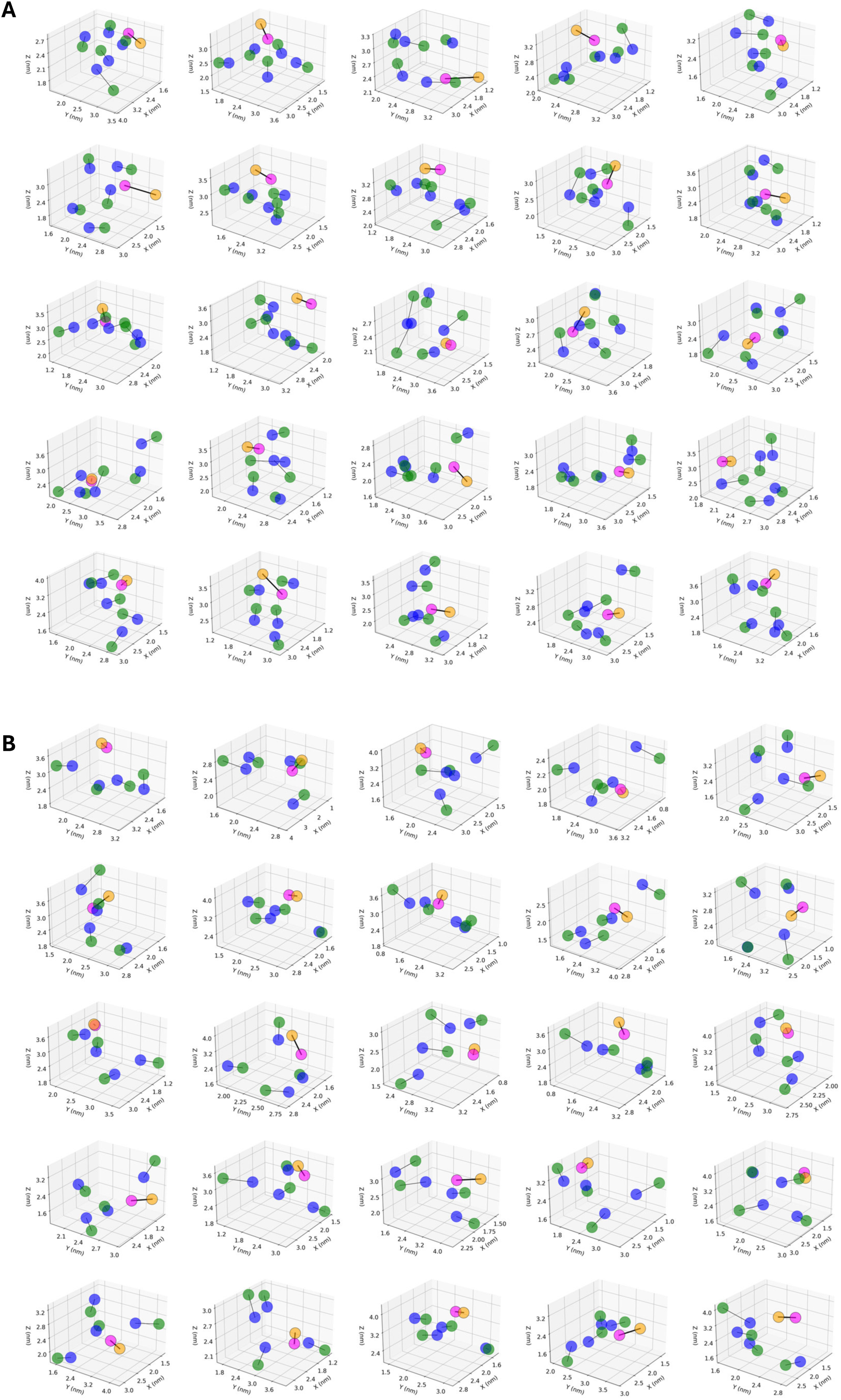
Peptide overlays within the IQR of the Fig. 3C.i RMSD distribution. Same analysis and representation as Supplementary Fig. 15, for peptides with RMSD within the interquartile range of Fig. 3C.i.

**Supplementary Figure 17.**
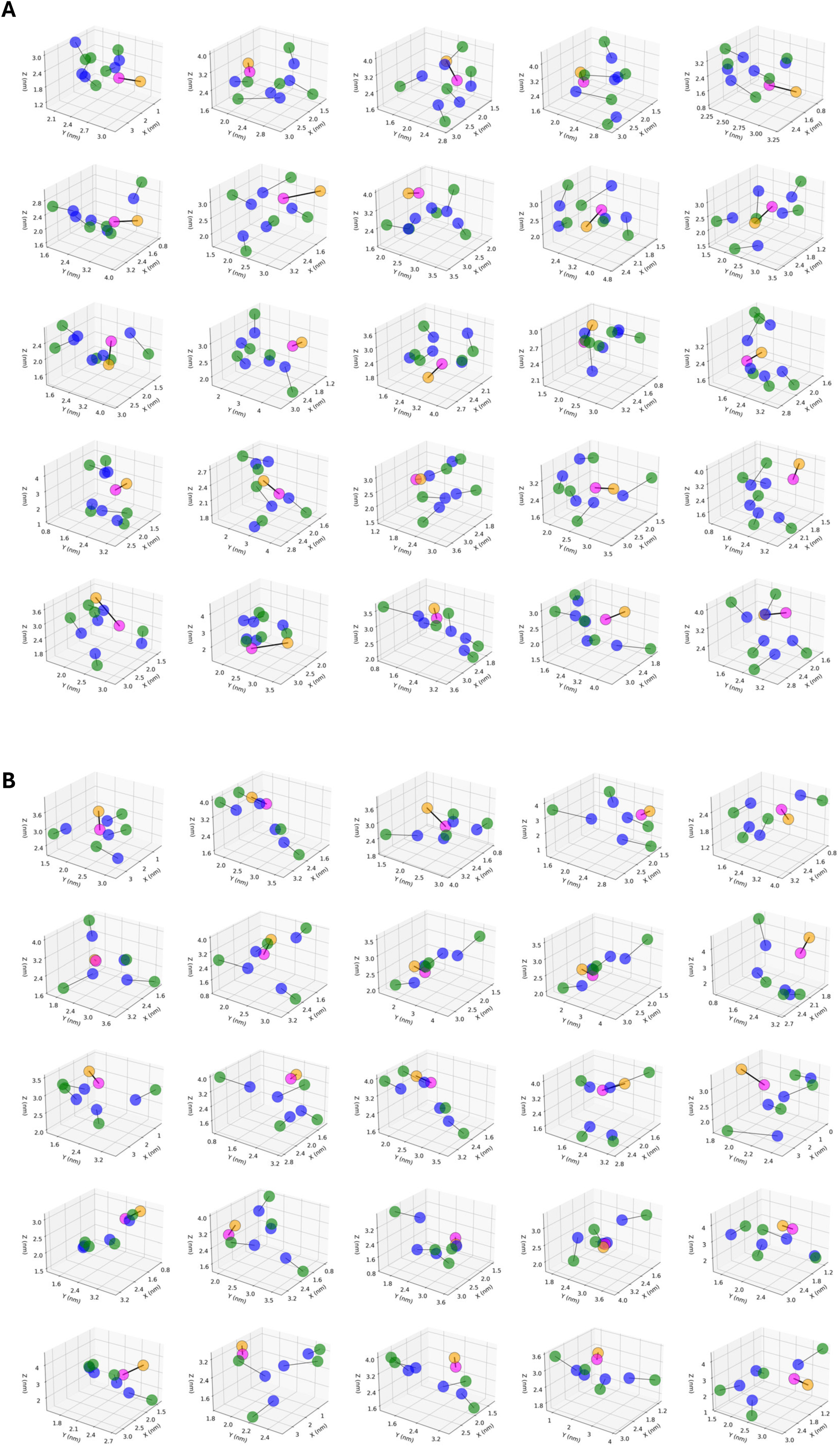
Peptide overlays above the IQR of the Fig. 3C.i RMSD distribution. Same analysis and representation as Supplementary Fig. 15, for peptides with RMSD above the upper quartile of Fig. 3C.i.

**Supplementary Figure 18.**
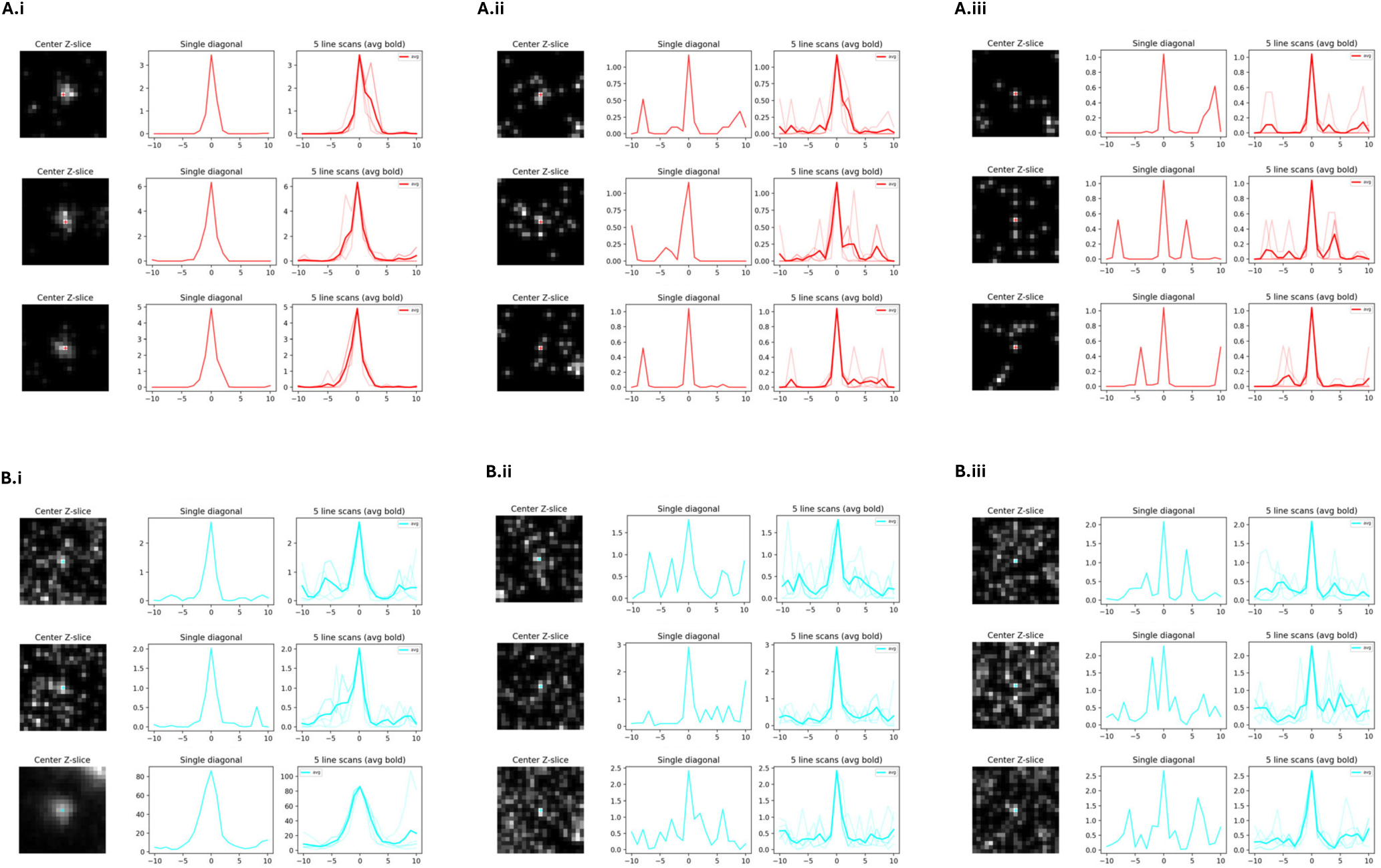
Emitter detection using Gaussian correlation to identify true fluorescent emitters. Confocal images were acquired as described in Methods (confocal z-stack acquisition). All images shown are raw, with no background subtraction or additional processing. Candidate fluorescent emitters were filtered by intensity and size and evaluated by Gaussian correlation (Methods, 3D emitter detection), using a correlation coefficient >0.70 as the acceptance threshold. Each subpanel (A.i–A.iii, B.i–B.iii) shows three representative emitters. For each emitter, a central z-slice image is displayed with the emitter center marked by a cross. From this center, one diagonal line scan (upper left to lower right) is shown, along with 3 additional line scans: upper right to lower left, vertical (top to bottom), and horizontal (left to right). Intensity profiles from these 4 line scans are plotted, and their average is shown as a bold trace. In the line-scan plots, the x-axis denotes distance along the line scan and the y-axis denotes fluorescence intensity (arbitrary units). (A) Atto647N emitters. (A.i) Accepted emitters with high Gaussian correlation (≈0.9), drawn from 27,426 Atto647N emitters across three gels and eight regions. (A.ii) Accepted emitters near the Gaussian-correlation threshold (≈0.7), drawn from the same 27,426 Atto647N emitters. (A.iii) Rejected emitters with Gaussian correlation <0.7, drawn from the 21,705 emitters that failed the Gaussian-correlation threshold after intensity and size filtering (49,131 total candidates across three gels and eight regions). (B) Fluorescein emitters. (B.i) Accepted emitters with high Gaussian correlation (≈0.9), drawn from 125,395 fluorescein emitters across three gels and eight regions. (B.ii) Accepted emitters near the Gaussian-correlation threshold (≈0.7), drawn from the same 125,395 fluorescein emitters. (B.iii) Rejected emitters with Gaussian correlation <0.7, drawn from the 462,408 emitters that failed the Gaussian-correlation threshold after intensity and size filtering (587,803 total candidates across three gels and eight regions).

**Supplementary Figure 19.**
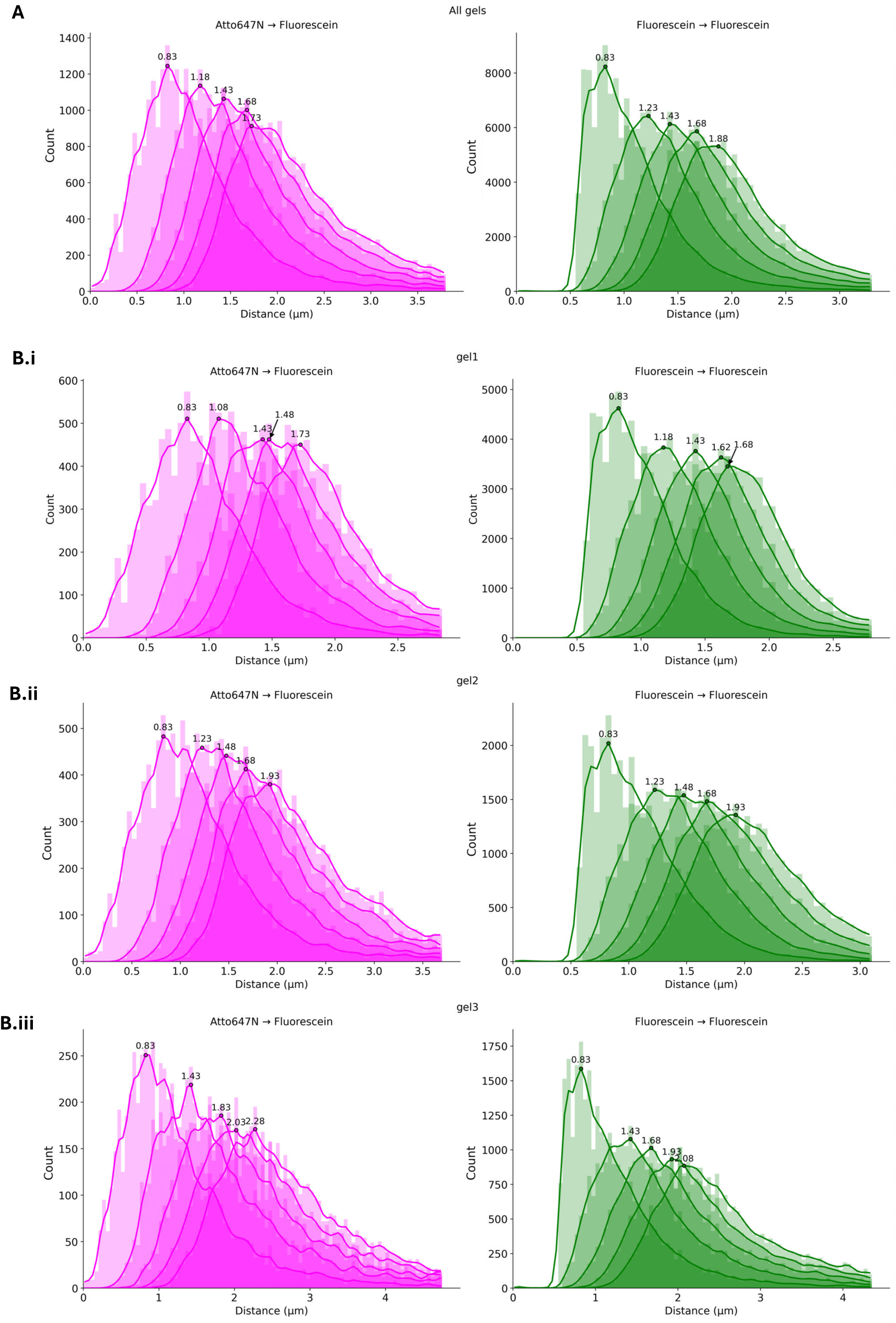
k-nearest-neighbor (kNN) 1–5 distance distributions for Atto647N–fluorescein (magenta) and fluorescein–fluorescein (green). Histograms use 0.05 µm bins with Gaussian smoothing (σ = 1 bin). Peaks are defined as the maxima of the smoothed distributions. n denotes the number of distances per kNN rank. Summary distance statistics corresponding to each panel are reported in **Tables S5–S8**. A) All gels combined (3 gels, 8 regions): Atto647N–fluorescein, n = 27,426; fluorescein–fluorescein, n = 125,395. (See pooled distance statistics in **Table S5**.) B) Partitioned by four-round gelation experiment (biological replicates); each ∼1000× gel was generated by an independent four-network casting sequence, initiated from an independently cast first-round ∼18× gel. B.i) Gel 1 (2 regions): Atto647N–fluorescein, n = 9,266; fluorescein–fluorescein, n = 63,733 (statistics in **Table S6**). B.ii) Gel 2 (4 regions): Atto647N–fluorescein, n = 11,671; fluorescein–fluorescein, n = 32,765 (statistics in Table S7). B.iii) Gel 3 (2 regions): Atto647N–fluorescein, n = 6,489; fluorescein–fluorescein, n = 28,897 (statistics in **Table S8**).

**Supplementary Figure 20.**
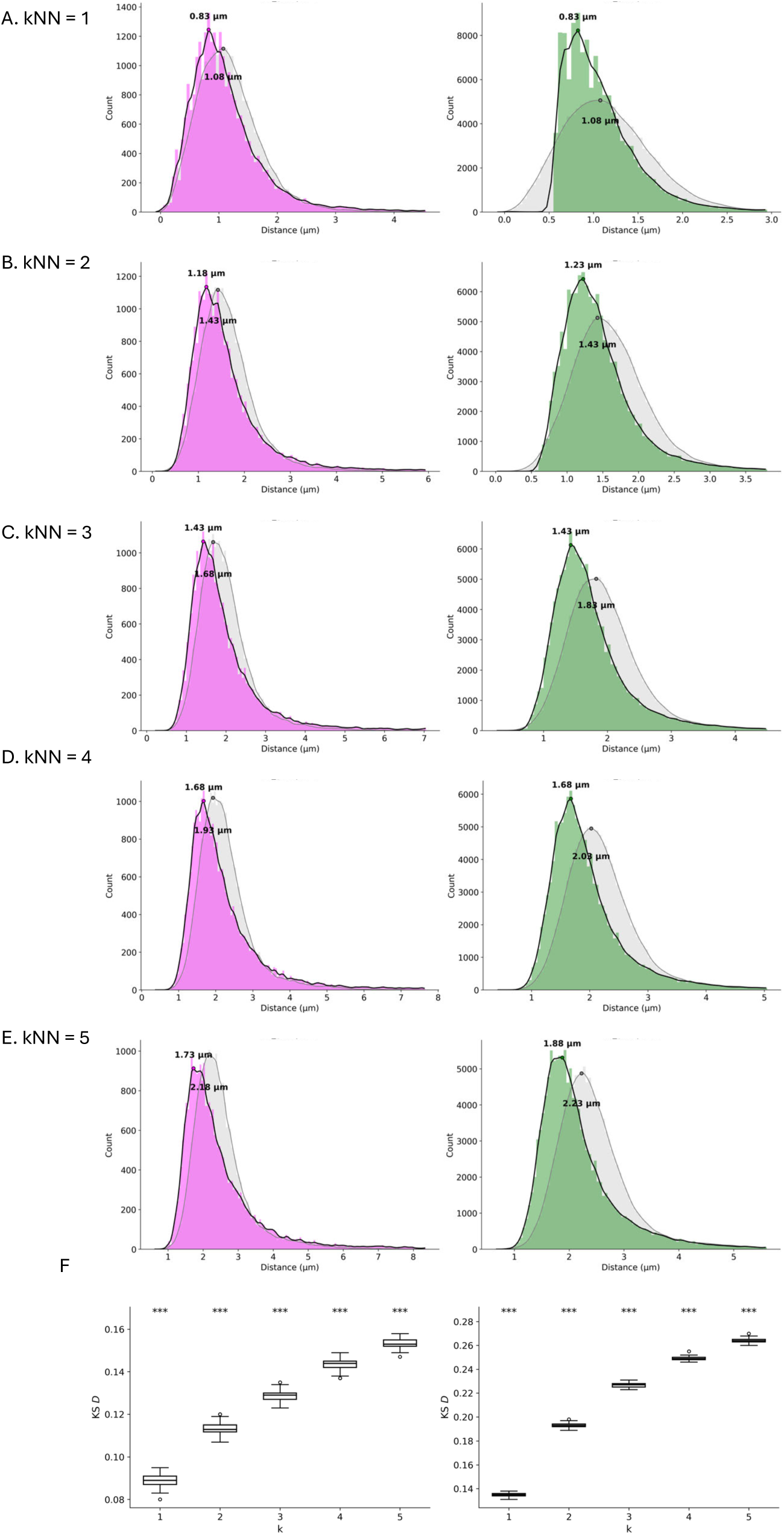
Randomization analysis of k-nearest-neighbor (kNN) distance distributions, all gels. Each region is a three-dimensional imaging volume containing Atto647N and fluorescein emitters, each represented by an (x, y, z) coordinate. For the randomized control, emitter counts were preserved and all coordinates were randomly chosen within the same volume. kNN distance distributions were recomputed from the randomized coordinates. Histograms use 0.05 µm bins with Gaussian smoothing (σ = 1 bin). Peak positions are defined as the bin center corresponding to the global maximum of the smoothed distribution. Distances are aggregated across eight regions from three gels. Observed Atto647N–fluorescein distributions are shown in magenta with randomized controls in gray (n = 27,426 distances per kNN rank). Observed fluorescein–fluorescein distributions are shown in green with randomized controls in gray (n = 125,395 distances per kNN rank). Panels A–E show kNN ranks 1–5. Panel F shows robustness across 100 randomizations assessed using two-sample Kolmogorov–Smirnov tests (***, p < 0.001); KS D statistics report effect size. Corresponding Kolmogorov–Smirnov D statistics by kNN rank (1–5) are reported in **Table S9**. Supplementary Figures 21–23 show the same randomization test as Supplementary Figure 20, with distances aggregated at the level of individual gels (statistics in **Tables S10-12**).

**Supplementary Figure 21.**
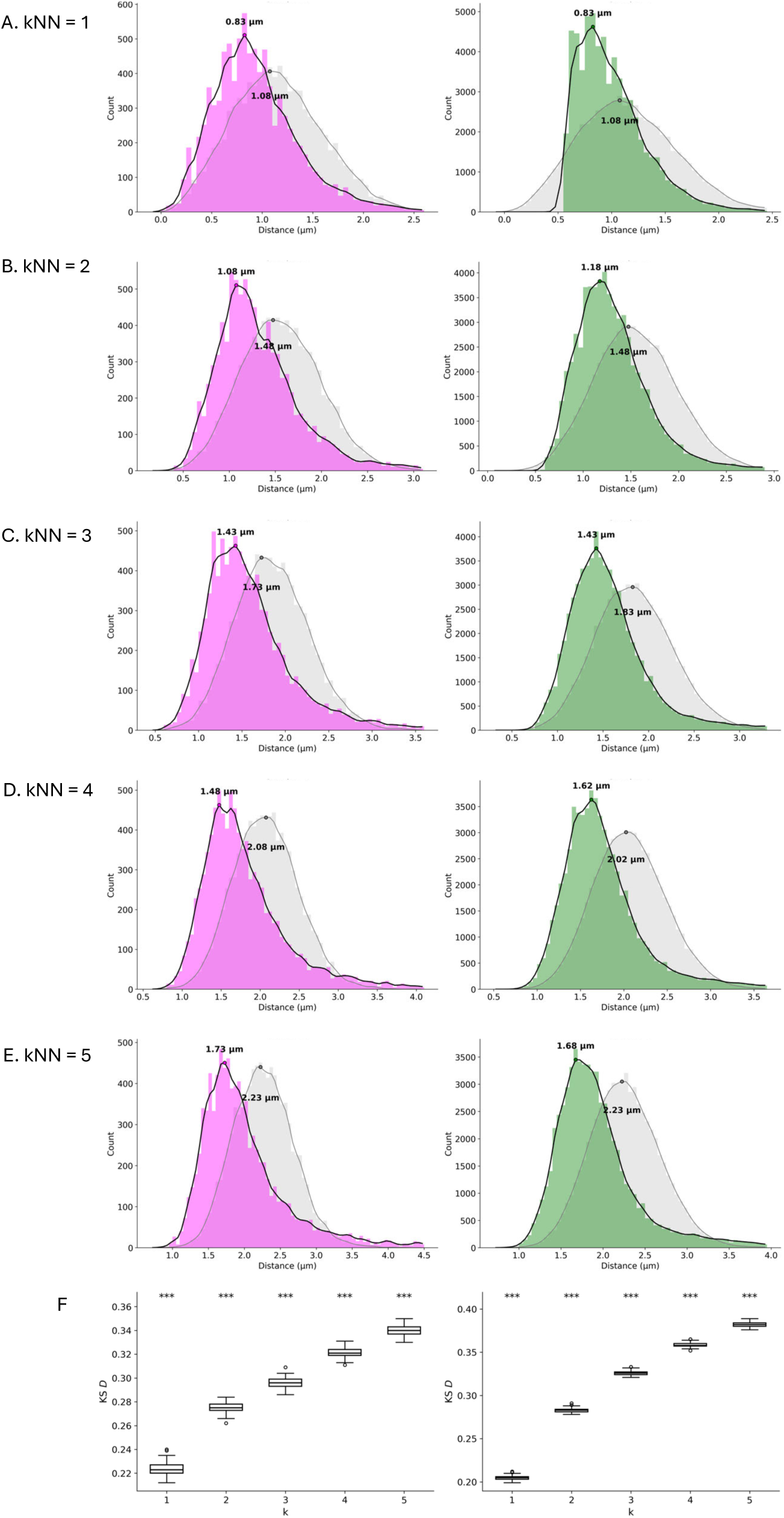
Randomization analysis of k-nearest-neighbor (kNN) distance distributions for gel 1 (statistics in Table S10).

**Supplementary Figure 22.**
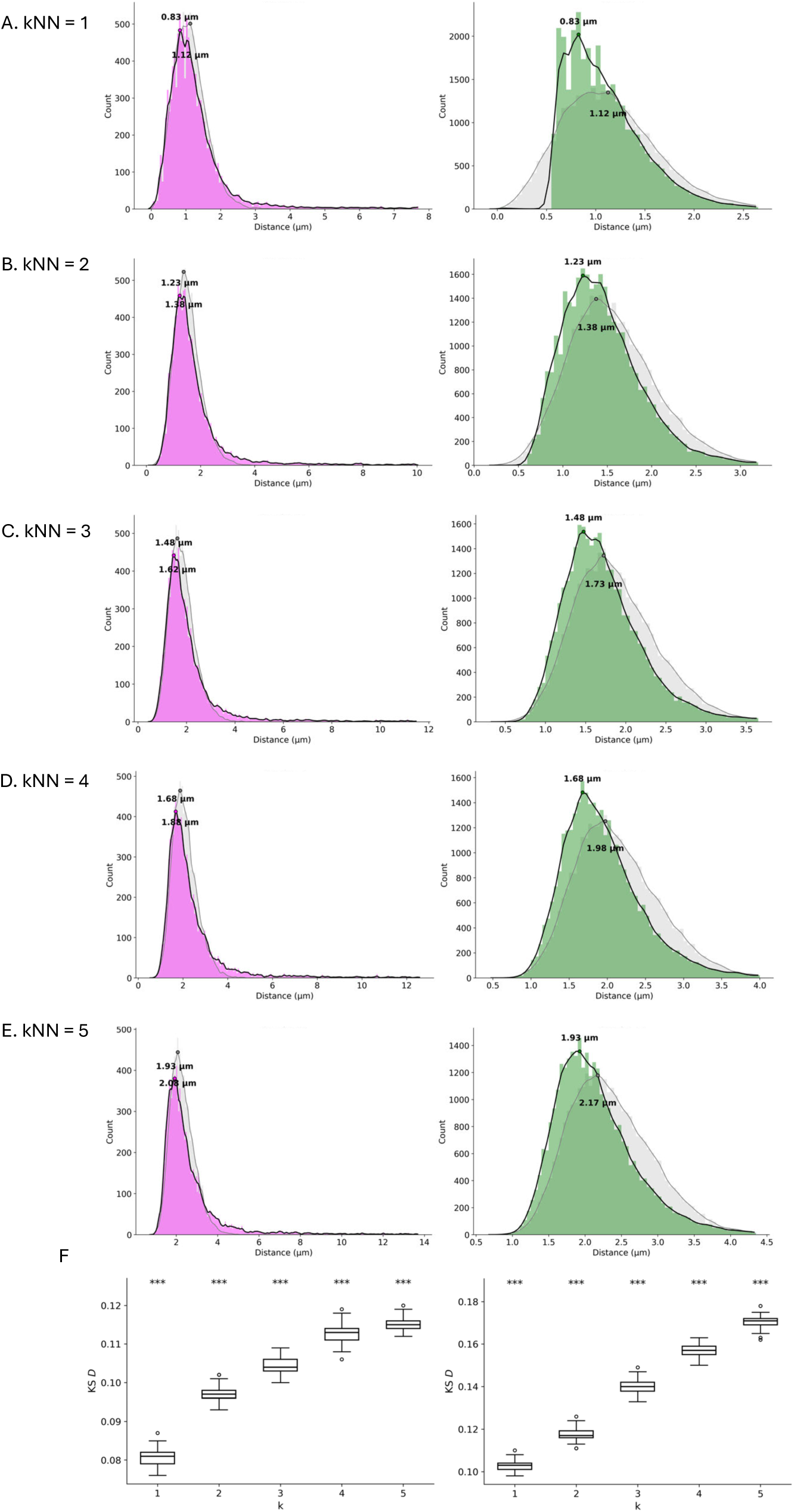
Randomization analysis of k-nearest-neighbor (kNN) distance distributions for gel 2 (statistics in Table S11).

**Supplementary Figure 23.**
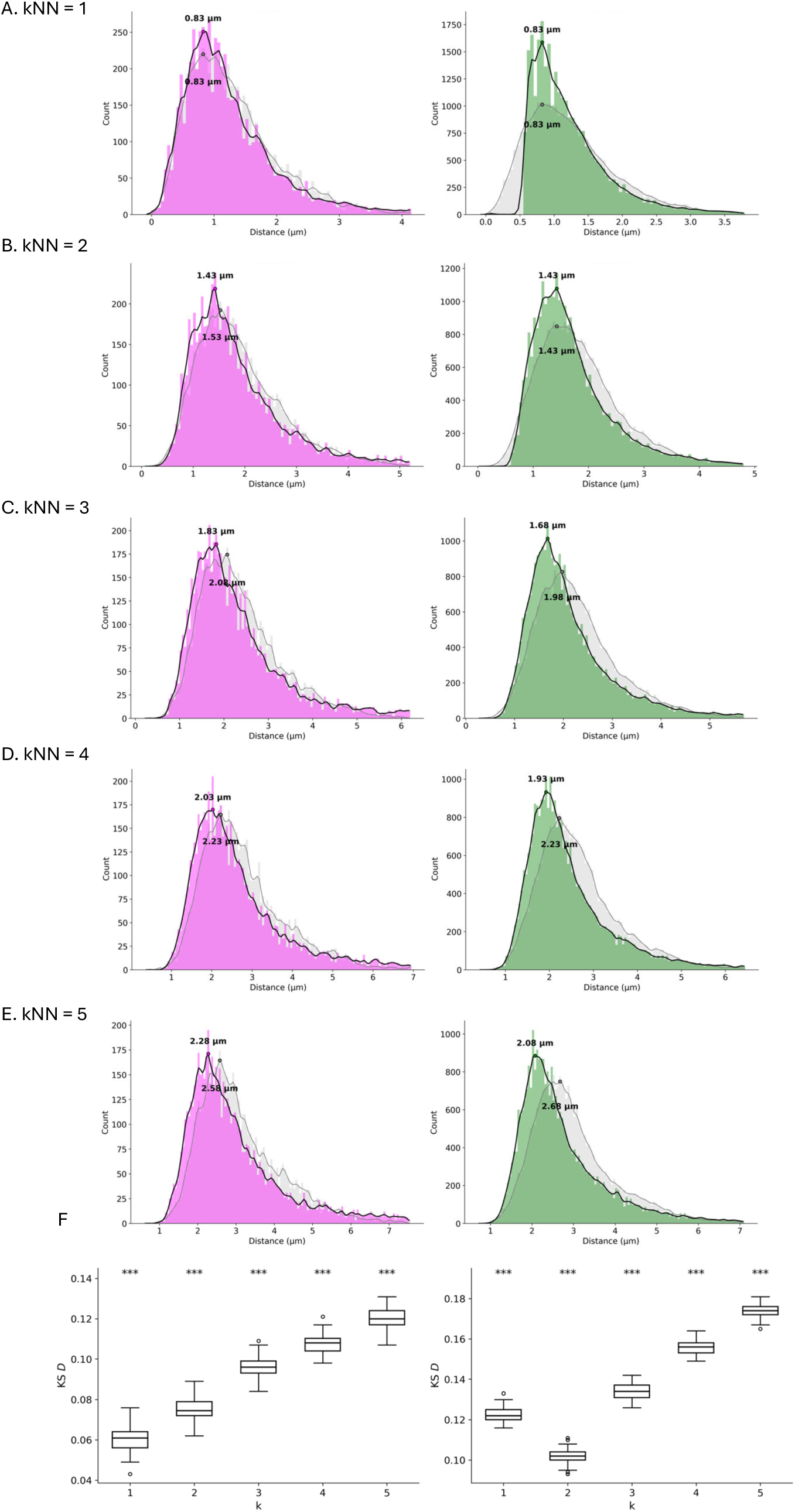
Randomization analysis of k-nearest-neighbor (kNN) distance distributions for gel 3 (statistics in Table S12).

**Supplementary Figure 24.**
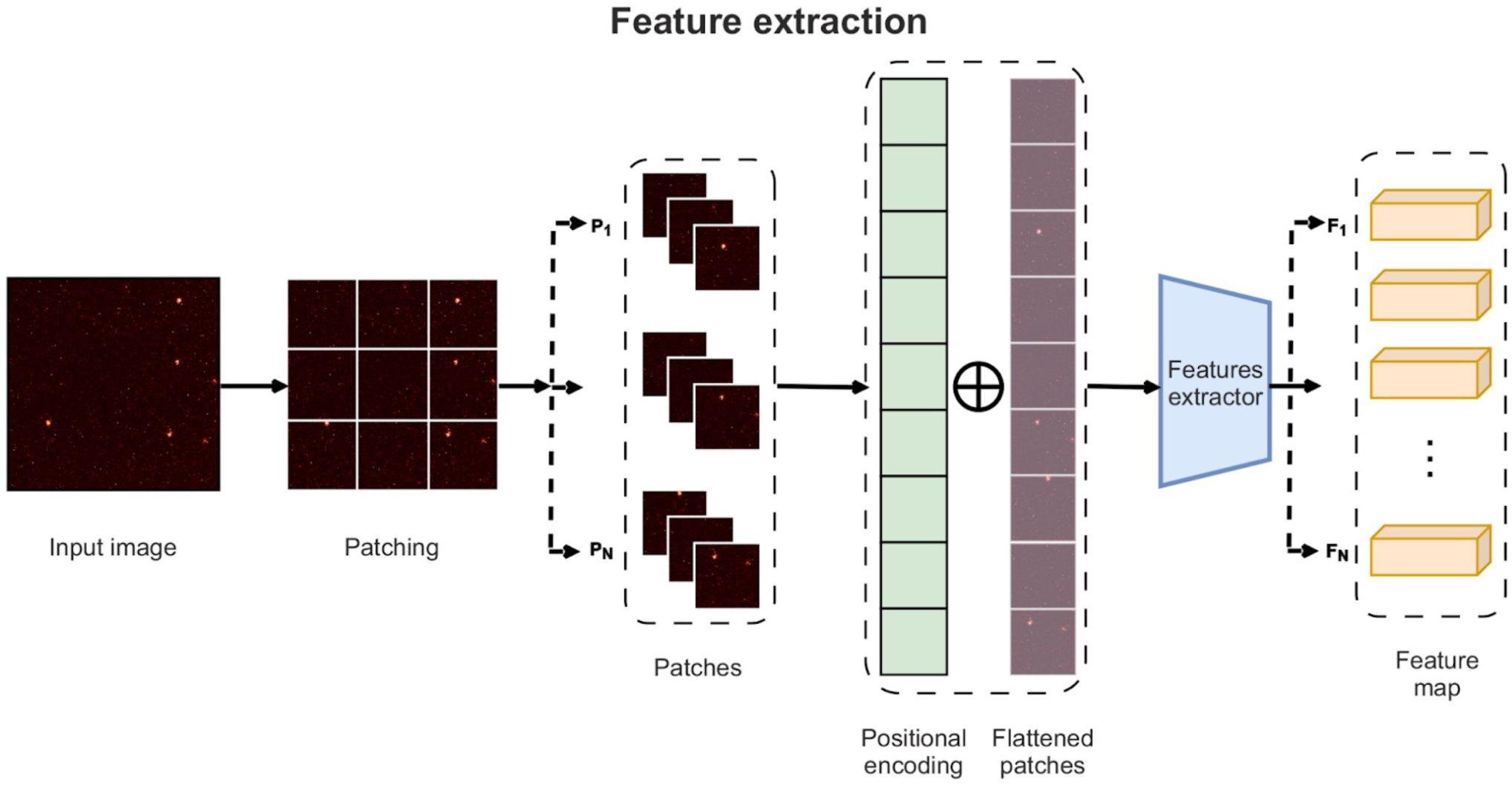
Preprocessing and transformer-based feature extraction pipeline for microscopy images. Raw fluorescence microscopy images are first partitioned into fixed-size patches. Each patch is flattened into a one-dimensional vector and supplemented with positional encodings to retain spatial information. The resulting sequence of embedded patches is processed by a transformer encoder, which integrates local texture features with global spatial relationships. The output is a structured feature map that serves as input for downstream graph-based modeling.

**Supplementary Figure 25.**
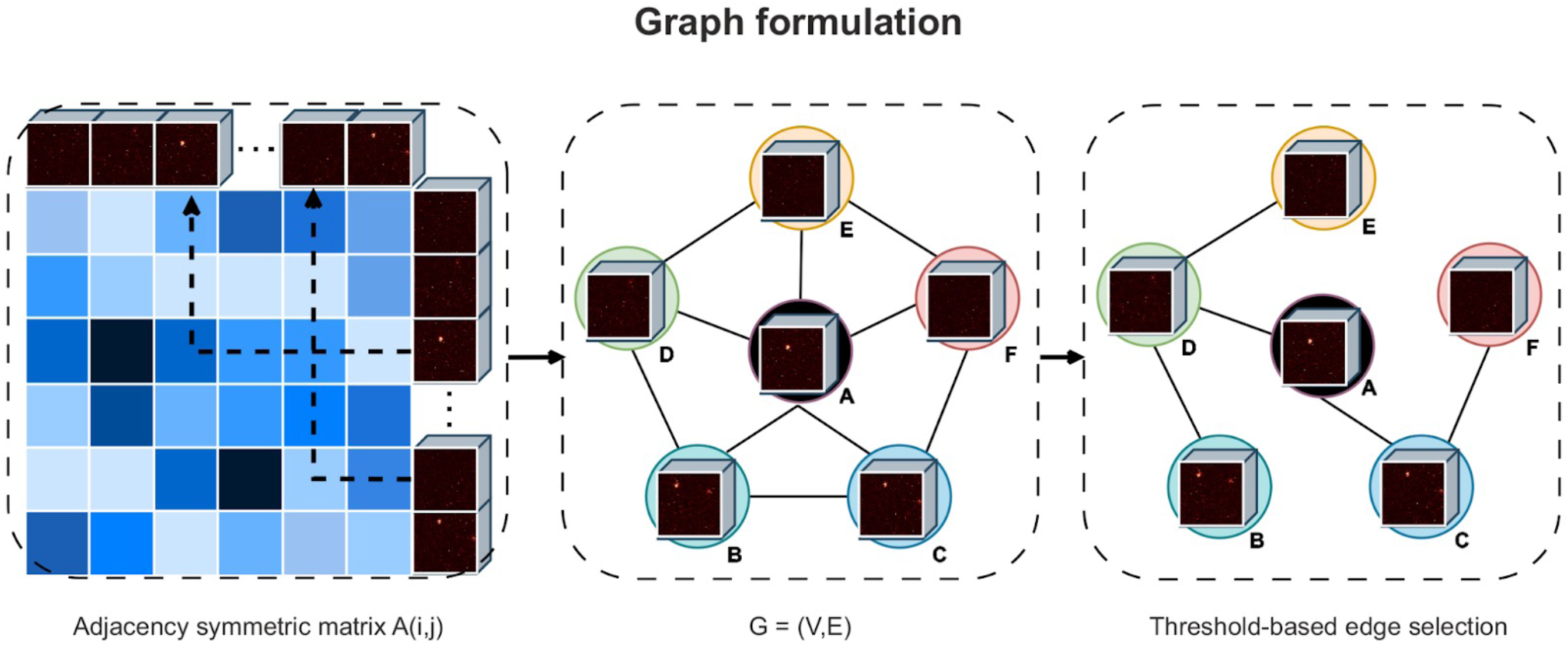
Graph formulation from transformer-derived patch features. Each node corresponds to a transformer-derived patch feature. Pairwise similarities between feature vectors define edge weights, forming a weighted, undirected graph G(V, E). Edges are retained only when their similarity exceeds a predefined threshold. The resulting symmetric adjacency matrix A(i, j) encodes structural relationships across the image and enables graph-based segmentation and clustering.

**Supplementary Figure 26.**
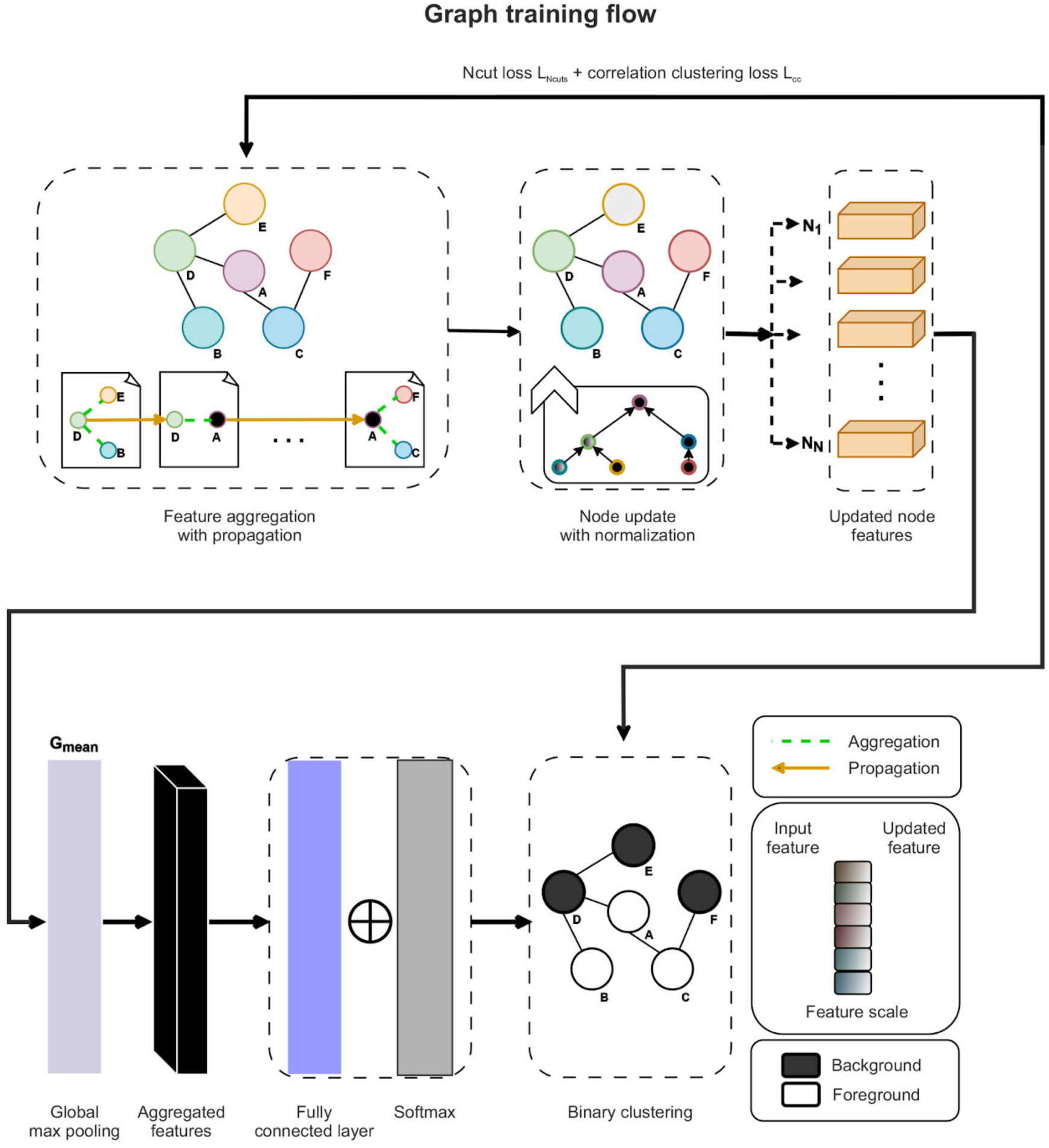
Unsupervised graph-based segmentation framework integrating Normalized Cut and correlation clustering losses. Node features are iteratively updated through aggregation, propagation, and normalization steps to refine graph embeddings. The training objective combines the Normalized Cut (NCut) loss and a correlation clustering loss to promote coherent partitioning. A global max pooling layer aggregates node-wise information, followed by a fully connected layer and softmax classifier that assign each node to foreground or background. The combined loss enforces consistent separation of structurally similar regions within the graph.

**Supplementary Figure 27.**
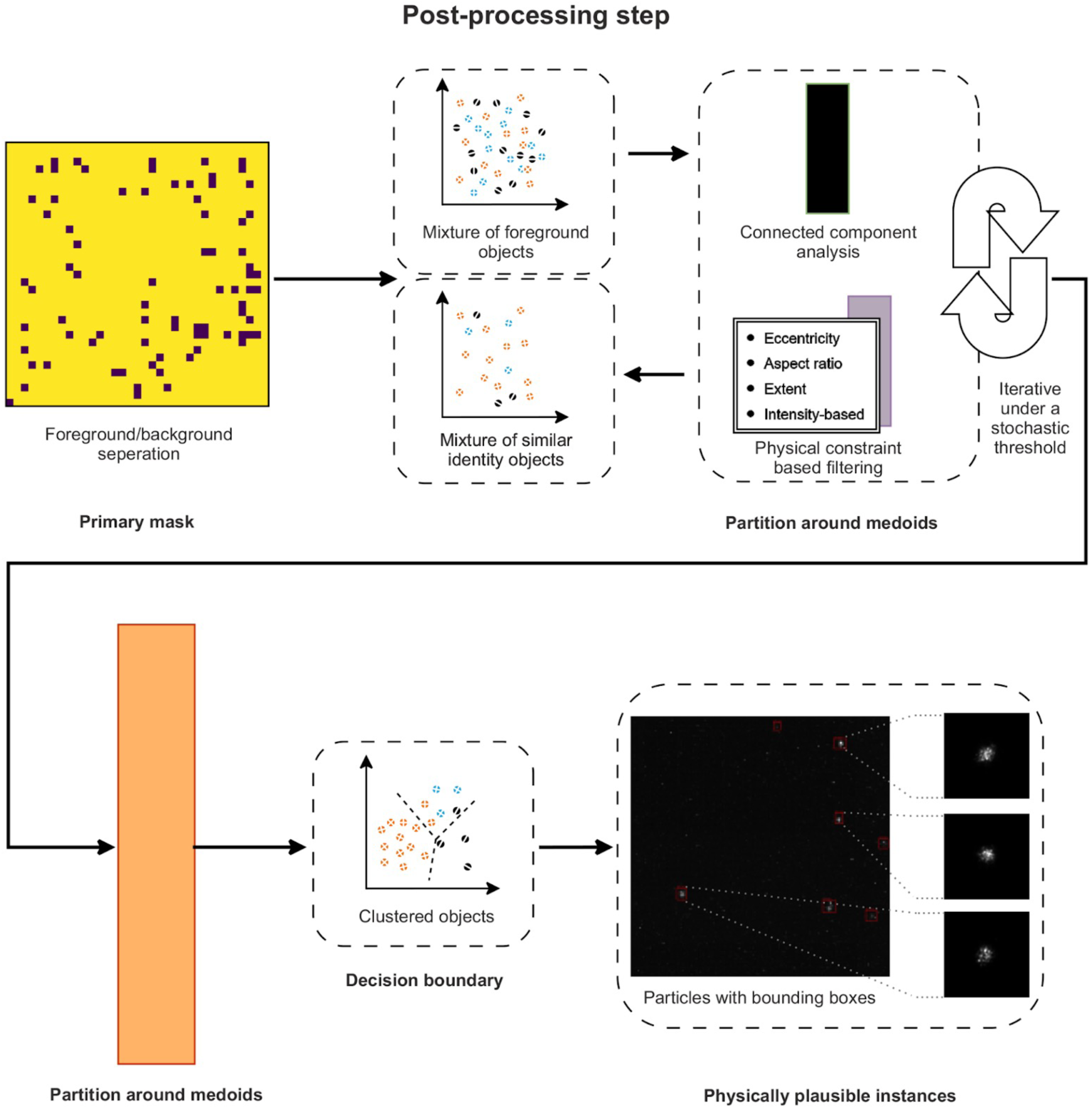
Post-segmentation analysis for separation of heterogeneous foreground objects. Following foreground–background separation, connected components are extracted from the primary mask. Objects are filtered using geometric and intensity-based descriptors, including eccentricity, aspect ratio, extent, and intensity criteria. Partitioning around medoids clustering is subsequently applied to group objects of similar identity. Decision boundaries delineate the resulting clusters, and bounding boxes indicate individual, physically plausible particle instances.

**Supplementary Figure 28.**
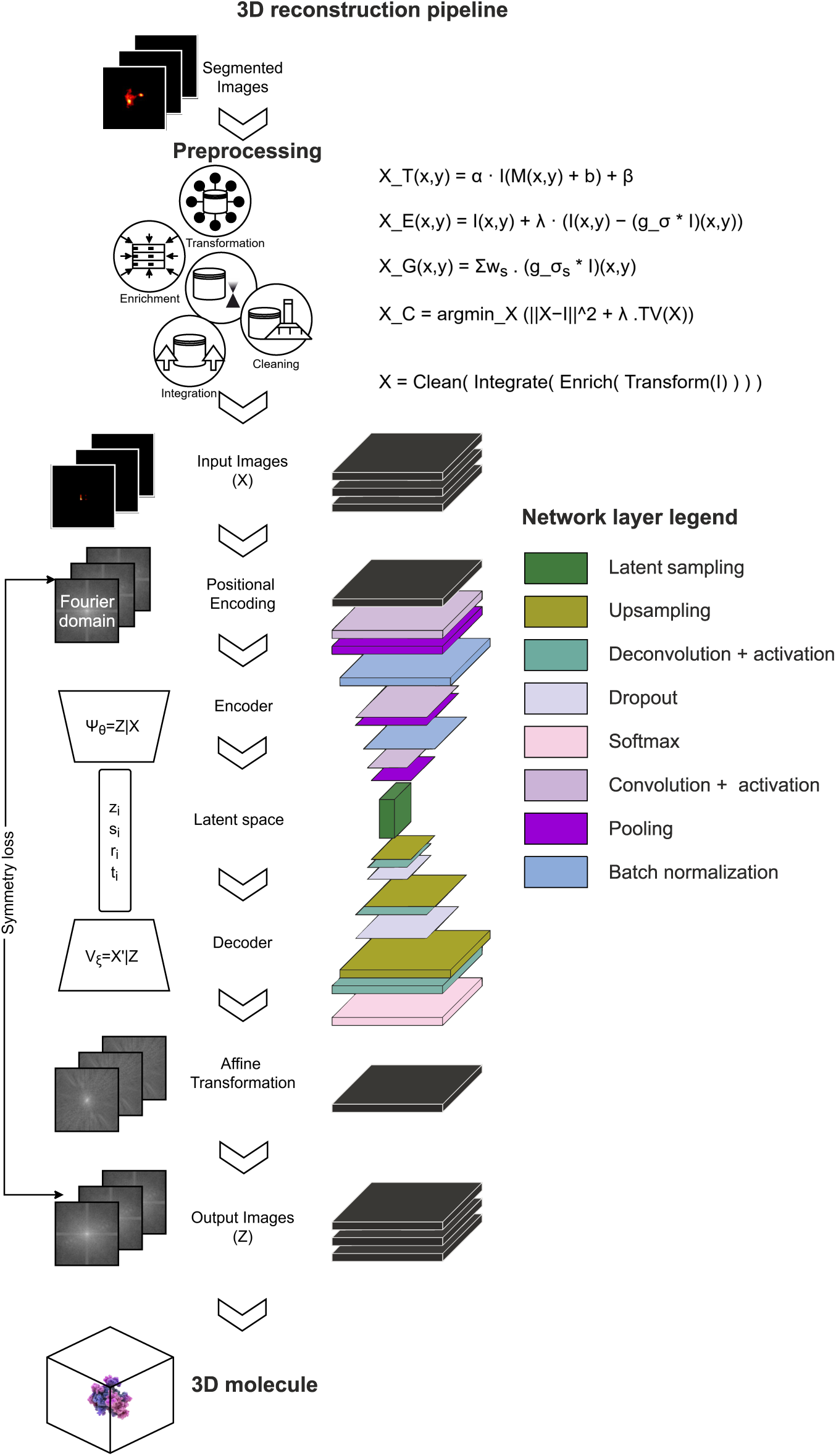
End-to-end *ab initio* 3D reconstruction pipeline from preprocessed 2D images. The workflow begins with segmented images and preprocessing steps including intensity normalization, background suppression, Gaussian smoothing, and total variation–based enrichment. Preprocessed inputs are mapped into a Fourier domain representation and encoded using a convolutional variational autoencoder (VAE). The encoder performs amortized inference to learn latent variables under a combined objective incorporating KL divergence and symmetry loss, followed by affine transformations in the decoder to generate consistent reconstructions. Reconstructed 2D slices are stacked and integrated in the spatial domain to produce a volumetric density map V(u, v, w), yielding an ab initio 3D molecular model (**Fig. 4**).

**Supplementary Figure 29.**
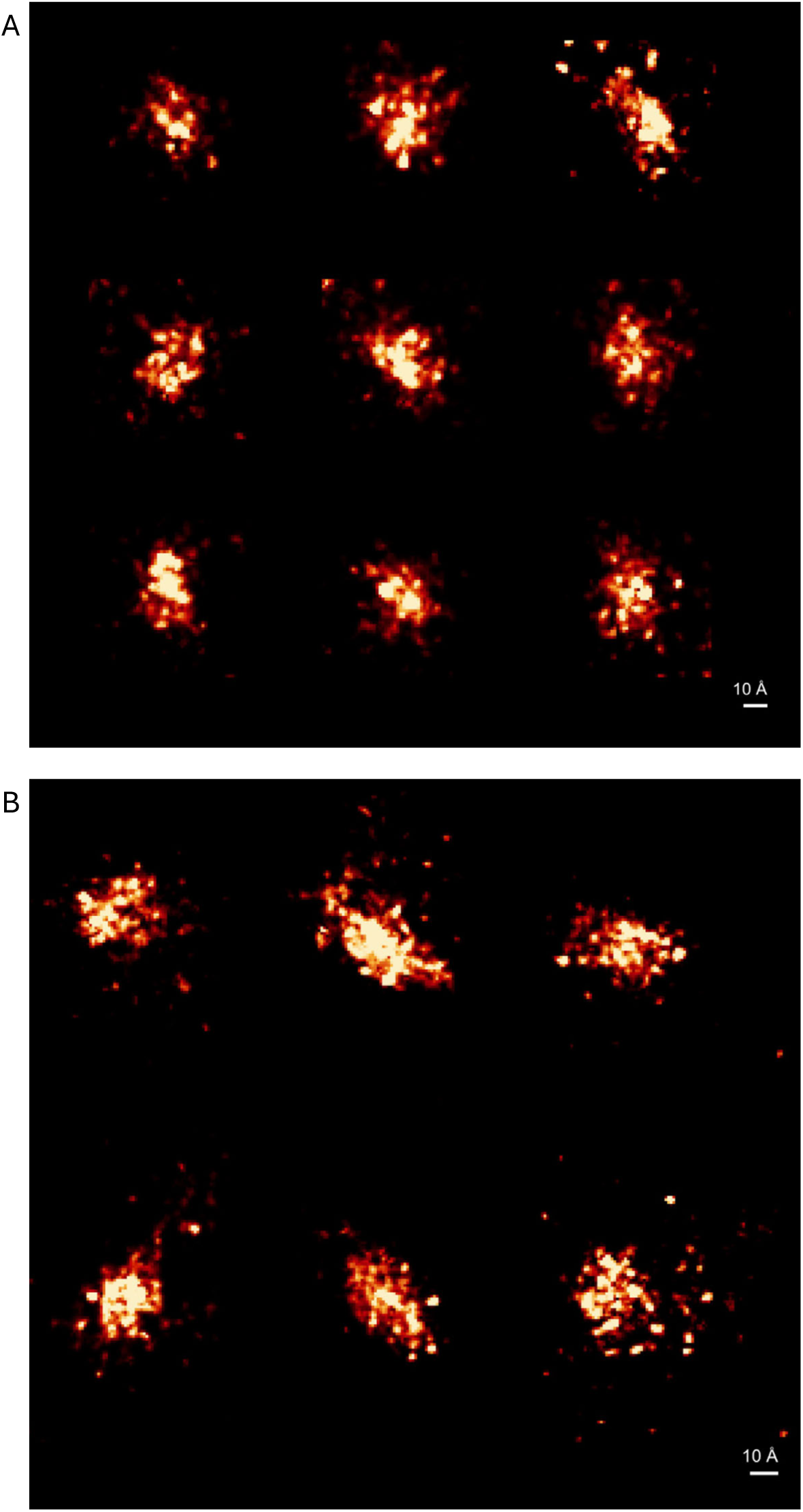
Additional GFP examples at different expansion factors. Representative 2D ONE GFP images are shown at (A) 50-fold expansion and (B) 100-fold expansion.

**Supplementary Figure 30.**
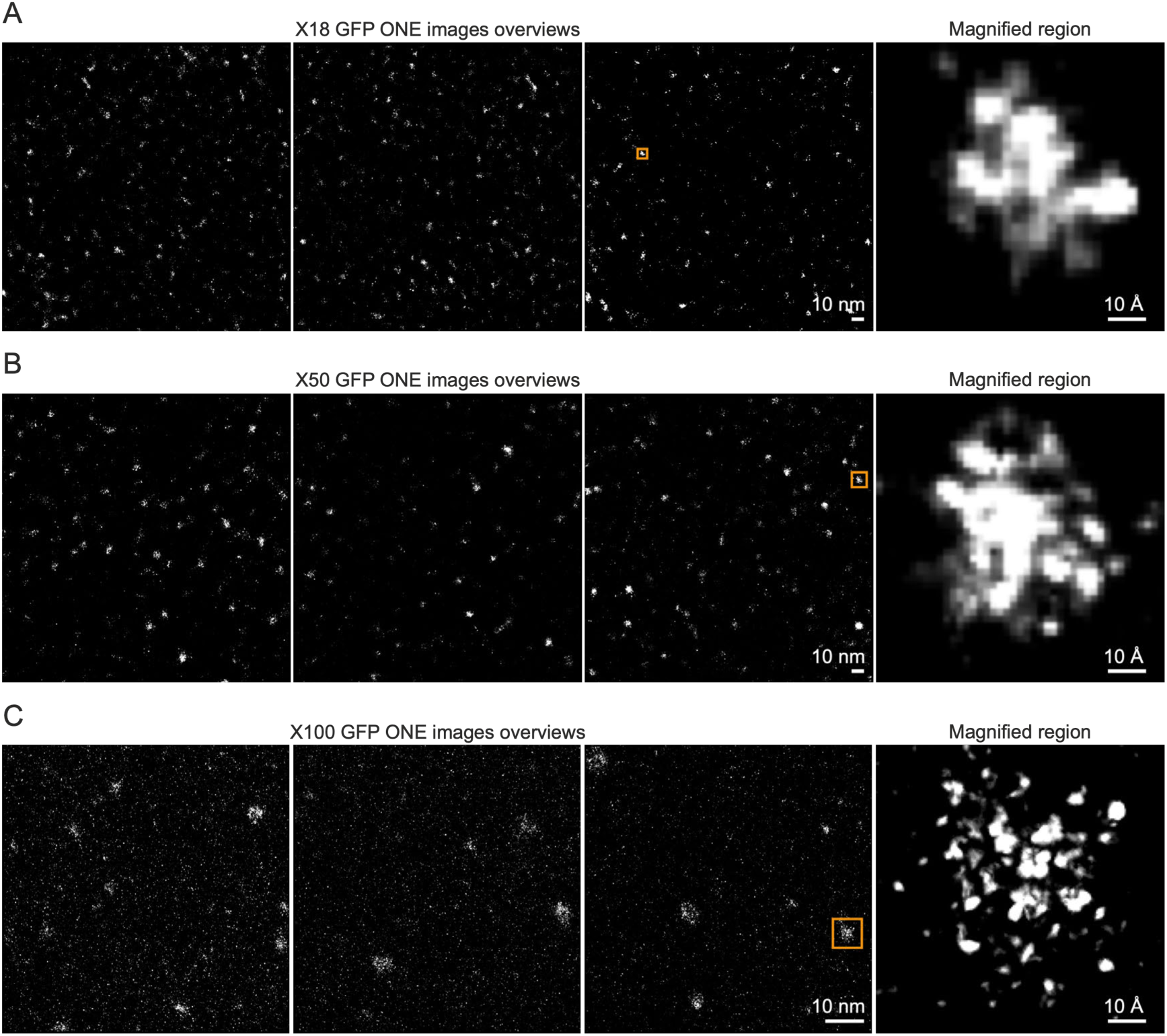
GFP imaged with ONE microscopy across expansion factors. Representative GFP images at (A) 18-fold, (B) 50-fold, and (C) 100-fold linear expansion. Each panel shows a field-of-view overview and a corresponding magnified region from the same image.

**Supplementary Figure 31.**
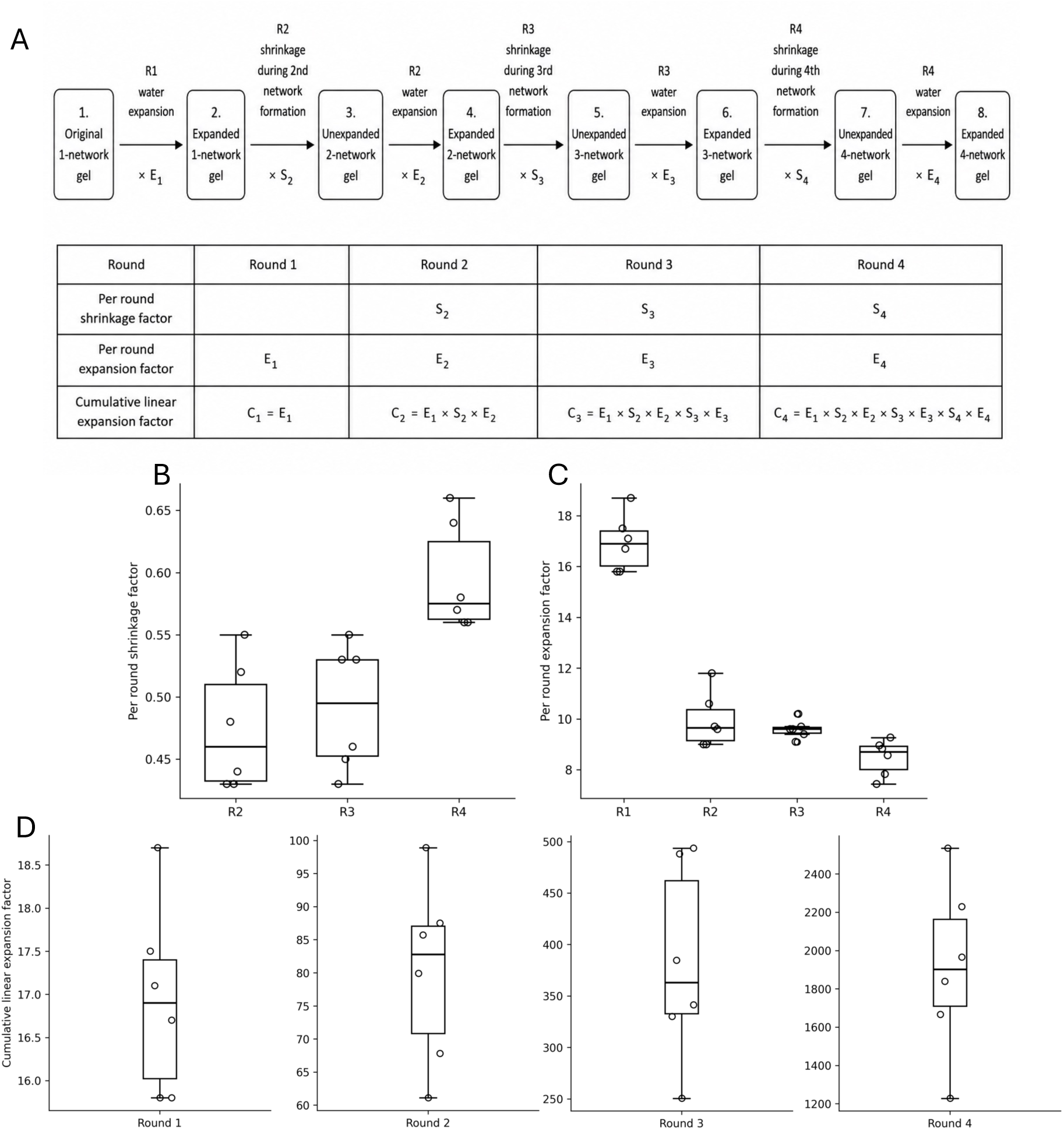
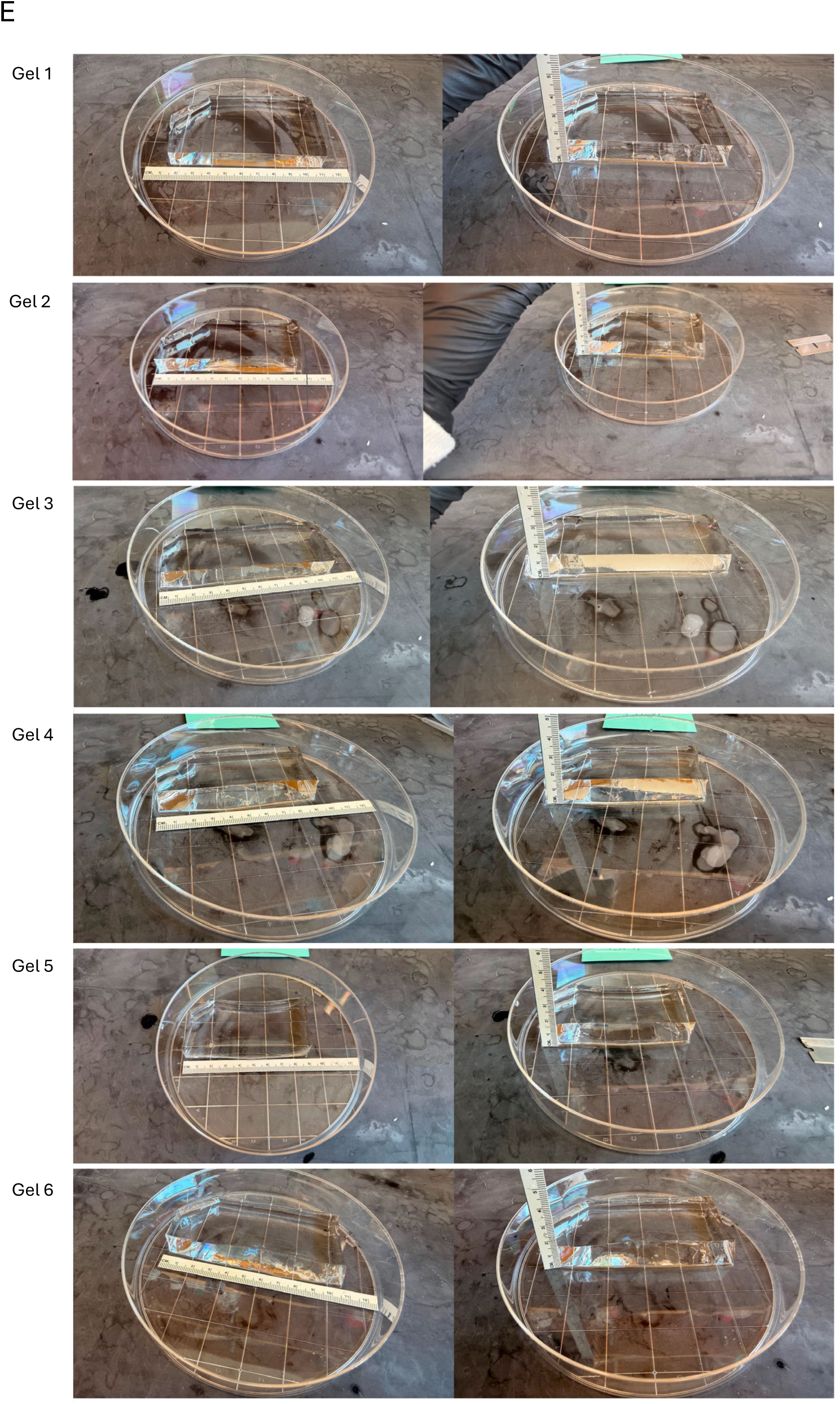
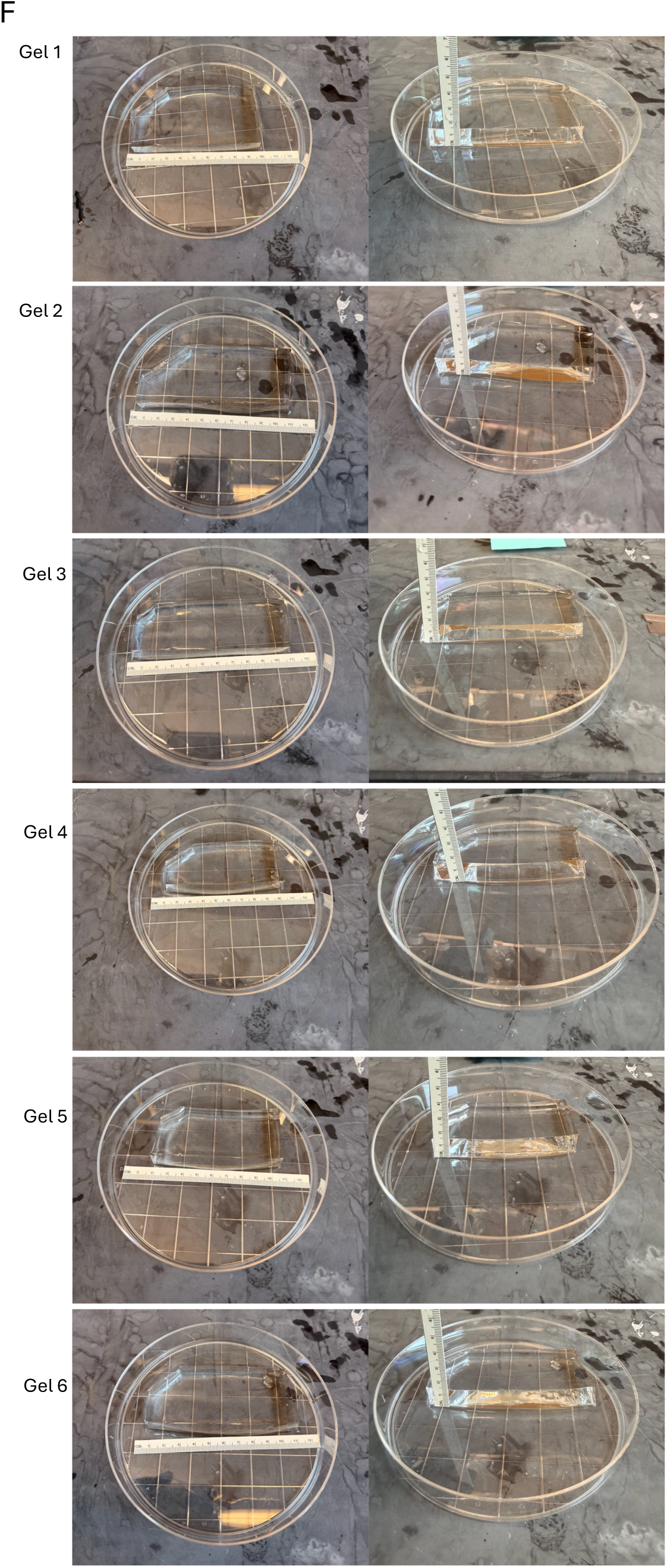
Four-network gels generated using a modified 1000ExM process. In a modified 1000ExM process, each sequential casting round includes two incubations in activated monomer solution before polymerization. The second incubation replaces the monomer-depleted and/or diluted first solution with fresh activated monomer solution, for further infiltration. (A) Schematic showing how per-round shrinkage factors, per-round expansion factors, and cumulative linear expansion factors were calculated during four-network gel formation. After the first gel was expanded, each subsequent casting round included incubation in activated monomer solution (two rounds), polymerization of the next network, measurement of the fully gelled composite, and re-expansion in water. Per-round shrinkage factors were measured after full gelation and calculated as gel length after polymerization divided by gel length before monomer solution addition and polymerization (i.e., at the end of the last expansion round). Per-round expansion factors were calculated as gel length after water expansion divided by gel length before water expansion. Cumulative linear expansion factors were calculated by multiplying the sequential shrinkage and expansion factors through all rounds up to the current round. (B) Per-round shrinkage factors for rounds 2–4. Boxplots (throughout this figure) show the median (middle line), interquartile range (top and bottom of box), whiskers extending to the farthest data point within 1.5×IQR of each box edge, and individual gel measurements as open circles. (C) Per-round expansion factors for rounds 1–4. Note that some circles have similar or identical values - this is not an error, and we include all the raw data in Table S18. (D) Cumulative linear expansion factors after each expansion round, with round 4 gels reaching up to 2534× linear expansion. (E) Images of fully expanded round 4 gels. (F) Images of round 4 gels (made at the same time) maintained at room temperature on a dry surface for 98 hours (chosen arbitrarily). Gel linear size and height remained roughly unchanged over this period, with linear measurements showing a 0.3±0.5% change in expansion factor.

## Supplementary Tables

**Table S1:** x, y, and z dimensions and total volume of the eight imaged 3D regions.

| gel_region | Z thickness (μm) | Y width (μm) | X height (μm) | Volume (μm<sup>3</sup>) |
| --- | --- | --- | --- | --- |
| gel1:region1 | 4.4 | 279.09 | 279.09 | 3.42×10<sup>5</sup> |
| gel1:region2 | 4.4 | 279.09 | 279.09 | 3.42×10<sup>5</sup> |
| gel2:region1 | 3.4 | 139.59 | 232.59 | 1.10×10<sup>5</sup> |
| gel2:region2 | 3.4 | 226.49 | 131.4 | 1.01×10<sup>5</sup> |
| gel2:region3 | 3.4 | 223.31 | 134.95 | 1.02×10<sup>5</sup> |
| gel2:region4 | 3.4 | 179.18 | 90.82 | 5.53×10<sup>4</sup> |
| gel3:region1 | 2.2 | 1906.59 | 46.59 | 1.95×10<sup>5</sup> |
| gel3:region2 | 2.2 | 260.98 | 340.42 | 1.95×10<sup>5</sup> |

**Table S2:** Atto647N and fluorescein emitter counts after local maxima detection.

| gel_region | Atto647N emitters | Fluorescein emitters | Fluorescein / Atto647N |
| --- | --- | --- | --- |
| gel1:region1 | 584,761 | 436,827 | 0.75 |
| gel1:region2 | 589,176 | 437,280 | 0.74 |
| gel2:region1 | 172,503 | 145,811 | 0.85 |
| gel2:region2 | 1,309,470 | 709,989 | 0.54 |
| gel2:region3 | 472,152 | 152,744 | 0.32 |
| gel2:region4 | 98,142 | 82,268 | 0.84 |
| gel3:region1 | 349,057 | 287,299 | 0.82 |
| gel3:region2 | 17,664,588 | 16,559,923 | 0.94 |

**Table S3:**
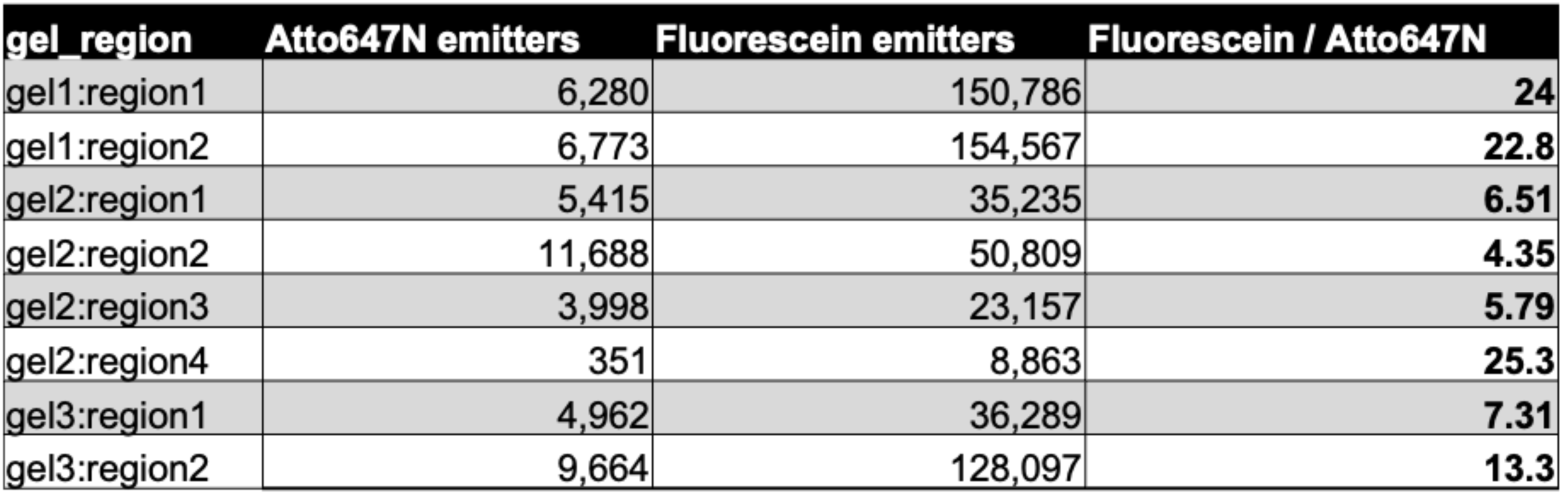
Atto647N and fluorescein emitter counts after local maxima detection and intensity- and size-based filtering

| gel_region | Atto647N emitters | Fluorescein emitters | Fluorescein / Atto647N |
| --- | --- | --- | --- |
| gel1:region1 | 6,280 | 150,786 | 24 |
| gel1:region2 | 6,773 | 154,567 | 22.8 |
| gel2:region1 | 5,415 | 35,235 | 6.51 |
| gel2:region2 | 11,688 | 50,809 | 4.35 |
| gel2:region3 | 3,998 | 23,157 | 5.79 |
| gel2:region4 | 351 | 8,863 | 25.3 |
| gel3:region1 | 4,962 | 36,289 | 7.31 |
| gel3:region2 | 9,664 | 128,097 | 13.3 |

**Table S4:** Atto647N and fluorescein emitter counts after local maxima detection, intensity- and size-based filtering, and Gaussian shape filtering

| gel_region | Atto647N emitters | Fluorescein emitters | Fluorescein / Atto647N |
| --- | --- | --- | --- |
| gel1:region1 | 4,445 | 31,567 | 7.1 |
| gel1:region2 | 4,821 | 32,166 | 6.67 |
| gel2:region1 | 3,254 | 7,206 | 2.21 |
| gel2:region2 | 6,074 | 13,444 | 2.21 |
| gel2:region3 | 2,133 | 8,573 | 4.02 |
| gel2:region4 | 210 | 3,542 | 16.9 |
| gel3:region1 | 2,225 | 8,161 | 3.67 |
| gel3:region2 | 4,264 | 20,736 | 4.86 |

**Table S5.**
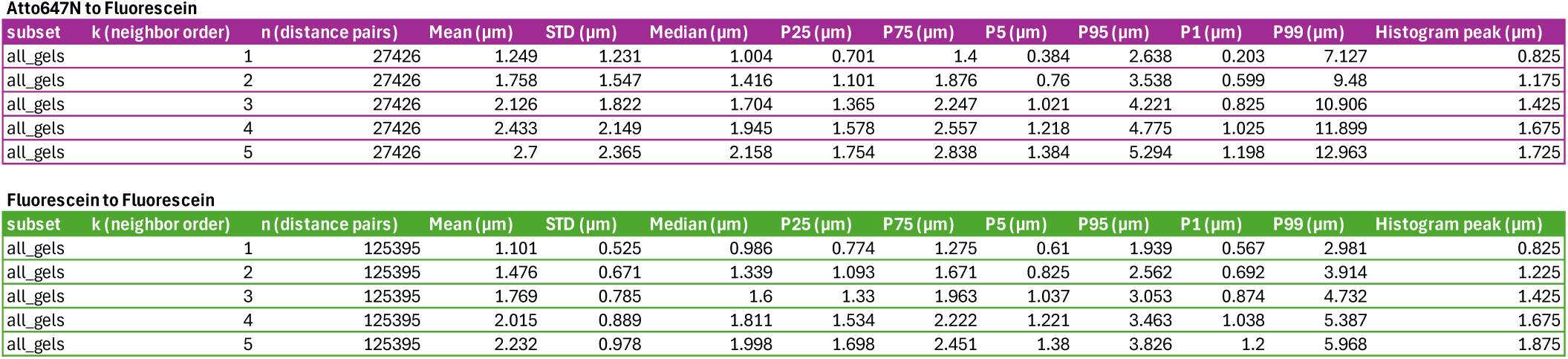
Pooled k-nearest neighbor (kNN) distance statistics (k = 1–5) across all gels and all regions (see **Supplementary Figure 19A**).

Atto647N to Fluorescein
| subset | k (neighbor order) | n (distance pairs) | Mean (μm) | STD (μm) | Median (μm) | P25 (μm) | P75 (μm) | P5 (μm) | P95 (μm) | P1 (μm) | P99 (μm) | Histogram peak (μm) |
| --- | --- | --- | --- | --- | --- | --- | --- | --- | --- | --- | --- | --- |
| all_gels | 1 | 27426 | 1.249 | 1.231 | 1.004 | 0.701 | 1.4 | 0.384 | 2.638 | 0.203 | 7.127 | 0.825 |
| all_gels | 2 | 27426 | 1.758 | 1.547 | 1.416 | 1.101 | 1.876 | 0.76 | 3.538 | 0.599 | 9.48 | 1.175 |
| all_gels | 3 | 27426 | 2.126 | 1.822 | 1.704 | 1.365 | 2.247 | 1.021 | 4.221 | 0.825 | 10.906 | 1.425 |
| all_gels | 4 | 27426 | 2.433 | 2.149 | 1.945 | 1.578 | 2.557 | 1.218 | 4.775 | 1.025 | 11.899 | 1.675 |
| all_gels | 5 | 27426 | 2.7 | 2.365 | 2.158 | 1.754 | 2.838 | 1.384 | 5.294 | 1.198 | 12.963 | 1.725 |

| subset | k (neighbor order) | n (distance pairs) | Mean (μm) | STD (μm) | Median (μm) | P25 (μm) | P75 (μm) | P5 (μm) | P95 (μm) | P1 (μm) | P99 (μm) | Histogram peak (μm) |
| --- | --- | --- | --- | --- | --- | --- | --- | --- | --- | --- | --- | --- |
| all_gels | 1 | 125395 | 1.101 | 0.525 | 0.986 | 0.774 | 1.275 | 0.61 | 1.939 | 0.567 | 2.981 | 0.825 |
| all_gels | 2 | 125395 | 1.476 | 0.671 | 1.339 | 1.093 | 1.671 | 0.825 | 2.562 | 0.692 | 3.914 | 1.225 |
| all_gels | 3 | 125395 | 1.769 | 0.785 | 1.6 | 1.33 | 1.963 | 1.037 | 3.053 | 0.874 | 4.732 | 1.425 |
| all_gels | 4 | 125395 | 2.015 | 0.889 | 1.811 | 1.534 | 2.222 | 1.221 | 3.463 | 1.038 | 5.387 | 1.675 |
| all_gels | 5 | 125395 | 2.232 | 0.978 | 1.998 | 1.698 | 2.451 | 1.38 | 3.826 | 1.2 | 5.968 | 1.875 |

**Table S6.** Pooled k-nearest neighbor (kNN) distance statistics (k = 1–5) across all regions from gel 1 (see **Supplementary Figure 19B.i**).

Atto647N to Fluorescein
| subset | k (neighbor order) | n (distance pairs) | Mean (μm) | STD (μm) | Median (μm) | P25 (μm) | P75 (μm) | P5 (μm) | P95 (μm) | P1 (μm) | P99 (μm) | Histogram peak (μm) |
| --- | --- | --- | --- | --- | --- | --- | --- | --- | --- | --- | --- | --- |
| gel1 | 1 | 9266 | 0.986 | 0.704 | 0.883 | 0.643 | 1.186 | 0.35 | 1.841 | 0.203 | 2.834 | 0.825 |
| gel1 | 2 | 9266 | 1.381 | 0.868 | 1.24 | 1.017 | 1.559 | 0.713 | 2.311 | 0.583 | 3.569 | 1.075 |
| gel1 | 3 | 9266 | 1.665 | 1.004 | 1.5 | 1.249 | 1.832 | 0.96 | 2.72 | 0.8 | 4.21 | 1.425 |
| gel1 | 4 | 9266 | 1.898 | 1.136 | 1.691 | 1.437 | 2.06 | 1.153 | 3.124 | 0.989 | 4.777 | 1.475 |
| gel1 | 5 | 9266 | 2.101 | 1.26 | 1.872 | 1.602 | 2.256 | 1.305 | 3.45 | 1.151 | 5.531 | 1.725 |

| subset | k (neighbor order) | n (distance pairs) | Mean (μm) | STD (μm) | Median (μm) | P25 (μm) | P75 (μm) | P5 (μm) | P95 (μm) | P1 (μm) | P99 (μm) | Histogram peak (μm) |
| --- | --- | --- | --- | --- | --- | --- | --- | --- | --- | --- | --- | --- |
| gel1 | 1 | 63733 | 1.021 | 0.422 | 0.937 | 0.758 | 1.182 | 0.609 | 1.669 | 0.567 | 2.295 | 0.825 |
| gel1 | 2 | 63733 | 1.342 | 0.53 | 1.254 | 1.047 | 1.515 | 0.809 | 2.101 | 0.683 | 3.025 | 1.175 |
| gel1 | 3 | 63733 | 1.592 | 0.606 | 1.493 | 1.268 | 1.758 | 1.011 | 2.484 | 0.862 | 3.559 | 1.425 |
| gel1 | 4 | 63733 | 1.8 | 0.678 | 1.681 | 1.453 | 1.961 | 1.181 | 2.832 | 1.019 | 4.005 | 1.625 |
| gel1 | 5 | 63733 | 1.983 | 0.739 | 1.842 | 1.608 | 2.139 | 1.329 | 3.131 | 1.17 | 4.415 | 1.675 |

**Table S7.** Pooled k-nearest neighbor (kNN) distance statistics (k = 1–5) across all regions from gel 2 (see **Supplementary Figure 19B.ii**).

Atto647N to Fluorescein
| subset | k (neighbor order) | n (distance pairs) | Mean (μm) | STD (μm) | Median (μm) | P25 (μm) | P75 (μm) | P5 (μm) | P95 (μm) | P1 (μm) | P99 (μm) | Histogram peak (μm) |
| --- | --- | --- | --- | --- | --- | --- | --- | --- | --- | --- | --- | --- |
| gel2 | 1 | 11671 | 1.443 | 1.643 | 1.079 | 0.757 | 1.502 | 0.41 | 3.404 | 0.22 | 9.989 | 0.825 |
| gel2 | 2 | 11671 | 1.981 | 2.065 | 1.479 | 1.163 | 1.966 | 0.803 | 4.533 | 0.608 | 12.921 | 1.225 |
| gel2 | 3 | 11671 | 2.368 | 2.444 | 1.763 | 1.425 | 2.313 | 1.047 | 5.313 | 0.843 | 15.236 | 1.475 |
| gel2 | 4 | 11671 | 2.699 | 2.92 | 2.004 | 1.632 | 2.632 | 1.252 | 5.988 | 1.05 | 16.898 | 1.675 |
| gel2 | 5 | 11671 | 2.981 | 3.213 | 2.212 | 1.811 | 2.909 | 1.431 | 6.686 | 1.207 | 19.129 | 1.925 |

| subset | k (neighbor order) | n (distance pairs) | Mean (μm) | STD (μm) | Median (μm) | P25 (μm) | P75 (μm) | P5 (μm) | P95 (μm) | P1 (μm) | P99 (μm) | Histogram peak (μm) |
| --- | --- | --- | --- | --- | --- | --- | --- | --- | --- | --- | --- | --- |
| gel2 | 1 | 32765 | 1.11 | 0.509 | 1.017 | 0.79 | 1.309 | 0.61 | 1.887 | 0.567 | 2.55 | 0.825 |
| gel2 | 2 | 32765 | 1.478 | 0.628 | 1.383 | 1.119 | 1.706 | 0.835 | 2.341 | 0.693 | 3.266 | 1.225 |
| gel2 | 3 | 32765 | 1.76 | 0.733 | 1.653 | 1.375 | 1.996 | 1.061 | 2.728 | 0.881 | 3.747 | 1.475 |
| gel2 | 4 | 32765 | 1.995 | 0.831 | 1.866 | 1.58 | 2.239 | 1.252 | 3.031 | 1.058 | 4.205 | 1.675 |
| gel2 | 5 | 32765 | 2.205 | 0.91 | 2.063 | 1.756 | 2.464 | 1.413 | 3.323 | 1.232 | 4.608 | 1.925 |

**Table S8.** Pooled k-nearest neighbor (kNN) distance statistics (k = 1–5) across all regions from gel 3 (see **Supplementary Figure 19B.iii**).

Atto647N to Fluorescein
| subset | k (neighbor order) | n (distance pairs) | Mean (μm) | STD (μm) | Median (μm) | P25 (μm) | P75 (μm) | P5 (μm) | P95 (μm) | P1 (μm) | P99 (μm) | Histogram peak (μm) |
| --- | --- | --- | --- | --- | --- | --- | --- | --- | --- | --- | --- | --- |
| gel3 | 1 | 6489 | 1.274 | 0.821 | 1.082 | 0.734 | 1.592 | 0.384 | 2.888 | 0.203 | 4.344 | 0.825 |
| gel3 | 2 | 6489 | 1.896 | 1.026 | 1.638 | 1.219 | 2.274 | 0.817 | 3.958 | 0.612 | 5.745 | 1.425 |
| gel3 | 3 | 6489 | 2.348 | 1.181 | 2.035 | 1.568 | 2.743 | 1.112 | 4.806 | 0.869 | 6.631 | 1.825 |
| gel3 | 4 | 6489 | 2.721 | 1.309 | 2.36 | 1.857 | 3.146 | 1.367 | 5.466 | 1.112 | 7.398 | 2.025 |
| gel3 | 5 | 6489 | 3.052 | 1.428 | 2.653 | 2.113 | 3.506 | 1.598 | 6.118 | 1.317 | 8.041 | 2.275 |

| subset | k (neighbor order) | n (distance pairs) | Mean (μm) | STD (μm) | Median (μm) | P25 (μm) | P75 (μm) | P5 (μm) | P95 (μm) | P1 (μm) | P99 (μm) | Histogram peak (μm) |
| --- | --- | --- | --- | --- | --- | --- | --- | --- | --- | --- | --- | --- |
| gel3 | 1 | 28897 | 1.267 | 0.684 | 1.085 | 0.815 | 1.495 | 0.612 | 2.548 | 0.567 | 3.942 | 0.825 |
| gel3 | 2 | 28897 | 1.768 | 0.872 | 1.557 | 1.203 | 2.055 | 0.862 | 3.463 | 0.707 | 5.089 | 1.425 |
| gel3 | 3 | 28897 | 2.169 | 1.013 | 1.911 | 1.521 | 2.492 | 1.118 | 4.117 | 0.916 | 5.985 | 1.675 |
| gel3 | 4 | 28897 | 2.51 | 1.134 | 2.201 | 1.795 | 2.867 | 1.363 | 4.708 | 1.129 | 6.921 | 1.925 |
| gel3 | 5 | 28897 | 2.81 | 1.24 | 2.458 | 2.025 | 3.209 | 1.571 | 5.232 | 1.329 | 7.704 | 2.075 |

**Table S9:** KS D randomization statistics by kNN rank (1–5) from all gels (see **Supplementary Figure 20**).

**Atto647N to Fluorescein**
| k (neighbor order) | Mean (μm) | STD (μm) | Median (μm) | P25 (μm) | P75 (μm) | P5 (μm) | P95 (μm) | KS tests with p < 0.001 (out of 100) | KS tests with p ≥ 0.001 (out of 100) |
| --- | --- | --- | --- | --- | --- | --- | --- | --- | --- |
| 1 | 0.089 | 0.003 | 0.089 | 0.087 | 0.091 | 0.085 | 0.094 | 100 | 0 |
| 2 | 0.113 | 0.003 | 0.113 | 0.112 | 0.115 | 0.108 | 0.118 | 100 | 0 |
| 3 | 0.129 | 0.003 | 0.129 | 0.127 | 0.13 | 0.124 | 0.133 | 100 | 0 |
| 4 | 0.144 | 0.002 | 0.144 | 0.142 | 0.145 | 0.141 | 0.148 | 100 | 0 |
| 5 | 0.153 | 0.002 | 0.153 | 0.152 | 0.155 | 0.15 | 0.157 | 100 | 0 |

| k (neighbor order) | Mean (μm) | STD (μm) | Median (μm) | P25 (μm) | P75 (μm) | P5 (μm) | P95 (μm) | KS tests with p < 0.001 (out of 100) | KS tests with p ≥ 0.001 (out of 100) |
| --- | --- | --- | --- | --- | --- | --- | --- | --- | --- |
| 1 | 0.135 | 0.001 | 0.135 | 0.134 | 0.136 | 0.133 | 0.137 | 100 | 0 |
| 2 | 0.193 | 0.002 | 0.193 | 0.192 | 0.194 | 0.19 | 0.196 | 100 | 0 |
| 3 | 0.227 | 0.002 | 0.227 | 0.225 | 0.228 | 0.224 | 0.229 | 100 | 0 |
| 4 | 0.249 | 0.002 | 0.249 | 0.248 | 0.25 | 0.247 | 0.252 | 100 | 0 |
| 5 | 0.264 | 0.002 | 0.264 | 0.263 | 0.265 | 0.261 | 0.267 | 100 | 0 |

**Table S10:**
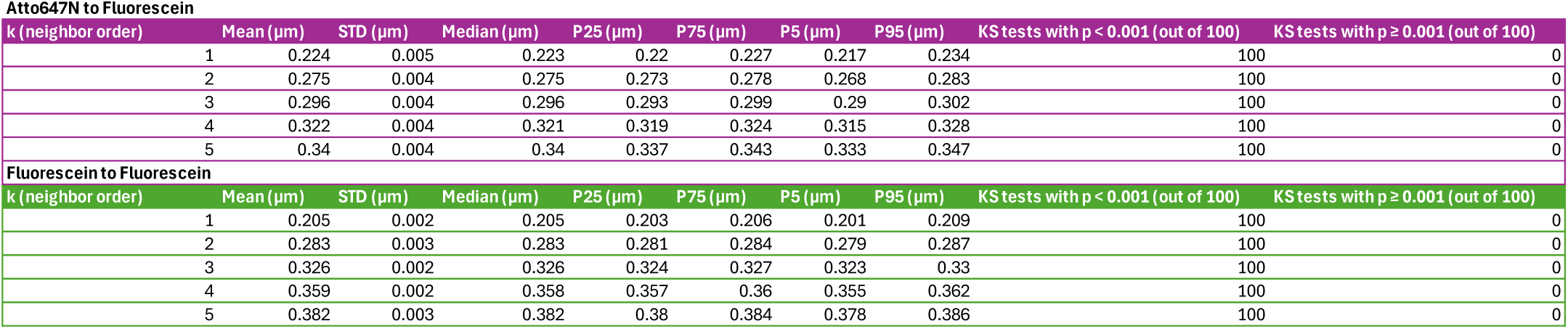
KS D randomization statistics by kNN rank (1–5) from gel 1 (see **Supplementary Figure 21**).

**Table S11:**
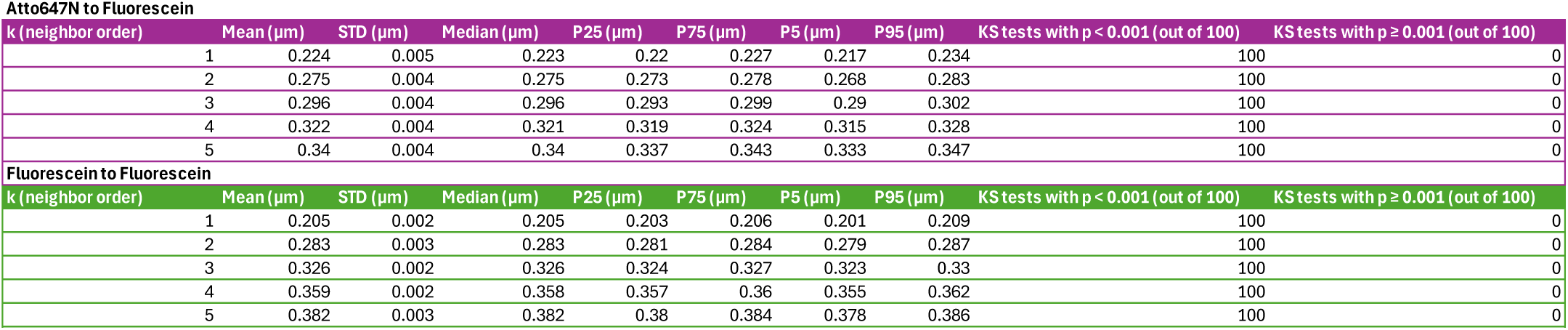
KS D randomization statistics by kNN rank (1–5) from gel 2 (see **Supplementary Figure 22**).

**Table S12:**
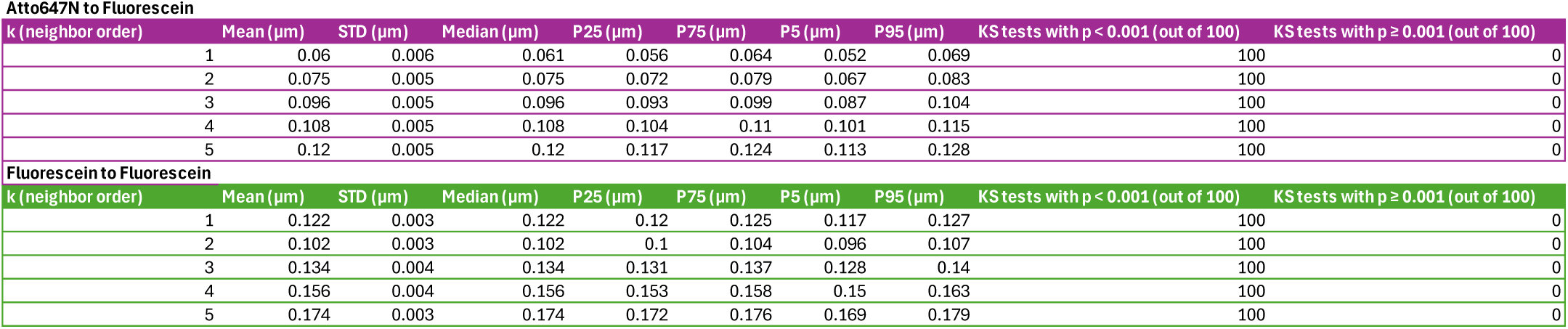
KS D randomization statistics by kNN rank (1–5) from gel 3 (see **Supplementary Figure 23**).

**Table S13:** Summary statistics of single-molecule expansion factors (**Supplementary Figure 11A**), per-axis RMSD (**Supplementary Figure 10A**), and total RMSD (see **Supplementary Figure 9A**, boxplots; **Supplementary Figure 12A**, histograms) for all putative peptides pooled across all gels and regions.

| All Gels | n (molecules) | Mean | STD | Median | P25 | P75 | P10 | P90 |
| --- | --- | --- | --- | --- | --- | --- | --- | --- |
| Single-molecule expansion factor | 6284 | 1055.92 | 21.554 | 1058.372 | 1043.009 | 1071.639 | 1027.077 | 1081.537 |
| RMSD (x-axis, nm) | 6284 | 0.317 | 0.121 | 0.308 | 0.228 | 0.392 | 0.17 | 0.475 |
| RMSD (y-axis, nm) | 6284 | 0.316 | 0.12 | 0.306 | 0.228 | 0.393 | 0.166 | 0.475 |
| RMSD (z-axis, nm) | 6284 | 0.325 | 0.126 | 0.312 | 0.232 | 0.404 | 0.174 | 0.493 |
| Total RMSD (nm) | 6284 | 0.573 | 0.151 | 0.572 | 0.476 | 0.668 | 0.372 | 0.764 |

**Table S14:** Summary statistics of single-molecule expansion factors (**Supplementary Figure 11B**), per-axis RMSD (**Supplementary Figure 10B**), and total RMSD (see **Supplementary Figure 9A**, boxplots; **Supplementary Figure 12A**, histograms) for all putative peptides, stratified by gel (Gel 1-3).

|  | n (molecules) | Mean | STD | Median | P25 | P75 | P10 | P90 |
| --- | --- | --- | --- | --- | --- | --- | --- | --- |
| **Gel 1** |  |  |  |  |  |  |  |  |
| Single-molecule expansion factor | 3155 | 1054.678 | 20.745 | 1057.095 | 1042.238 | 1069.63 | 1026.919 | 1079.035 |
| RMSD (x-axis, nm) | 3155 | 0.313 | 0.117 | 0.305 | 0.225 | 0.386 | 0.17 | 0.466 |
| RMSD (y-axis, nm) | 3155 | 0.314 | 0.117 | 0.305 | 0.23 | 0.387 | 0.17 | 0.471 |
| RMSD (z-axis, nm) | 3155 | 0.318 | 0.118 | 0.308 | 0.231 | 0.393 | 0.175 | 0.474 |
| Total RMSD (nm) | 3155 | 0.565 | 0.14 | 0.567 | 0.476 | 0.655 | 0.378 | 0.742 |
| **Gel 2** |  |  |  |  |  |  |  |  |
| Single-molecule expansion factor | 1918 | 1055.137 | 22.519 | 1057.447 | 1041.44 | 1071.891 | 1025.949 | 1081.618 |
| RMSD (x-axis, nm) | 1918 | 0.313 | 0.123 | 0.303 | 0.225 | 0.393 | 0.164 | 0.48 |
| RMSD (y-axis, nm) | 1918 | 0.31 | 0.12 | 0.304 | 0.219 | 0.389 | 0.156 | 0.469 |
| RMSD (z-axis, nm) | 1918 | 0.322 | 0.13 | 0.305 | 0.225 | 0.405 | 0.166 | 0.493 |
| Total RMSD (nm) | 1918 | 0.565 | 0.16 | 0.562 | 0.461 | 0.665 | 0.352 | 0.772 |
| **Gel 3** |  |  |  |  |  |  |  |  |
| Single-molecule expansion factor | 1211 | 1060.396 | 21.496 | 1063.804 | 1047.249 | 1076.174 | 1030.608 | 1085.43 |
| RMSD (x-axis, nm) | 1211 | 0.331 | 0.124 | 0.32 | 0.241 | 0.405 | 0.184 | 0.496 |
| RMSD (y-axis, nm) | 1211 | 0.329 | 0.129 | 0.315 | 0.231 | 0.414 | 0.17 | 0.501 |
| RMSD (z-axis, nm) | 1211 | 0.349 | 0.137 | 0.332 | 0.251 | 0.444 | 0.182 | 0.53 |
| Total RMSD (nm) | 1211 | 0.604 | 0.158 | 0.6 | 0.5 | 0.708 | 0.401 | 0.811 |

**Table S15:** Summary statistics of single-molecule expansion factors (**Supplementary Figure 11C**), per-axis RMSD (**Supplementary Figure 10C**), and total RMSD (see **Supplementary Figure 9C**, boxplots; **Supplementary Figure 12C**, histograms) for all putative peptides, stratified by imaging region across Gel 1-3 (n=8 regions).

|  | n (molecules) | Mean | STD | Median | P25 | P75 | P10 | P90 |
| --- | --- | --- | --- | --- | --- | --- | --- | --- |
| **Gel 1, Region 1** |  |  |  |  |  |  |  |  |
| Single-molecule expansion factor | 1531 | 1055.032 | 20.607 | 1057.373 | 1042.318 | 1070.055 | 1027.387 | 1079.503 |
| RMSD (x-axis, nm) | 1531 | 0.315 | 0.119 | 0.306 | 0.226 | 0.387 | 0.171 | 0.474 |
| RMSD (y-axis, nm) | 1531 | 0.314 | 0.116 | 0.305 | 0.23 | 0.389 | 0.169 | 0.472 |
| RMSD (z-axis, nm) | 1531 | 0.32 | 0.119 | 0.309 | 0.231 | 0.397 | 0.176 | 0.476 |
| Total RMSD (nm) | 1531 | 0.568 | 0.14 | 0.568 | 0.477 | 0.658 | 0.385 | 0.746 |
| **Gel 1, Region 2** |  |  |  |  |  |  |  |  |
| Single-molecule expansion factor | 1624 | 1054.344 | 20.876 | 1056.945 | 1042.213 | 1069.255 | 1026.531 | 1078.649 |
| RMSD (x-axis, nm) | 1624 | 0.312 | 0.115 | 0.304 | 0.225 | 0.385 | 0.168 | 0.461 |
| RMSD (y-axis, nm) | 1624 | 0.314 | 0.117 | 0.304 | 0.231 | 0.386 | 0.171 | 0.47 |
| RMSD (z-axis, nm) | 1624 | 0.317 | 0.117 | 0.308 | 0.232 | 0.392 | 0.173 | 0.472 |
| Total RMSD (nm) | 1624 | 0.563 | 0.14 | 0.566 | 0.475 | 0.65 | 0.369 | 0.741 |
| **Gel 2, Region 1** |  |  |  |  |  |  |  |  |
| Single-molecule expansion factor | 523 | 1053.636 | 22.632 | 1055.618 | 1037.743 | 1071.221 | 1024.864 | 1081.227 |
| RMSD (x-axis, nm) | 523 | 0.301 | 0.116 | 0.293 | 0.219 | 0.369 | 0.16 | 0.466 |
| RMSD (y-axis, nm) | 523 | 0.303 | 0.117 | 0.292 | 0.217 | 0.382 | 0.151 | 0.452 |
| RMSD (z-axis, nm) | 523 | 0.322 | 0.137 | 0.306 | 0.222 | 0.41 | 0.165 | 0.512 |
| Total RMSD (nm) | 523 | 0.553 | 0.161 | 0.552 | 0.447 | 0.663 | 0.34 | 0.766 |
| **Gel 2, Region 2** |  |  |  |  |  |  |  |  |
| Single-molecule expansion factor | 849 | 1050.75 | 21.995 | 1052.864 | 1037.705 | 1065.365 | 1022.149 | 1077.294 |
| RMSD (x-axis, nm) | 849 | 0.296 | 0.112 | 0.289 | 0.217 | 0.367 | 0.159 | 0.445 |
| RMSD (y-axis, nm) | 849 | 0.294 | 0.114 | 0.292 | 0.21 | 0.369 | 0.15 | 0.438 |
| RMSD (z-axis, nm) | 849 | 0.302 | 0.121 | 0.287 | 0.21 | 0.378 | 0.156 | 0.462 |
| Total RMSD (nm) | 849 | 0.533 | 0.146 | 0.537 | 0.439 | 0.624 | 0.332 | 0.716 |
| **Gel 2, Region 3** |  |  |  |  |  |  |  |  |
| Single-molecule expansion factor | 497 | 1062.861 | 21.276 | 1065.479 | 1050.542 | 1077.634 | 1034.341 | 1088.583 |
| RMSD (x-axis, nm) | 497 | 0.35 | 0.139 | 0.334 | 0.25 | 0.437 | 0.184 | 0.533 |
| RMSD (y-axis, nm) | 497 | 0.338 | 0.127 | 0.33 | 0.245 | 0.419 | 0.176 | 0.505 |
| RMSD (z-axis, nm) | 497 | 0.348 | 0.133 | 0.326 | 0.251 | 0.433 | 0.193 | 0.52 |
| Total RMSD (nm) | 497 | 0.619 | 0.166 | 0.621 | 0.509 | 0.722 | 0.401 | 0.85 |
| **Gel 2, Region 4** |  |  |  |  |  |  |  |  |
| Single-molecule expansion factor | 49 | 1068.823 | 16.836 | 1070.129 | 1058.476 | 1079.936 | 1043.815 | 1088.267 |
| RMSD (x-axis, nm) | 49 | 0.366 | 0.135 | 0.363 | 0.257 | 0.46 | 0.189 | 0.532 |
| RMSD (y-axis, nm) | 49 | 0.376 | 0.129 | 0.385 | 0.27 | 0.457 | 0.207 | 0.53 |
| RMSD (z-axis, nm) | 49 | 0.383 | 0.132 | 0.385 | 0.296 | 0.443 | 0.235 | 0.534 |
| Total RMSD (nm) | 49 | 0.675 | 0.137 | 0.643 | 0.589 | 0.727 | 0.547 | 0.881 |
| **Gel 3, Region 1** |  |  |  |  |  |  |  |  |
| Single-molecule expansion factor | 153 | 1063.514 | 22.452 | 1070.043 | 1050.811 | 1079.503 | 1031.755 | 1086.738 |
| RMSD (x-axis, nm) | 153 | 0.337 | 0.121 | 0.313 | 0.257 | 0.417 | 0.192 | 0.501 |
| RMSD (y-axis, nm) | 153 | 0.344 | 0.139 | 0.345 | 0.241 | 0.435 | 0.173 | 0.517 |
| RMSD (z-axis, nm) | 153 | 0.376 | 0.151 | 0.358 | 0.261 | 0.48 | 0.191 | 0.579 |
| Total RMSD (nm) | 153 | 0.635 | 0.164 | 0.656 | 0.525 | 0.744 | 0.416 | 0.839 |
| **Gel 3, Region 2** |  |  |  |  |  |  |  |  |
| Single-molecule expansion factor | 1058 | 1059.946 | 21.327 | 1063.189 | 1046.846 | 1075.58 | 1030.574 | 1085.255 |
| RMSD (x-axis, nm) | 1058 | 0.33 | 0.124 | 0.321 | 0.24 | 0.403 | 0.18 | 0.492 |
| RMSD (y-axis, nm) | 1058 | 0.326 | 0.127 | 0.313 | 0.231 | 0.409 | 0.169 | 0.498 |
| RMSD (z-axis, nm) | 1058 | 0.345 | 0.135 | 0.329 | 0.251 | 0.437 | 0.179 | 0.521 |
| Total RMSD (nm) | 1058 | 0.6 | 0.157 | 0.596 | 0.499 | 0.701 | 0.399 | 0.801 |

**Table S16.**
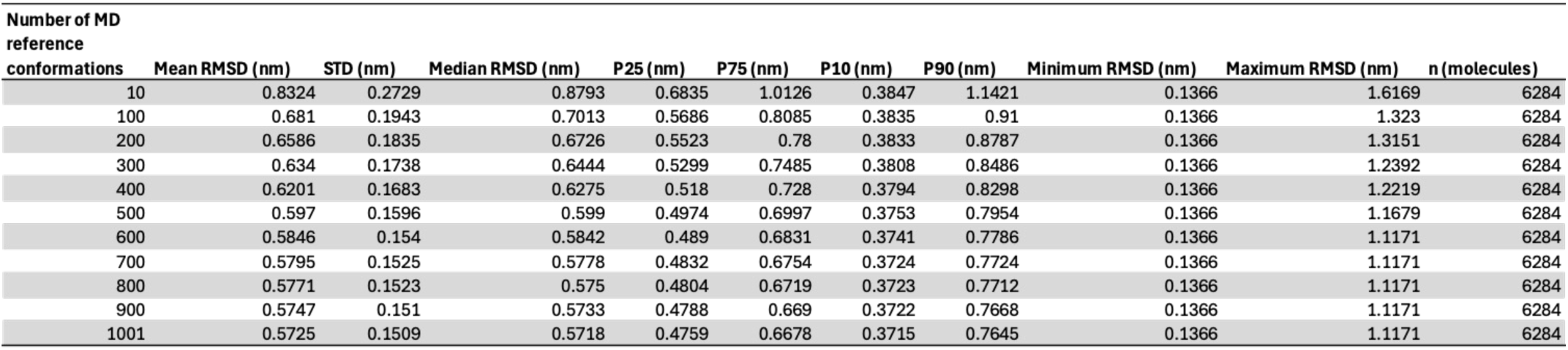
Summary statistics of total RMSD distributions as a function of the number of molecular-dynamics (MD) reference conformations (see **Supplementary Figure 13**, boxplots; **Supplementary Figure 14**, histograms).

**Table S17:** Statistics elements for **Fig. 2**.

|  |  |  |  |  |  |  |  |
| --- | --- | --- | --- | --- | --- | --- | --- |
| **Panel B** | 18x (ONE) | 50x (ONE) | 100x (ONE) | 100x (confocal) | 500x (confocal) | 1000x (confocal) | model |
| Mean | 4.434317761 | 5.210717151 | 5.157849609 | 6.177624755 | 3.900797385 | 7.090570252 | 4.88207877 |
| Median | 4.220428333 | 4.269811 | 4.222855 | 5.460546 | 2.408406 | 5.822544 | 4.959 |
| Std deviation | 1.991380873 | 3.786436472 | 3.851486303 | 3.813227427 | 4.826060053 | 6.186493755 | 1.419420211 |
| Std error | 0.02054499 | 0.122462107 | 0.1289573 | 6.67E-03 | 0.037042528 | 0.22819226 | 0.044886008 |
| % CV | 44.90839359 | 72.66632139 | 74.67232655 | 61.72643336 | 123.7198341 | 87.24959396 | 29.07409482 |
| 90% conf | 0.033799843 | 0.201645797 | 0.212355577 | 0.010972448 | 0.060938286 | 0.375850696 | 0.073906027 |
| 95% conf | 0.04027337 | 0.240330327 | 0.253100129 | 0.013073567 | 0.072608534 | 0.447995574 | 0.088083297 |
| 99% conf | 0.052934652 | 0.316094153 | 0.332907159 | 0.017182438 | 0.095432286 | 0.589355788 | 0.1158477 |
| N | 9395 | 956 | 892 | 326824 | 16974 | 735 | 1000 |
| **Panel F** |  |  |  |  |  |  |  |
| R | 0.9255 |  | Coefficient | Std. Error | t | P | VIF |
| R2 | 0.8566 | y0 | 0.2809 | 0.618 | 0.4545 | 0.6685 | 4.7978< |
| Adj R2 | 0.8279 | a | 0.809 | 0.148 | 5.4646 | 0.0028 | 4.7978< |
| Standard error of estimate | 0.7465 |  |  |  |  |  |  |
| **Panel G** |  |  |  |  |  |  |  |
| R | 0.925 |  | Coefficient | Std. Error | t | P | VIF |
| R2 | 8556 |  |  |  |  |  |  |
| Adj R2 | 0.8483 | y0 | 0.2613 | 0.1059 | 2.4682 | 0.0227 | 3.0004 |
| Standard error of estimate | 0.3055 | a | 0.699 | 0.0569 | 12.2836 | <0.0001 | 3.0004 |
| **Panel H** |  |  |  | **Panel I** |  |  |  |
| Mean | 1.518872 | 1.161491304 |  | Mean | 0.655143478 | 0.568928571 |  |
| Median | 1.524 | 1.2634 |  | Median | 0.606 | 0.549 |  |
| Std deviation | 1.096045975 | 0.79576071 |  | Std deviation | 0.255672923 | 0.06306863 |  |
| Std error | 0.219209195 | 0.165927578 |  | Std error | 0.05331149 | 0.013762703 |  |
| % CV | 72.16183952 | 68.51198169 |  | % CV | 39.02548546 | 11.08550943 |  |
| 90% conf | 0.375076408 | 0.284948583 |  | 90% conf | 0.091552192 | 0.023739047 |  |
| 95% conf | 0.45243455 | 0.344119637 |  | 95% conf | 0.11056348 | 0.028709077 |  |
| 99% conf | 0.613161459 | 0.467745354 |  | 99% conf | 0.150283647 | 0.039162624 |  |
| N | 25 | 23 |  | N | 23 | 21 |  |
| Mann-Whitney Ranksum test Pvalue |  | 0.2156 |  | Mann-Whitney Ranksum test Pvalue |  | 0.3827 |  |

**Table S18.** Measured gel lengths and calculated shrinkage and expansion factors during four-network gel formation. For each round, the table reports lengths before and after expansion, per-round expansion factors, and cumulative linear expansion factors. For rounds 2–4, the table also reports lengths before monomer addition and after gelation, which were used to calculate per-round shrinkage factors. Cumulative linear expansion factors were calculated by multiplying all sequential shrinkage and expansion factors through the indicated round.

| | | | | Round 1<br>before<br>expansion<br>length (cm) | Round 1 after<br>expansion<br>length (cm) | Round 1<br>expansion<br>factor ($E_1 = C_1$) |
| --- | --- | --- | --- | --- | --- | --- |
|  |  |  |  | 1 | 15.8 | 15.8 |
|  |  |  |  | 1 | 15.8 | 15.8 |
|  |  |  |  | 1 | 17.5 | 17.5 |
|  |  |  |  | 1 | 16.7 | 16.7 |
|  |  |  |  | 1 | 17.1 | 17.1 |
|  |  |  |  | 1 | 18.7 | 18.7 |
| Round 2<br>before<br>monomer<br>addition<br>length (cm) | Round 2 after<br>gelation<br>length (cm) | Round 2<br>shrinkage<br>factor ($S_2$) | Round 2<br>before<br>expansion<br>length (cm) | Round 2 after<br>expansion<br>length (cm) | Round 2<br>expansion<br>factor ($E_2$) | Round 2<br>cumulative<br>expansion<br>factor ($C_2$) |
| 2.9 | 1.25 | 0.43 | 1 | 9 | 9.0 | 61.1 |
| 2.3 | 1.1 | 0.48 | 1 | 9 | 9.0 | 67.8 |
| 2.9 | 1.25 | 0.43 | 0.85 | 9 | 10.6 | 79.9 |
| 2.7 | 1.2 | 0.44 | 0.8 | 9.45 | 11.8 | 87.5 |
| 2.9 | 1.5 | 0.52 | 0.9 | 8.7 | 9.7 | 85.7 |
| 2.9 | 1.6 | 0.55 | 1 | 9.6 | 9.6 | 98.9 |
| Round 3<br>before<br>monomer<br>addition<br>length (cm) | Round 3 after<br>gelation<br>length (cm) | Round 3<br>shrinkage<br>factor ($S_3$) | Round 3<br>before<br>expansion<br>length (cm) | Round 3 after<br>expansion<br>length (cm) | Round 3<br>expansion<br>factor ($E_3$) | Round 3<br>cumulative<br>expansion<br>factor ($C_3$) |
| 1.4 | 0.6 | 0.43 | 1.2 | 11 | 9.6 | 250.5 |
| 1.9 | 1 | 0.53 | 1.2 | 11 | 9.6 | 341.3 |
| 2.2 | 1 | 0.45 | 1.1 | 10 | 9.1 | 330.0 |
| 2.9 | 1.6 | 0.55 | 0.9 | 9.2 | 10.2 | 493.6 |
| 1.4 | 0.65 | 0.46 | 0.9 | 8.7 | 9.7 | 384.6 |
| 2 | 1.05 | 0.53 | 1.0 | 9.4 | 9.4 | 488.0 |
| Round 4<br>before<br>monomer<br>addition<br>length (cm) | Round 4 after<br>gelation<br>length (cm) | Round 4<br>shrinkage<br>factor ($S_4$) | Round 4<br>before<br>expansion<br>length (cm) | Round 4 after<br>expansion<br>length (cm) | Round 4<br>expansion<br>factor ($E_4$) | Round 4<br>cumulative<br>expansion<br>factor ($C_4$) |
| 2.2 | 1.45 | 0.66 | 1.25 | 9.3 | 7.44 | 1228.2 |
| 2.1 | 1.35 | 0.64 | 1.15 | 10.3 | 8.96 | 1965.1 |
| 2.8 | 1.6 | 0.57 | 1.2 | 10.6 | 8.83 | 1665.9 |
| 2.6 | 1.5 | 0.58 | 1.15 | 9 | 7.83 | 2228.6 |
| 2.6 | 1.45 | 0.56 | 1.05 | 9 | 8.57 | 1838.6 |
| 2.5 | 1.4 | 0.56 | 1.1 | 10.2 | 9.27 | 2534.1 |

## Supplementary Notes

### Supplementary Note 1: 1000ExM Protocol, Updated

#### Activated monomer solution preparation

| Reagent | Amount in 620 μL Activated Monomer Solution (mol) | Concentration (M) | Percentage |
| --- | --- | --- | --- |
| Sodium Acrylate (AK Scientific, R624) | $1.69 \times 10^{-3}$ mol | 2.73 M | 25.6% (w/v) |
| N,N-Dimethylacrylamide (DMAA; ThermoFisher, 432021000) | $2.60 \times 10^{-3}$ mol | 4.19 M | 44.2% (v/v) |
| N,N,N',N'-Tetramethylethylenediamine (TEMED; Sigma, T9281) | $1.52 \times 10^{-6}$ mol | 2.45 mM | 0.037% (v/v) |
| Potassium persulfate (KPS; Sigma, 216224) | $3.33 \times 10^{-6}$ mol | 5.37 mM | 0.145% (w/v) |

1. Weigh 3.14 g sodium acrylate into a 50 mL Eppendorf tube and dissolve it in 6 mL acidified Tris buffer, prepared from: 1 M Tris pH 8.0 (ThermoFisher, #AM9856), 32% HCl solution (Sigma, #W530574), and water. (To make 50 mL of acidified Tris, add 40 mL water, then 5.00 mL 1 M Tris pH 8.0, then add 1.18 mL 32% HCl, and bring the final volume to 50.0 mL with water.) Vortex the acrylate solution for approximately 5 to 10 minutes, until the solution is completely clear.
2. Add 5.4 mL N,N-dimethylacrylamide to the sodium acrylate solution and vortex for less than 1 minute.
3. Prepare a 10% N,N,N′,N′-tetramethylethylenediamine solution in water. Add 45 µL of this solution to the above mixture and mix again.
4. Check that the solution is clear and free of precipitate. If precipitate is present (possible for some batches of sodium acrylate), filter the solution using a syringe filter (Sigma, #SLGSR33SS) and collect the clear filtrate.
5. Degas the solution with nitrogen gas at room temperature (∼20°C) using a Pasteur pipette connected to a nitrogen source (Sigma, #BR747725). Place the tip of the pipette at the bottom of the 50 mL tube and bubble for 20 minutes at a low-to-moderate flow rate sufficient to produce steady bubbling without splashing.
6. Pipette 600 µL of the degassed monomer solution into an Eppendorf tube. Add 20 µL potassium persulfate solution (45 mg/mL stock in water) to activate polymerization (which will proceed slowly). The resulting mixture is the activated monomer solution.

Note: These volumes yield enough activated monomer solution for 9 iterative gels (620 µL per gel, applied twice, 1240 µL total per gel), incubated in 12-well plastic plates (do not use glass plates, as this seems to cause premature gelation).

#### To make the first gel

7. Prepare the sealable plastic container (Rubbermaid 9.6 Cup Brilliance Food Storage Container) used to hold the gel chamber. Modify the lid to contain two small holes: one inlet and one outlet. Confirm the container does not leak by sealing both holes with tape, adding water to the container, and checking whether any water escapes; if leaks are found, do not proceed.
8. Use the lid of the 12-well plate as a small platform to hold the hydrophobic microscope slides during gelation. Add 30 to 50 µL of activated monomer solution onto a hydrophobic microscope slide (CytoSlide Fluorosilane, CYTONIX); the volume used depends on the desired gel thickness. Place a 1.2 mm coverslip with the attached sample (proteins or cells) on top of the monomer solution, biological sample face down, with no spacer. Together, the slide, monomer solution, and coverslip form the gel chamber. Place the gel chamber, still resting on the lid platform, inside the prepared container. Insert a pipette tip, connected via rubber tubing to a nitrogen line, into the inlet hole to deliver nitrogen gas, and leave the second hole open as a vent to allow air to escape as nitrogen fills the container.
9. Purge the container with nitrogen on high for 20 minutes at room temperature to displace oxygen, making sure the nitrogen stream is not blowing directly at the gel. No supplemental moisture was added to the Tupperware in this or any other step. After purging, remove the nitrogen line and seal both holes with tape to maintain a nitrogen atmosphere. Allow the gel to polymerize overnight (8-16 hours) at room temperature.
10. Immerse the gel in excess deionized water at room temperature (∼20 °C) for 3 hours to remove unreacted components and allow expansion. Replace the water at least twice, continuing exchanges as needed until full linear expansion of approximately 15-18× is reached.

To form the next network within an already-expanded gel (all steps, unless otherwise noted, are performed at room temperature):

11. Cut each expanded gel into pieces up to 2 cm in length and approximately 1 cm in width, on the lid of a 12-well plate (see photo below), with an asymmetric corner cut to mark which side the sample is on. Then, place each gel piece into an individual well of a 12-well plate.

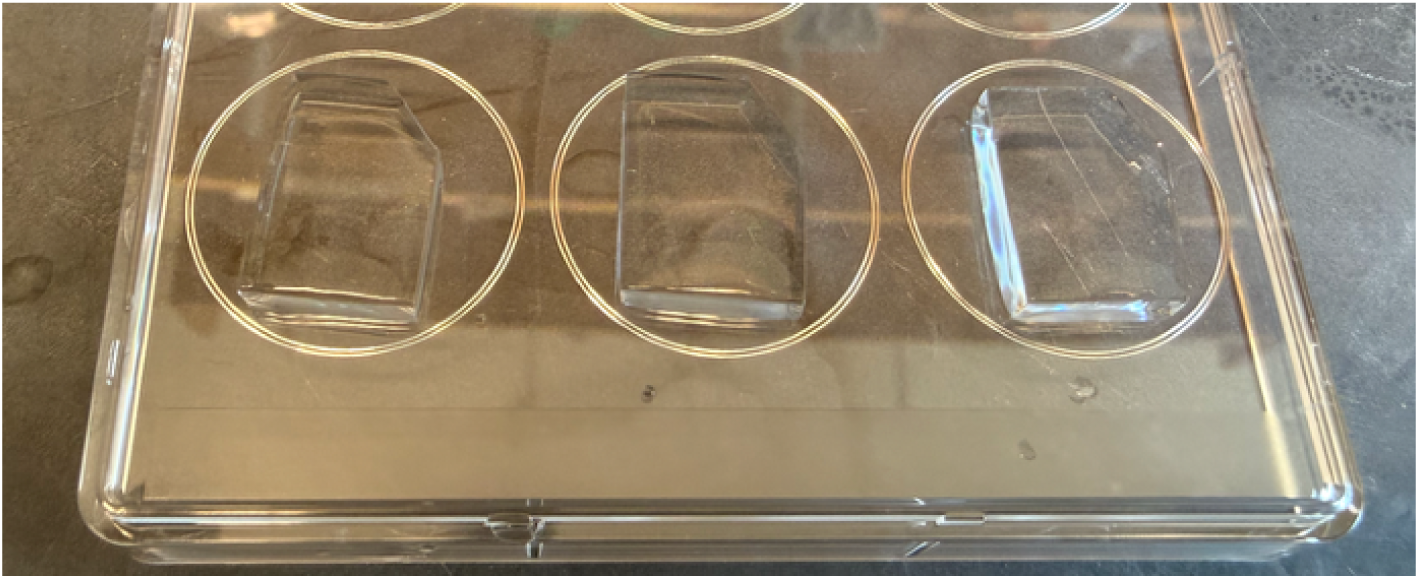
12. Add 600 µL degassed monomer solution and 20 µL potassium persulfate solution (45 mg/mL stock) to each well to make the activated monomer solution. Position the plate at approximately a 45° angle in a Tupperware container so that the gels are fully immersed from the top, bottom, and sides (critical). Insert a pipette tip, connected via rubber tubing to a nitrogen line, into the inlet hole to deliver nitrogen gas, and leave the second hole open as a vent to allow air to escape as nitrogen fills the container. Leave the nitrogen on and shake for 20 minutes.
13. Prepare a new 12-well plate with 600 µL monomer solution and 20 µL KPS solution in each well. Transfer the gels into the new solution, keeping the nitrogen on, and shake for another 20 minutes.
14. Remove the gels quickly. Place each gel onto a hydrophobic glass slide with the sample facing down, and quickly cover it with a coverslip 12mm in diameter. No spacer and no backfilling were used.
15. Purge the Tupperware with nitrogen gas for 1 hour, then allow the gel to polymerize overnight (≥8 hours).
16. After gelation is complete, measure the linear dimensions of each gel. The shrinkage factor for each round can be determined by measuring gel size after polymerization, which typically shrinks from 2 cm to approximately 1 to 1.2 cm. Example:

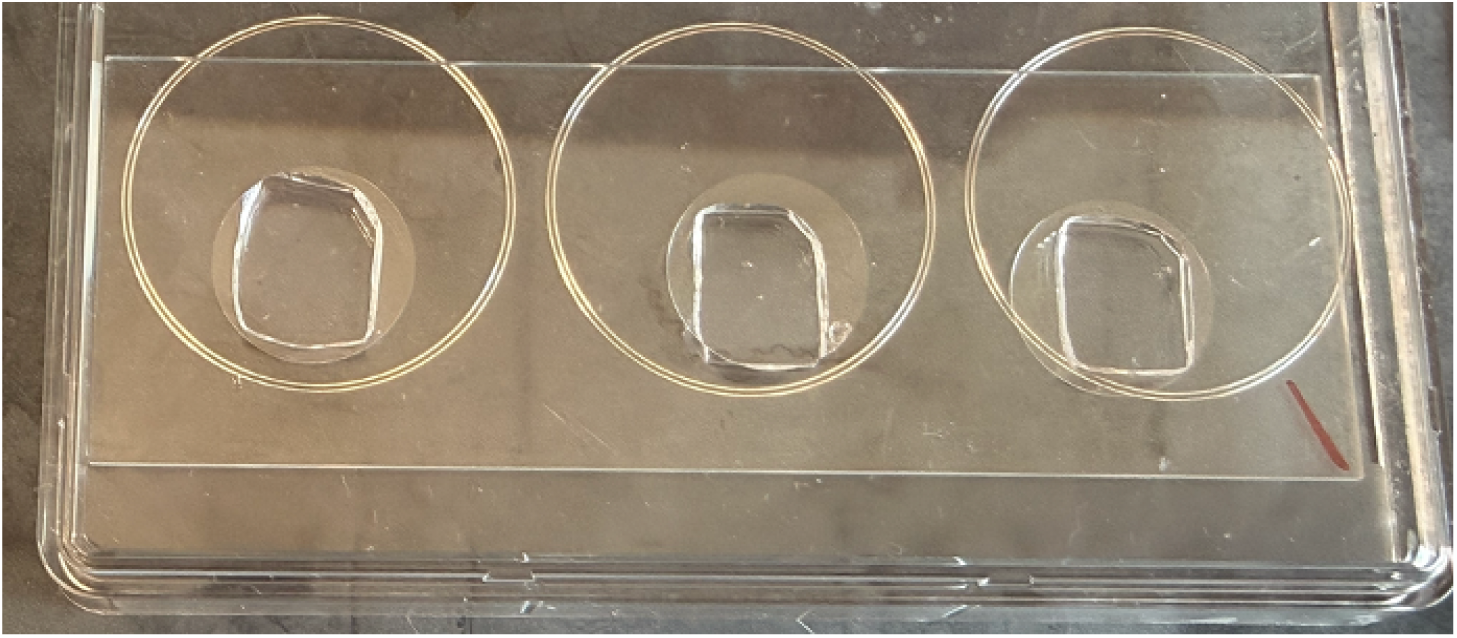
17. Use a paintbrush dipped in 1x PBS to detach the gel from the glass, then cut away the edges of the gel.

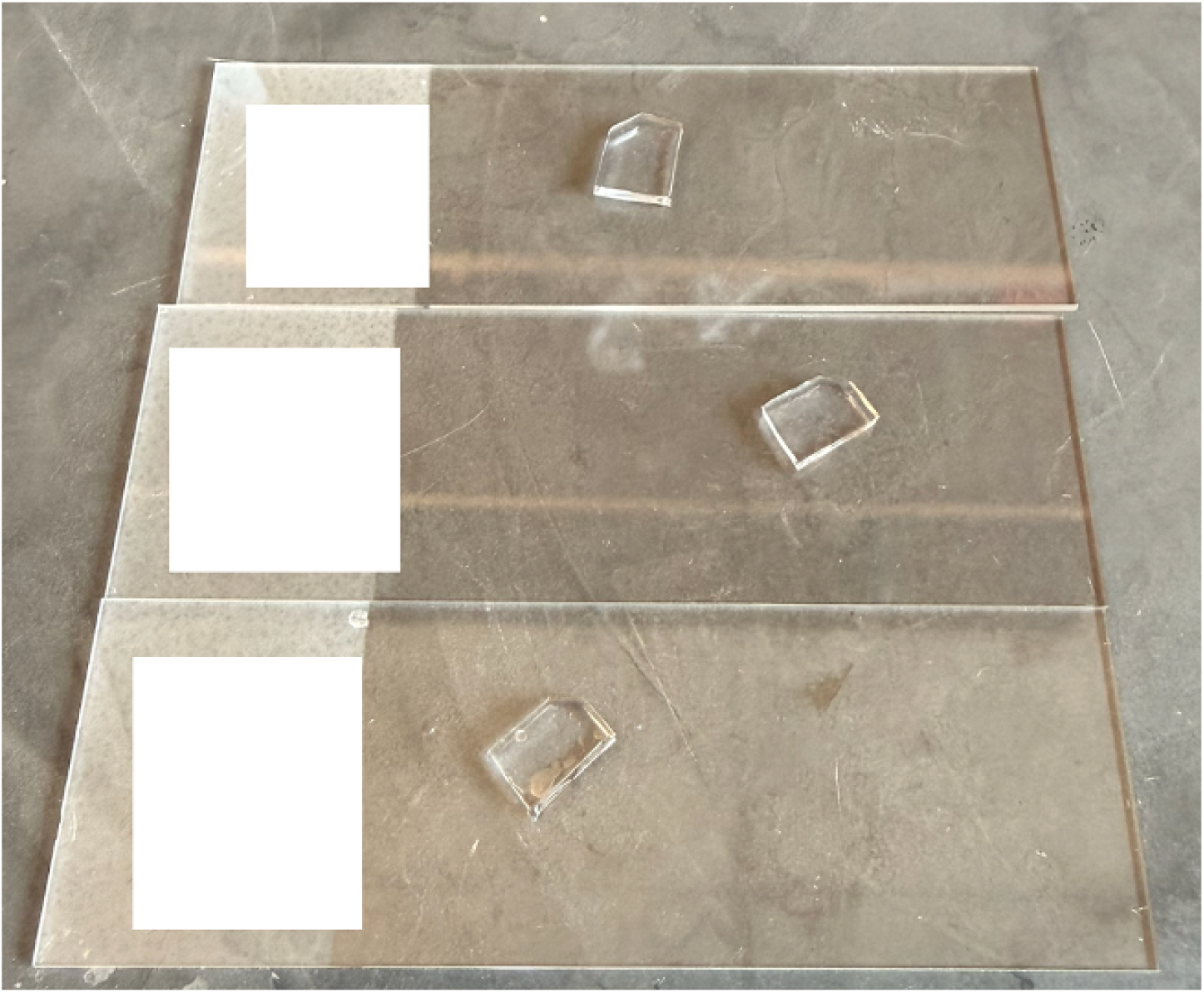
18. Place each gel piece (less than 1 cm in length) in a 140 mm petri dish and incubate in water, filled to the top of the dish, until expansion reaches equilibrium, typically 24 to 48 hours. Measure the final length and height of the gel to determine the expansion factor for each round. The gel typically expands approximately 10-fold (for example, from 0.8 cm to 8 cm). No intermediate water exchanges are required.

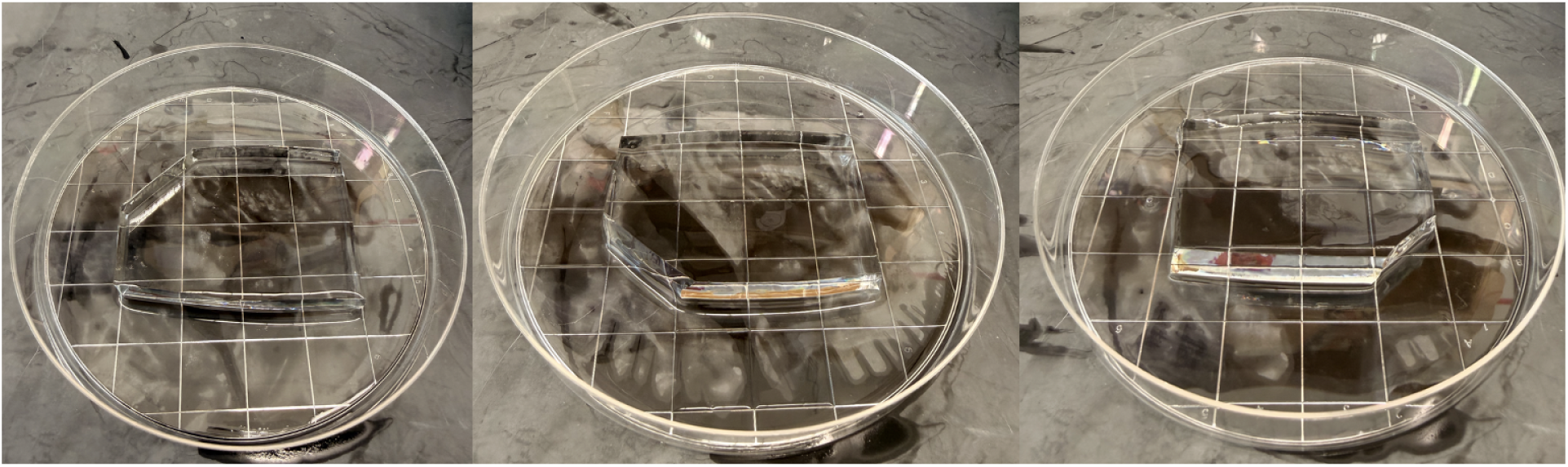
19. After round 2 (typically 60 to 100×), axially section the gel, sample side facing down, to generate thinner gel pieces for the next casting round. To prepare a slicing guide, glue (using superglue) three microscope slides together to make a 3 mm thick spacer, and prepare two such spacers. Place one spacer on either side of the gel, rest a long razor blade (Leica microtome blades, Electron Microscopy Sciences #63065-LP) on top of the spacers, and slide it through the gel to produce an approximately 3 mm thick axial section. Use these 3 mm thick gel sections for the subsequent iterative gel-casting round. Shown are the sectioning chamber (left) and two pieces of gel, sectioned on (right).

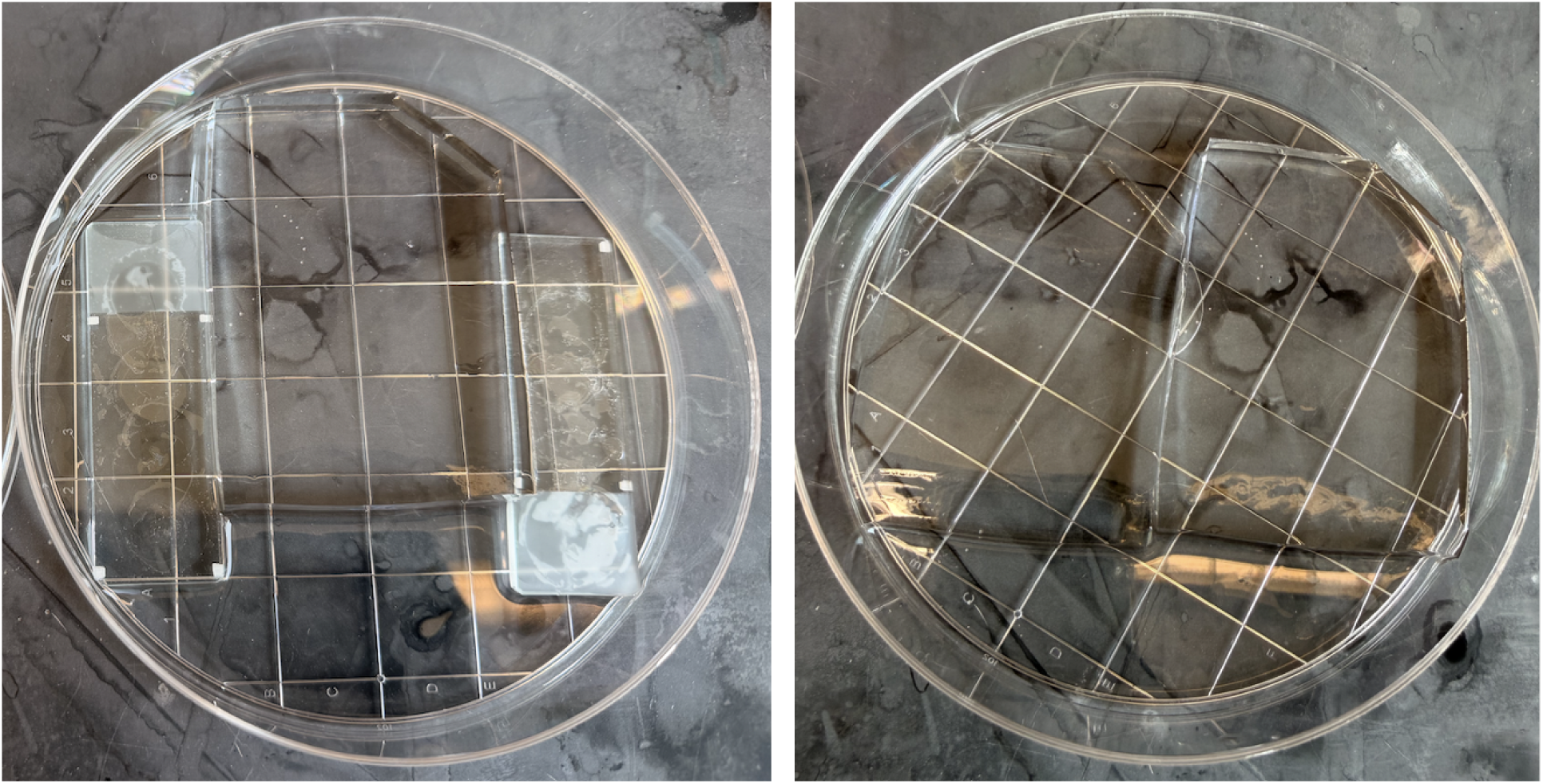
20. Cut the 3 mm thick gel into pieces 2 cm in length to repeat the iterative gelation steps and form the round 3 gels (typically 250 to 500×). Similarly, after axially slicing the fully expanded round 3 gels, cut them into 2 cm pieces to form the round 4 gels (typically 1000 to 2500×).

### Supplementary Note 2: 1000ExM Protocol, Original

To prepare the activated monomer solution:

1. Weigh 2.090 ± 0.002 g of sodium acrylate and dissolve it in 4 mL of acidified Tris buffer prepared as described in Supplementary Note 1.
2. Add 3.600 mL N,N-dimethylacrylamide to the sodium acrylate solution and mix thoroughly.
3. Prepare a 10% solution of N,N,N′,N′-tetramethylethylenediamine (90 µL water + 10 µL pure compound). Add 30 µL of this solution to the above mixture and mix again until fully incorporated.
4. Check if the solution is precipitate free. Otherwise, filter the solution using a syringe filter (Sigma, #SLGSR33SS) and collect the clear supernatant.
5. Bubble the solution with nitrogen gas using a Pasteur pipette directly connected to a nitrogen source (Sigma, #BR747725) at room temperature (∼20 °C) for 15 min using a low-to-moderate flow rate sufficient to produce steady bubbling without splashing.
6. Pipette 900 µL of the degassed monomer solution into an Eppendorf tube.
7. Add 30 µL potassium persulfate solution (45 mg/mL stock in water). The resulting solution is the activated monomer solution.

Note: These volumes yield enough activated monomer solution for 6 iterative gels (930 µL per gel), incubated in 6 well plastic plates.

To make the first gel:

8. Add 50 µL activated monomer solution onto a hydrophobic microscope slide (CytoSlide Fluorosilane, CYTONIX) in a Tupperware container.
9. Place a coverslip with the attached sample (proteins or cells) on top, with the biological sample face down (no spacer).
10. Place the gel chamber inside a sealable plastic container (e.g., Tupperware). Before beginning any procedures, modify the lid to contain two small holes: one inlet and one outlet. Insert a pipette tip (connected via rubber tubing to a nitrogen line) into the inlet hole to deliver nitrogen gas. Leave the second hole open to serve as a vent, allowing air to escape as nitrogen fills the container.
11. Purge with nitrogen for 1 hour to displace oxygen at room temperature. (No supplemental moisture was added to the Tupperware, in this or any other step.) After purging, remove the nitrogen line and seal both holes with tape to maintain a nitrogen atmosphere. Allow the gel to polymerize overnight (≥8 hours) at room temperature.
12. Immerse the gel in excess deionized water at room temperature (∼20 °C) for 3 hours to remove unreacted components and allow expansion. Replace the water at least twice, continuing exchanges as needed until full ∼18× linear expansion is reached.

To make an iterative gel throughout an expanded gel (all steps unless otherwise noted are done at room temperature):

13. Place each expanded gel piece into an individual well of a 6-well plate. Before doing so, cut the gel into pieces up to approximately 3 cm in length and 1.5 cm in width.
14. Add 900 µL degassed monomer solution and 30 µL potassium persulfate solution (45 mg/mL stock) to each well to make the activated monomer solution.
15. Position the plate at approximately 45° angle in a Tupperware container so that gels are fully immersed from the top, bottom, and sides (critical).
16. Degas the Tupperware with nitrogen gas for 10 minutes, then place on a shaker for 35 minutes to incubate.
17. Remove gels quickly. Place each gel onto a hydrophobic glass slide with the sample facing down and quickly cover with a coverslip (within 30 seconds) to minimize oxygen exposure. No spacer, and no backfilling, were performed.
18. Purge the Tupperware with nitrogen gas for 1 hour, then allow the gel to form overnight (≥8 hours).
19. After gelation is complete, measure the linear dimensions of each gel. To determine the shrinkage factor for each round, compare the gel size immediately after transfer to the 6-well plate with the gel size after gelation is complete.
20. Then incubate the gel in water until the expansion reaches equilibrium, typically 24–48 h. Measure the final linear dimensions to determine the expansion factor for each round. Measure the gel along multiple edges, as well as the gel height, to assess expansion in all dimensions.
21. To reach the maximum iterative expansion capacity, perform up to three additional rounds of iterative gel casting after the initial gel. The second, third, and fourth rounds yield approximately 100×, 500×, and 1500× linear expansion, respectively. To fully expand the 100×, 500×, and 1000–1500× gels, place an approximately 1 cm gel piece in a large Petri dish (140 mm diameter; Thermo Fisher, 08-757-100). Add ∼100 mL deionized water, or enough water to completely fill the dish. Incubate at room temperature (∼20 °C) for 24–48 h, until the gel is fully expanded. Water exchange is not required.
22. After the 100× expansion round, axially section the gel to generate thinner gel pieces for the next casting round to reach ∼500× expansion. After the 500× expansion round, repeat the axial sectioning step to generate thinner gel pieces for the next casting round to reach ∼1000–1500× expansion.
23. To prepare a slicing guide, glue three microscope slides together to make a 3 mm-thick spacer. Prepare two spacers and place them on either side of the gel. Place a razor blade on top of the spacers and slide it through the gel to produce an approximately 3 mm-thick axial section. Use these 3 mm-thick gel sections for the subsequent iterative gel-casting round.

Note: In the original gel recipe (single monomer solution incubation), gels are sometimes not completely permeated with polymer throughout, resulting in variable structural soundness. Approximately 1 to 2 gels out of a 6-well batch per plate come out suboptimal. Use the improved protocol (double incubation) for maximum reproducibility. The composition of the monomer solution is unchanged between protocols

### Supplementary Note 3: GFP 3D Reconstruction

Raw fluorescence images acquired from One-step Nanometer-scale Expansion (ONE) microscopy [5] are processed to isolate individual instances before reconstruction. For segmentation, we develop an unsupervised framework based on graph neural networks. As shown in **Supplementary Figure 24**, each image is divided into transformer derived [59] patches, that serve as local descriptors of the underlying signal. As shown in **Supplementary Figure 25**, the resulting feature vectors are embedded into a weighted graph whose edges encode patch similarity. Cluster assignments are obtained by minimizing a correlation clustering objective, L_CC_(S)=−∑_i,j_W_ij_∑_c_S_ic_S_jc_, where S_ic_ ∈ 0,1 denotes the assignment of node i to cluster c. As shown in **Supplementary Figure 26**, foreground and background separation is achieved in a two-stage process, ensuring robust isolation of protein complexes. Each segmented particle is extracted as a fixed-size sub-image I_0._ Based on connected component analysis and physical constraint-based filtering, we are performing partitions around mediods [60] to segregate the similar identity objects from the foreground mixed population. As shown in **Supplementary Figure 27**, this formulation encourages patches with similar appearance to co-cluster, yielding segmentation masks that isolate individual protein complexes for downstream analysis.

To enhance the structural signal of each isolated complex, we apply a physics-informed deconvolution [61] procedure in polar coordinates. Given an observed blurred image I_0_(r,θ), the estimate I_k_(r,θ) at iteration k, is iteratively refined according to I_k+1_(r,θ)=I_k_(r,θ).I_0_(r,θ)/{(I_k_∗R)(r,θ)}∗R^flip^(r,θ)], where R^flip^(r,θ) is the flipped PSF kernel, * is the convolution in polar coordinates. This formulation improves recovery of fine structural details compared to Gaussian PSF-based Lucy-Richardson deconvolution.

After normalization, the resized image is then processed by an encoder network Ψ_θ_(X) as shown in **Supplementary Figure 28**, which extracts a compact latent representation of the underlying molecular state. This encoder maps the normalized image to a low-dimensional latent variable: Ψ_θ_(X)=(R,t,s,z), where R ∈ SO(3) is a rotation matrix defining particle orientation, t ∈ R^2^ is a translation vector, s > 0 is a scaling factor accounting for gel expansion variability and z ∈ R^d^ encodes molecular conformation.

As shown in **Supplementary Figure 28**, the decoder V_ɛ_ acts as a continuous implicit representation of the 3D density in Fourier space, mapping frequency coordinates and conformation codes to Fourier coefficients: V_ɛ_(k,z)→X′(k;z), X′(k;z)∈R, where k=(k_x_,k_y_,k_z_)^T^ is a 3D frequency coordinate and z is the latent conformation vector. Thus, V_ɛ_ defines the Fourier space representation of the protein complex.

To generate a 2D projection, the decoder is evaluated slice-by-slice on a grid of frequency coordinates corresponding to the central slice through the 3D Fourier volume, rotated according to the estimated pose: k^′^_grid_=s^−1^⋅Rk_grid_, P^’^_c_=V_ɛ_(k^’^_grid_,z), P^’^(u)=P^’^_c_(u)⋅e^−2πi(uxtx+uyty)^. Here, k_grid_ are the 2D frequency coordinates in the projection plane, k′_grid_ are these coordinates rotated into the volume’s frame, P^’^_c_ is the centered Fourier slice decoded from the density field, and P^’^(u) is the final Fourier slice after applying the translation phase shift.

Model optimization is performed by comparing this predicted Fourier slice to the Fourier transform of the target image. The target image is transformed to the Fourier domain, Y^’^=F[X]. The symmetric mean squared error loss is minimized through the equation: L (P^’^, Y^’^) =min (∥P^’^−Y^’^∥^2^_2_, ∥P^’^−F[flip(X)]∥^2^_2_). The overall training objective is min_θ,ɛ_E_X∼data_[L(P^’^(X;Ψ_θ_,V_ɛ_);Y^’^)], where θ are the encoder parameters and ɛ are the decoder parameters.

Following training, the generator is evaluated slice-by-slice across a full grid of frequency coordinates to build a complete 3D Fourier volume. An inverse Fourier transform then produces a volumetric density map that represents a physically plausible 3D molecular structure obtained without imposing predefined atomic coordinates or using external templates or supervision. The resulting volumetric data is saved in MRC format, and final reconstructions were visualized and analyzed using UCSF ChimeraX [62].

